# Nodal signaling establishes a competency window for stochastic cell fate switching

**DOI:** 10.1101/2022.04.22.489156

**Authors:** Andrew D. Economou, Luca Guglielmi, Philip East, Caroline S. Hill

## Abstract

Specification of the germ layers by Nodal signaling has long been regarded as an archetype of how graded morphogens induce different cell fates. However, this deterministic model cannot explain why only a subset of cells at the margin of the early zebrafish embryo adopt the endodermal fate, while their immediate neighbours, experiencing similar signaling profiles, become mesoderm. Combining pharmacology, quantitative imaging and single cell transcriptomics, we demonstrate that sustained Nodal signaling establishes a bipotential progenitor state where cells initially fated to become mesoderm can switch to an endodermal fate. Switching is a random event, the likelihood of which is modulated by Fgf signaling. This inherently imprecise mechanism nevertheless leads to robust endoderm formation because of buffering at later stages. Thus, in contrast to previous deterministic models of morphogen action, Nodal establishes a temporal window when cells are competent to undergo a stochastic cell fate switch, rather than determining fate itself.

## Introduction

During embryonic development, pluripotent cells are progressively guided to a variety of cell fates by a series of decision-making processes involving chemical and mechanical signals. Crucially, these different cell types must be specified at the correct time and location. One of the most influential principles in developmental biology for how this happens has been the patterning of embryos by morphogen gradients. In its simplest form, a morphogen is produced at a localized source and diffuses across a tissue to form a gradient, with cells in different positions being exposed to different levels of the morphogen and as a consequence adopting different fates. Since it was originally proposed by Lewis Wolpert as a possible solution to the French Flag problem (Wolpert, 1969, 2011), numerous examples of possible morphogens have been identified across a diversity of developmental systems. These studies have resulted in several variants on the mode of morphogen gradient function, such as temporal signal integration, and the importance of downstream transcriptional circuits in morphogen interpretation (Briscoe and Small, 2015; Economou and Hill, 2020; Huang and Saunders, 2020; Pages and Kerridge, 2000). However, the importance of the amount of morphogen exposure in ultimately determining cell fate has remained a major theme in developmental biology.

One of the best-known examples of a morphogen in vertebrate development is the role of the transforming growth factor β (TGF-β) family member Nodal in mesoderm and endoderm specification in early zebrafish development (Hill, 2018; Rogers and Muller, 2019; Rogers and Schier, 2011). Two Nodal ligands Nodal-related 1 and 2 (Ndr1/2) are expressed in the early zebrafish embryo, where they signal through a serine/threonine kinase receptor complex, comprising two type I receptors and toe type II receptors, along with the co-receptor Tdgf1 (also called One-eyed pinhead (Oep)) (Hill, 2018). On ligand binding, the activated receptors phosphorylate receptor-regulated Smads (Smad2 in the early zebrafish embryo), allowing them to complex with Smad4 and accumulate in the nucleus. Together with additional transcription factors such as Foxh1 and Mixl1, these Smad complexes bind to enhancers and initiate a new program of gene expression. Nodal signaling plays a key role in the specification of the germ layers, as mutants affecting the ligands, receptors, or Smad2 show a characteristic phenotype which lacks many mesodermal and all endodermal derivatives, which can also be phenocopied with small molecule inhibitors of the type I receptor kinase activity (Dubrulle et al., 2015; Feldman et al., 1998; Gritsman et al., 1999; Sun et al., 2006).

In the early zebrafish embryo, Nodal ligands have been proposed to act via a morphogen gradient to induce the endodermal and mesodermal cell fates (Schier, 2003). Endodermal progenitors are initially marked by the expression of *sox32,* a gene encoding a transcription factor that is essential for induction of the endodermal lineage. Embryos null for *sox32* (the *casanova* mutant) make no endoderm (Kikuchi et al., 2001). The endodermal progenitors are found in the cell tiers closest to the margin of the embryo, while well-established mesodermal markers, such as *tbxta* are expressed up to 10 cell tiers from the margin of the embryo (Chen and Schier, 2002; Economou and Hill, 2020; Ober et al., 2003; Warga and Nusslein-Volhard, 1999). As the ligands Ndr1 and 2 are initially expressed at the margin of the zebrafish embryo in an extraembryonic tissue called the yolk syncytial layer (YSL) during early epiboly stages, it was proposed that they diffuse to form a gradient across the marginal cell tiers of the embryo (Chen and Schier, 2002; Meinhardt and Gierer, 2000; Saijoh et al., 2000). In the cells closest to the source (the YSL) where ligand levels are assumed to be highest, endoderm is induced, with lower signaling levels more distant from the YSL, inducing mesoderm.

However, recent work has suggested that Nodal does not function as a classical morphogen. Direct visualization of the range of Nodal activity revealed that it only extends about five cell tiers and therefore is not able to account for the expression of mesodermal target genes up to 10 cell tiers from the YSL (van Boxtel et al., 2015). Indeed, it was demonstrated that these long-range mesodermal targets are in fact induced by Fgf signaling, through the Ras-Raf-Mek1/2-Erk1/2 branch of the pathway downstream of the receptors, with the Fgf ligands themselves targets of Nodal signaling. Moreover, Fgf signaling through phosphorylated Erk (P-Erk) was also demonstrated to inhibit endoderm induction (van Boxtel et al., 2018), as inhibition of Fgf/P-Erk signaling led to increased numbers of endodermal progenitors. We also showed that the endoderm fate is restricted to the first two cell tiers by the local suppression of Erk signaling through the dual specificity phosphatase Dusp4, which is itself also a direct Nodal target gene (van Boxtel et al., 2018). The Nodal gradient is temporal as well as spatial, with the cells closest to the YSL that show the highest levels of signaling (read out by levels of phosphorylated Smad2 (P-Smad2)), being those induced for the longest duration (Dubrulle et al., 2015; van Boxtel et al., 2018). Moreover, sustained Nodal signaling is important for endoderm progenitor specification, as induction of *sox32* by Nodal requires a cascade of expression of other Nodal-induced transcription factors, specifically, Tbxta, Tbx16, Mixl1 and Gata5 (Kikuchi et al., 2000; Nelson et al., 2017; Ober et al., 2003; Reiter et al., 2001).

One particularly striking feature of endoderm induction in the zebrafish is that while the domains of Nodal and Fgf signaling extend all around the embryonic margin, only a subset of the marginal cells are induced to the endodermal lineage. Lineage tracing studies show that the most marginal cells are a mixture of endodermal and mesodermal progenitors, with only a subset of the cells expressing *sox32* (Kikuchi et al., 2001; Warga and Nusslein-Volhard, 1999). Cell tracking has shown that the position of the cells relative to the margin is relatively constant at these early stages (Dubrulle et al., 2015) and thus these most marginal cells are presumably all exposed to relatively high levels of Nodal signaling and suppressed Fgf signaling. Why only a subset of the cells exposed to the same inductive signals are induced to the endodermal lineage remains a conundrum. There is no evidence for a pre-pattern within the first two cell tiers, as a similar ‘salt and pepper’ pattern of *sox32* expression is observed in the animal pole when a Nodal-expressing clone was introduced at the 128 cell stage (van Boxtel et al., 2018). Furthermore, the spacing these *sox32*-positive endoderm progenitors does not appear to be the result of other signaling pathways, for example lateral inhibition through Notch signaling (Kikuchi et al., 2004; Ober et al., 2003).

Here, we investigate how neighbouring cells located in the same region of the embryo, and therefore exposed to the same signaling environment, are induced to different lineages. Our data do not support the consensus view of morphogen gradients, that cell fates are specified as a result of cells reading out different levels of signaling. Instead, we demonstrate that the distribution of endodermal progenitors is not the result of a highly regulated process but of a stochastic one. In this model, Nodal signaling provides a competency window for the stochastic switching of bipotential progenitors to an endodermal fate, with lower levels of Erk signaling favoring the switching process. Cells that do not switch to the endodermal fate, differentiate to mesoderm. This raises the question of how such probabilistic switching can accurately regulate the number of progenitors induced. We demonstrate that zebrafish embryos are robust to significant variation in the numbers of early endodermal progenitors, as their numbers are corrected during and after gastrulation to produce viable embryos with the appropriate amount of endoderm.

## Results

### Dynamics of mesoderm and endoderm induction and their separation

In many species, a common progenitor state of mesendoderm has been suggested to give rise to both mesoderm and endoderm (Nowotschin et al., 2019). In morphogen gradient models, high Nodal signaling is thought to induce endoderm while lower levels of Nodal signaling lead to mesoderm (Schier, 2009; Zorn and Wells, 2009). However, as outlined above, in the zebrafish embryo, mesoderm and endoderm progenitors arise in a “salt and pepper” pattern in the first two cell tiers from the embryonic margin in cells that are apparently exposed to similar levels of both Nodal and Fgf signaling. To begin to investigate the underlying mechanism we first set out to establish the dynamics of mesoderm induction and its specification relative to the endoderm.

We collected embryos throughout early development (at hourly intervals from 4 h post fertilization (hpf) to 8 hpf) and performed RNAscope in situ hybridization for *sox32,* multiplexed with two well established mesodermal markers: *tbxta* and *tbx16* (Griffin et al., 1998; Schulte-Merker et al., 1994) (Figure 1A, B). We developed an in toto quantitative imaging pipeline, whereby we segmented all the nuclei in each embryo and quantified the staining intensity for the three markers (Figure 1C, D). Tracking the relative expression of *tbxta* and *tbx16* revealed that by 6 hpf both markers are expressed around the margin before separating at gastrulation when *tbxta* is expressed alone in an internalized dorsal domain (the future axial mesoderm), with *tbx16* internalized in ventrolateral cells (part of the future paraxial mesoderm) (Figure 1A– C; Figure S1). A domain of non-internalized cells coexpressing the two mesodermal markers is also maintained in the cells closest to the margin of the embryo (Figure 1B; Figure S1). These results suggest that the most marginal cells – where endodermal progenitors are known to be induced over time (Kikuchi et al., 2001) – already express mesodermal markers (Figure 1A, B).

**Figure 1.**
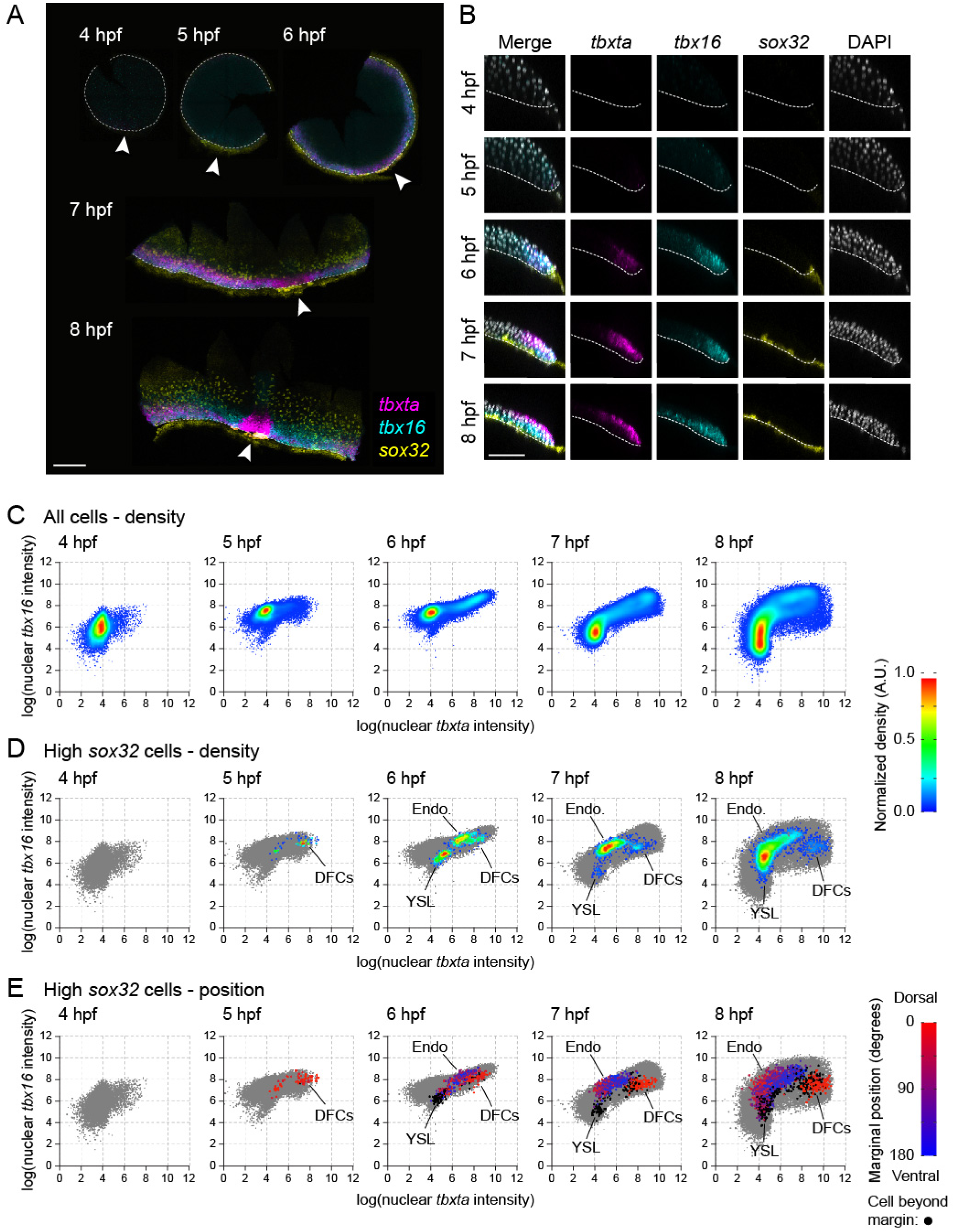
*sox32*-expressing endoderm progenitors emerge from a population of *tbxta/tbx16*-positive progenitors. (A) Time series showing maximum projections of RNAscope in situ hybridization for *tbxta*, *tbx16* and *sox32*, in embryos flat-mounted after staining. Dashed white line, embryo margin; arrowhead, dorsal. Scale bar, 250 µm. (B) Z-reconstruction from a 12.5-µm thick optical slice through the lateral regions of embryos in (A). Dashed line marks boundary between embryo and YSL. Colors as in (A); nuclei marked with DAPI (gray). Scale bar, 100 µm. (C) Density scatter plots showing nuclear fluorescence intensity staining for *tbxta* and *tbx16* for all cells pooled from four embryos for each time point. (D) Density scatter plots as in (C) showing cells with elevated *sox32* expression (defined as log(nuclear sox32 intensity) > 8.5) in the context of all cells (gray). The density scale in (C) and (D) is normalized to a maximum density of 1.0 for each plot. (E) Scatter plots from (D) colored to show the position of the closest point at the margin of the embryo from dorsal (0 degrees – red) to ventral (180 degrees – blue). Lateral regions on both sides of each embryo are given the same positional value. Nuclei beyond the margin are marked in black. Populations of cells in (D) and (E) composed predominantly of black cells are marked as YSL cells, while populations composed predominantly of dorsal (red) cells are marked as dorsal forerunner cells (DFCs). The remaining cells are endodermal progenitors (Endo). See also Figure S1.

Do endodermal progenitors themselves express markers of mesodermal progenitors? Plotting the levels of *tbxta* and *tbx16* for only the cells with the highest levels of *sox32* expression confirmed that endodermal progenitors do not begin to appear until after 5 hpf (at this time the only cells with elevated *sox32* were the YSL and the dorsal forerunner cells (DFCs), which are the precursors of Kupffer’s vesicle (Kikuchi et al., 2001)) (Figure 1D). At 6 hpf, endodermal progenitors are found among the most marginal cells, with elevated *tbxta* and *tbx16* (Figure 1D). These cells can then be seen to internalize and migrate away from the margin (Figure 1B; Figure S1). Of note, *tbxta* levels decrease in the majority of endodermal progenitors from 7 hpf, when they have begun to ingress (although *tbxta* levels remain high in the DFCs) (Figure 1D, E). In contrast, there is a broad range of *tbx16* expression levels among the endodermal progenitors (Figure 1D, E). Therefore, endodermal progenitors are induced among the population of marginal cells expressing mesodermal markers.

### Endoderm progenitors appear randomly without spatial or temporal bias

Having established broadly when endodermal progenitors are induced relative to early mesodermal markers, we set out to determine precisely where endodermal progenitors are located – both relative to their position around the margin and to each other – and how this pattern is generated. Using a dataset of densely sampled embryos across early epiboly (fish were spawned continuously for one hour, then embryos sampled every 15 min from 4.25 to 5.25 hpf) we used our quantitative imaging pipeline to identify endodermal progenitors as cells with the highest levels of nuclear staining for *sox32* using RNAscope in situ hybridization and mapped their position around the margin of the embryo, excluding nuclei corresponding to the YSL (Figure 2A–D; Figure S2A). The dorsal enrichment of *sox32*-positive cells corresponds to DFCs, which were also removed from endodermal cell counts.

**Figure 2.**
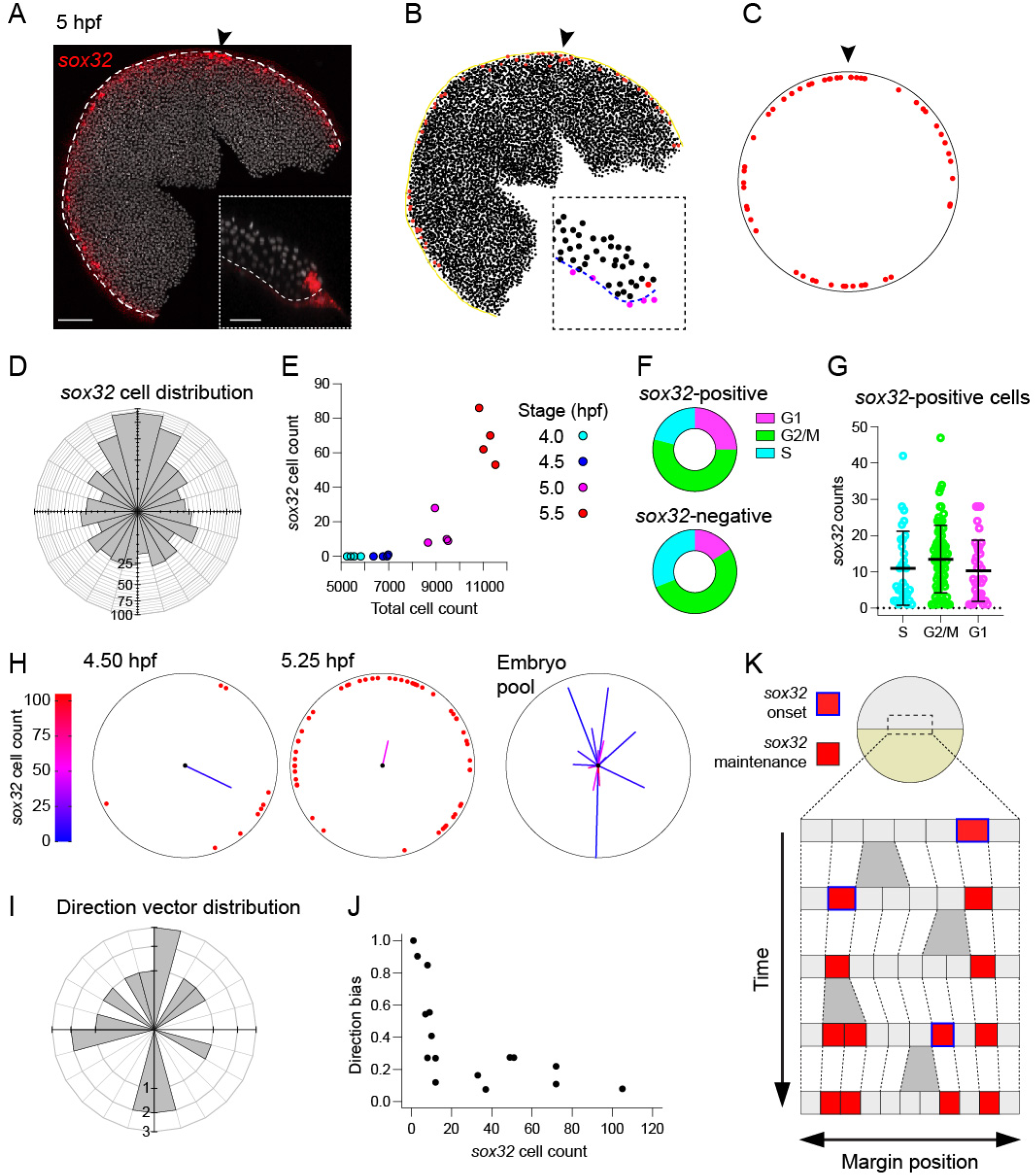
*sox32*-positive cells are induced randomly within the first two cell tiers. (A) Maximum projection of RNAscope in situ hybridization for *sox32*, in 5 hpf flat-mounted embryo. Nuclei marked with DAPI (gray). Dashed white line, embryo margin; arrowhead, dorsal. Scale bar, 125 µm. Inset: Z-reconstruction from a 12.5-µm thick optical slice through the lateral regions of embryo as in Figure 1B. Scale bar, 50 µm. (B) Digitization of embryo in (A) showing XY positions of centroids from segmentation of all nuclei in embryo. *sox32*-positive cells are marked in red, with those corresponding to the YSL removed. Yellow line, embryo margin; arrowhead, dorsal. Inset: Positions of all nuclear centroids from inset in (A). *sox32*-positive cells are red, with those that are in the YSL or beyond the margin, and thus eliminated from the analysis, in magenta. Blue dashed line, embryo margin. (C) Reconstruction of embryo in (B) showing the position of endodermal progenitors (red). Arrowhead, dorsal. (D) Distribution of 827 *sox32*-positive cells around the embryonic margin for 20 embryos collected continuously from 4.25–5.25 hpf after a mass spawning showing a significant dorsal enrichment which correspond to DFCs (Watson’s Test for Circular Uniformity: test statistic = 3.39, p-value < 0.01). Dorsal, up. (E) Relationship between number of *sox32*-positive endodermal progenitors (YSL and DFCs removed) and total cell number for embryos collected at 30 min intervals from a single clutch. Colors indicate embryonic stage. (F) Donut graphs showing the percentage of cells in G1, G2/M or S phase for *sox32*-positive cells (left) and *sox32*-negative cells (right) at the margin. (G) Scatterplot showing *sox32* expression levels in *sox32*-positive cells falling in the different phases of the cell cycle. Means ± SD are shown. Dotted line shows zero. Statistical difference was tested using a Kruskal-Wallis test with Dunn’s multiple comparison correction. n.s for all comparisons. (H) Two representative embryos showing the distribution of *sox32*-positive endodermal progenitors around the margin. Average direction vector shown on each embryo, where length indicates direction bias (vector touching circle indicated a direction bias of 1.0), colored by the total number of *sox32*-positive cells. Embryo pool shows vectors from the 20 embryos in (D) (only 16 vectors are shown as the four youngest embryos have no progenitors). Dorsal, up. (I)) Distribution of average direction vectors for the 16 embryos in (H), grouped into 15-degree bins. Dorsal, up. (J) Plot of direction bias against total number of *sox32*-positive endodermal progenitors for the 16 embryos in (H). (K) Schematic showing proposed model for the appearance of *sox32*-positive endodermal progenitors. A series of cells at the margin of the embryo proliferate through time (gray bifurcations). In a random manner, cells turn on *sox32* (blue outlined cells), and expression is maintained (red cells). See also Figure S2.

We then asked when the endoderm progenitors first appear. Collecting embryos at 30-min intervals from a single synchronized clutch from 4 hpf to 5.5 hpf, corresponding to early epiboly stages, we found that rather than all appearing at once, the number of endodermal progenitors increased steadily through time (Figure 2E). We noticed the same overall trend in the number of *sox32*-positive endodermal progenitors in a mass-spawned embryo dataset, although in this case the data were noisier as they came from many different clutches (Figure S2B). At these embryonic stages the cell cycle duration is 30–45 min (Siefert et al., 2015), meaning that most cells only divide once in the 1.5 h window we are studying. We therefore reasoned that the increase in endodermal progenitors through time could either be the result of progenitors being continuously induced throughout early epiboly, or a small number of progenitors induced during a short, early time window, before proliferating at an increased rate relative to their neighbors.

To distinguish between these two scenarios, we investigated expression of cell cycle markers at the margin. We analyzed a published scRNA-seq dataset for 50% epiboly zebrafish embryos (Farrell et al., 2018) and extracted cells positive for *gata5*, which is expressed in the first two cell tiers from the margin (Figure S2C, D; Supplementary File 1). Notably, cells within this pool did not display a particular bias towards a specific phase of the cell cycle (Figure S2E). Confirming this, the proportion of cells expressing G2/M markers in cells negative or positive for *sox32* was also equivalent (Figure 2F). Moreover, compared to classic G2/M markers, expression levels for *sox32* were uniform irrespective of the phase of the cell cycle (Figure 2G, Figure S2F). We thus excluded an increase in proliferation in endodermal progenitors as a mechanism to explain the steady increase in progenitor number over time. Instead, we conclude that they are continuously induced throughout early epiboly.

We then focused on how the endoderm progenitors are induced spatially. Pooling all endodermal progenitors from our densely sampled mass-spawned dataset where embryos were collected throughout early epiboly, we could not distinguish the distribution of progenitors around the margin from a uniform distribution, once the DFCs had been excluded (Figure S2G, H). Thus, cells were equally likely to be found at any position around the margin, suggesting no bias, for example around the dorsal–ventral axis.

To further explore whether there was any regular pattern in how progenitors were positioned relative to each other we generated an average progenitor direction vector for each embryo by summing the direction vectors for all progenitors in an embryo (Figure S2I). Again, there was no directional bias for the orientation between embryos, with the distribution of directions across all embryos indistinguishable from uniform (Figure 2H, I). For each embryo, the directional bias in the positions of progenitors was reflected by the magnitude of the direction vector (a large value would reflect all cells being localized in a similar region). We noticed that for embryos with very few progenitors, some had large magnitudes, with others small, and magnitudes decreased as the numbers of progenitors increased (Figure 2J). We found that all of these features could be recapitulated using an unbiased simulation where, during early epiboly, any marginal cell could turn on *sox32* with a very low probability, and once a cell expressed *sox32*, expression is maintained (Figure 2K; Figure S2J–P). The concordance between the simulation and our data indicates that there is no spatial or temporal bias in the induction of endodermal progenitors. Rather, marginal cells otherwise fated to become mesoderm turn on and maintain *sox32* expression in a random manner.

### Nodal and Fgf signaling levels affect the likelihood of endoderm induction, rather than determining cell fate itself

We next determined what role signaling plays in determining this spatially random pattern of *sox32* expression. Differences in morphogen levels have generally been considered at the level of broad graded profiles across fields of cells. Indeed, when we and others previously constructed Nodal and Fgf signaling profiles in early zebrafish embryos, we averaged the intensity of immunofluorescence for the signaling readouts P-Smad2 and P-Erk respectively across the multiple cells within each cell tier relative to the margin (Lord et al., 2021; Rogers et al., 2017; van Boxtel et al., 2018). However, it is possible that the heterogeneity seen in the appearance of endodermal progenitor cells could be the result of cell-to-cell differences in the levels of Nodal or Fgf signaling. We therefore asked whether the unpredictable nature of *sox32* expression could be explained by noise in cell signaling.

We performed a double immunostaining for P-Smad2 and P-Erk (as readouts for Nodal and Fgf signaling through Erk respectively) simultaneously with an RNAscope in situ hybridization for *sox32* (Figure 3A, B). By quantifying nuclear levels of *sox32* alongside P-Smad2 and P-Erk for all cells in the embryos, we were able to profile all *sox32*-positive and - negative cells for their Nodal and Fgf signaling states (Figure 3C). To understand whether the distribution of *sox32*-positive cells could be explained by cell-to-cell variation in the levels of Nodal signaling, we first asked whether the increase in the number of *sox32*-positive cells over time (see Figure 2E) was associated with an overall increase in Nodal signaling levels. Comparing the levels of P-Smad2 across the first two cell tiers (where endodermal progenitors are induced) throughout the early epiboly stages (4 hpf–5.5 hpf) showed that the initial increase in *sox32* at 5 hpf was associated with an increase in mean P-Smad2 levels, but the continued increase in *sox32*-positive cell numbers after this stage occurred without any further increase in overall P-Smad2 levels (Figure 3D, Figure S3A).

**Figure 3.**
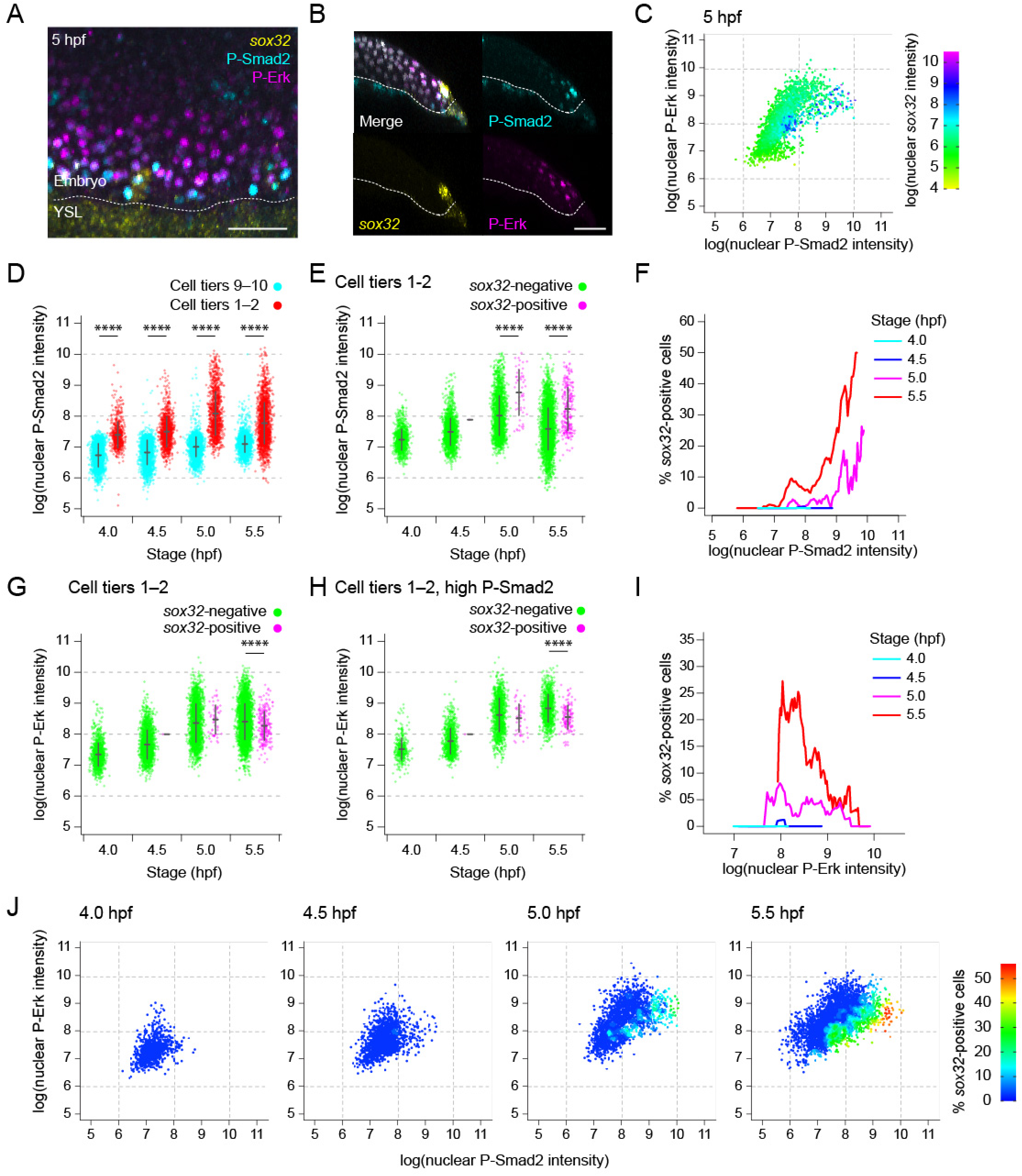
Levels of Nodal and Fgf signaling are not deterministic for endoderm progenitor induction. (A) Maximum projection through the margin of a 5 hpf embryo showing immunofluorescence for P-Smad2 and P-Erk, with RNAscope for *sox32*. Dashed white line, embryo margin. Scale bar, 50 µm. (B) Z-reconstruction from a 12.5-µm thick optical slice through the lateral regions of same embryo as in (A) showing *sox32* expression in endodermal progenitors relative to P-Smad2 and P-Erk staining. Dashed white line, embryo margin. Scale bar, 50 µm. (C) Scatter plot showing nuclear staining intensity of P-Smad2 and P-Erk for a single 5-hpf embryo. Cells colored by nuclear staining intensity for *sox32*. (D) Plot showing P-Smad2 staining intensity for all cells in the first and second cell tiers compared to background levels (ninth and tenth cell tiers). Means ± SD are shown. Statistical difference was tested using a t-test. For all timepoints: p < 0.0001. (E) Plot showing P-Smad2 staining intensity for all cells in the first and second cell tiers, broken down into *sox32*-positive and -negative cells. Means ± SD are shown. Statistical difference was tested using a t-test. For both 5.0 and 5.5 hpf: p < 0.0001. (F) Traces showing the proportion of cells that are *sox32*-positive for a given P-Smad2 staining intensity for cells in the first and second cell tiers at different stages. The proportion of *sox32*-positive cells for a given level of P-Smad2 staining is based on all cells within a window of ± 5% of the total range of P-Smad2 staining intensities for that stage. (G) Plot showing P-Erk staining intensity for all cells as in (E). Means ± SD are shown. Statistical difference was tested using a t-test. 5.0 hpf: n.s, 5.5 hpf: p < 0.0001. (H) Plot showing P-Erk staining intensity for all cells in the first and second cell tiers with P-Smad2 staining elevated above background levels. Means ± SD are shown. Statistical difference was tested using a t-test. 5.0 hpf: n.s, 5.5 hpf: p < 0.0001. (I) Traces showing the proportion of cells that are *sox32*-positive for a given P-Erk staining intensity for cells in the first and second cell tiers with elevated P-Smad2 staining (defined as in (H)) at different stages. Traces are as in (F). In (H, I) the 99^th^ percentile of P-Smad2 staining intensity for the cells from the ninth and tenth cell tiers is used as a threshold to define elevated P-Smad2. (J) Scatter plot showing nuclear staining intensity of P-Smad2 and P-Erk for cells in the first two cell tiers through time. Color coding indicates the proportion of cells for the given signaling profile which are *sox32*-positive. The data are a re-analysis of the data presented in panels F and I. Analyses in this figure are based on the same dataset as Figure 2E, comprising four embryos for each time point. See also Figure S3.

Moreover, while levels of P-Smad2 were elevated in the first two cell tiers relative to background, we were struck by how great the range of P-Smad2 levels was among these most marginal cells, with many cells having P-Smad2 levels comparable to background (i.e P-Smad2 levels in cell tiers 9–10 (van Boxtel et al., 2018)). This was not due to the antibody staining as it is also seen using subcellular localization of GFP-Smad2 as a Nodal signaling readout (Dubrulle et al., 2015), and was also independent of cell depth in the tissue (data not shown). Therefore, it is likely the result of heterogeneity in the responsiveness of cells to Nodal. We tested whether variation in the levels of P-Smad2 within the first two cell tiers could explain which cells become endodermal progenitors. Comparing P-Smad2 levels between *sox32*-positive and -negative cells in the first two cell tiers at 5.0 and 5.5 hpf showed that the mean P-Smad2 levels were indeed significantly higher in *sox32*-positive cells compared to negative cells (Figure 3E; see also Figure S3B) However, there was still considerable overlap in P-Smad2 levels between the two populations (Figure 3E; Figure 3SB).

To better explore the relationship between P-Smad2 and *sox32* expression we calculated the proportion of cells at different stages that were *sox32*-positive, for different levels of P-Smad2. We see evidence for a temporal effect of P-Smad2 as no *sox32*-positive cells appear before 5 hpf, despite moderate levels of P-Smad2 (Figure 3F). Then at both 5.0 and 5.5 hpf, the proportion of cells that were *sox32*-positive increased with the level of nuclear P-Smad2 (Figure 3F; Figure S3C). Strikingly, however, for any given level of nuclear P-Smad2, the number of cells that were *sox32*-positive also increased with time. Therefore, while variation in the level of P-Smad2 between cells is predictive of *sox32* expression, it alone is not sufficient to determine whether marginal cells become endodermal progenitors. A given level of P-Smad2 does not determine how many cells will be *sox32*-positive, as the progenitors accumulate through time, it just makes their appearance more likely.

Given that we and others have previously shown that Fgf signaling is inhibitory to the endodermal fate (Mizoguchi et al., 2006; Poulain et al., 2006; van Boxtel et al., 2018), we reasoned that cell-to-cell variation in levels of P-Erk within the first two cell tiers might determine whether or not a cell experiencing a given level of nuclear P-Smad2 would express *sox32*. We therefore compared P-Erk levels across cells in the first two cell tiers stratified by *sox32* expression (Figure 3G). At 5.0 and 5.5 hpf, there was substantial overlap in the levels of P-Erk between the two populations, suggesting variation in Fgf signaling between cells could not explain which cells were induced to become endoderm progenitors. As many of these cells would have low P-Smad2 levels, and are therefore not be expected to express *sox32*, we repeated the analysis, but restricted it only to cells with elevated P-Smad2. Again, we noted substantial overlap in the levels of P-Erk between *sox32*-positive versus-negative cells, but found that *sox32*-positive cells at 5.5 hpf had on average lower levels of P-Erk (Figure 3H). We therefore asked what proportion of the high P-Smad2 cells were *sox32*-positive for different levels of P-Erk. Again, at 5.5 hpf we noted that *sox32*-expressing cells were preferentially those with lower levels of P-Erk (Figure 3I). This suggested that while high levels of Nodal and low levels of Fgf signaling did not specifically define the endodermal progenitor population, the signaling levels dictated the likelihood of a cell expressing *sox32*.

To confirm this, we determined the percentage of cells in the first two cell tiers that expressed *sox32*, relative to their levels of P-Smad2 and P-Erk (Figure 3J). This analysis clearly showed that at 5.0 and 5.5 hpf, the percentage of *sox32*-positive cells increased with increasing Nodal and decreasing Fgf signaling, and that for a given signaling level, the proportion of *sox32*-positive cells increased with time.

### *sox32* expression, and hence endoderm progenitor induction, is regulated by a bistability

Our data show that *sox32* expression is not a simple readout of heterogeneous signaling inputs. Instead, we hypothesized that *sox32* expression could be bistable and Nodal signaling may push cells closer to a bifurcation point. Inherent noise in molecular cellular processes would mean that some cells would by chance cross this bifurcation point, resulting in the onset of *sox32* expression and irreversibly commit to the endodermal fate. As induction would be a random event in time (and space), *sox32*-positive cells would accumulate through time. In addition, if cells with lower P-Smad2 levels are further from the bifurcation point, fewer endodermal progenitors should accumulate at regions of lower Nodal signaling.

It is well established that Nodal signaling is required for endoderm induction (Kikuchi et al., 2001). If the system is bistable, commitment to this lineage should be maintained even if Nodal signaling is withdrawn. We therefore treated embryos with the Nodal signaling inhibitor SB-505124 (DaCosta Byfield et al., 2004) (see Figure S4A which shows the efficacy of the inhibitor) at 4.75 hpf, when endodermal induction has already begun, collected embryos at 5.25 hpf, and compared them with DMSO-treated controls. As a control we also treated embryos at 4.25 hpf, a time before any *sox32*-expressing cells were detectable (Figure 4A– C). If the Nodal receptor inhibitor was added early, no endodermal progenitors were induced (Figure 4C iv), but if it was added at 4.75 hpf, then endodermal progenitors were present, but in reduced numbers compared with control embryos of the same age treated with DMSO (Figure 4C - compare iii and v). To fully interpret this result it was important to demonstrate ongoing transcription in the absence of Nodal signaling. Because we could still see nuclear *sox32* transcripts in the absence of nuclear P-Smad2 staining (Figure S4A, B), this suggests that the maintenance of *sox32* in the absence of Nodal signaling is the result of active transcription rather than transcript stability. The instability of the *sox32* transcripts is also supported by the rapid clearance of *sox32* expression after 6 hpf (Figure S4C). Thus, we conclude that Nodal signaling is required for the induction of *sox32*, but not its maintenance.

**Figure 4.**
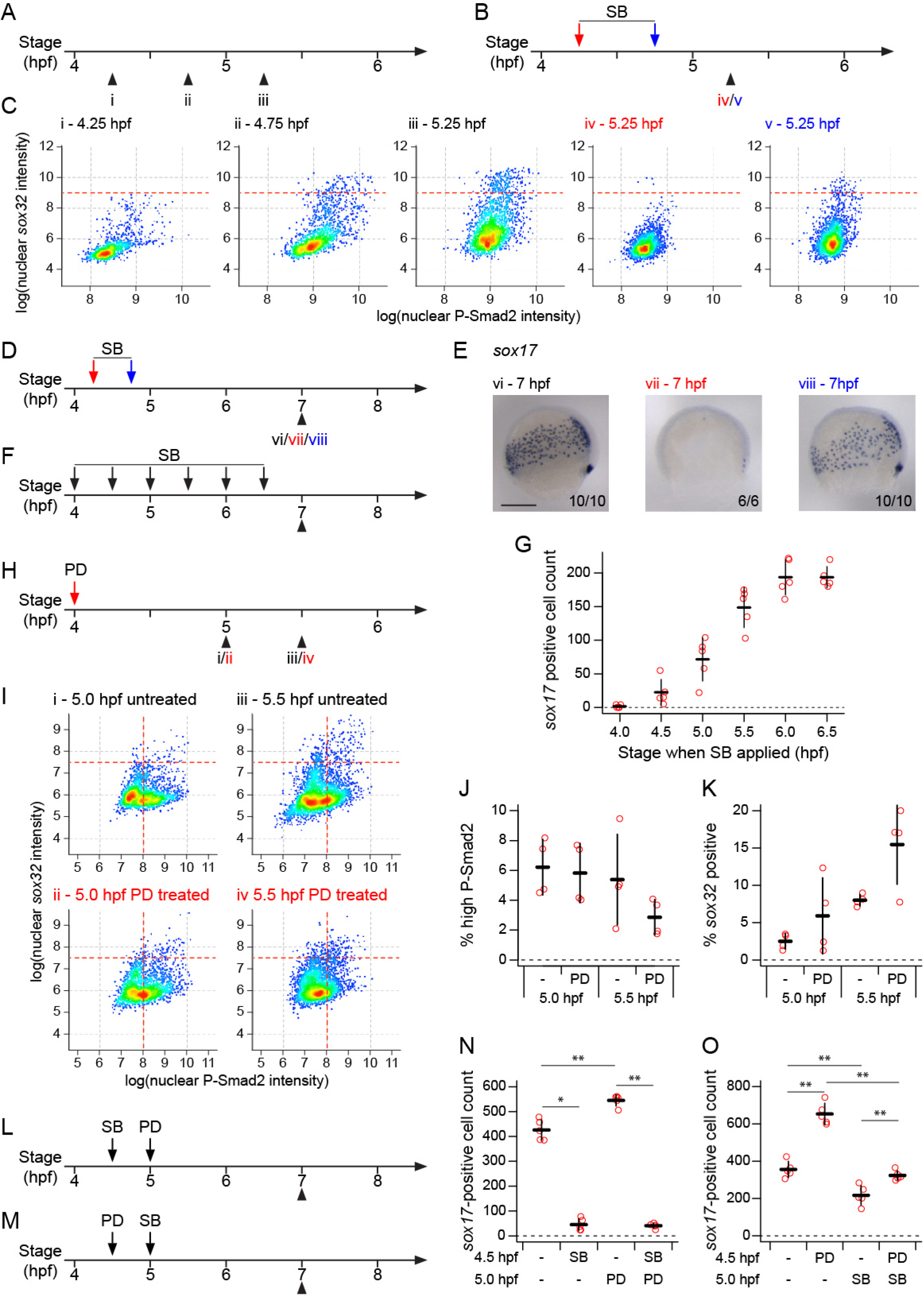
Endoderm progenitors require Nodal signaling for their induction, but not maintenance. (A–C) Schematics showing collection times of untreated embryos (A) and timings of SB-505124 (SB) application and subsequent embryo collection (B) for plots shown in (C). Colors of roman numerals in correspond to the treatment (red and blue correspond to arrows in (B), black to no SB). Density scatter plots showing nuclear fluorescence intensity staining for P-Smad2 and *sox32* for all cells in the first two cell tiers pooled from four embryos for each condition (C). Dashed red lines indicate threshold defining *sox32*-positive cells. (D, E) Schematic showing the timing of SB application and subsequent embryo collection for embryos in (E). Colors of roman numerals as in (A) and (B). Representative embryo following WMISH for *sox17* (E). Roman numerals correspond to treatments as defined in schematics in (D). Dorsal to right; animal pole, top. Scale bar, 250 µm. Numbers in the bottom right refer to the number of embryos showing the phenotype out of the total number studied. (F) Schematic (F) showing the timings of SB application and subsequent embryo collection for (G). (G) Quantification of *sox17*-positive cell numbers for embryos treated with SB as in (F). Means from five embryos per condition ± SD shown. (H) Schematic showing the timing of PD-0325901 (PD) application and subsequent embryo collection for (I). Roman numerals in red indicates PD treatment. (I) Density scatter plots showing nuclear fluorescence intensity staining for P-Smad2 and *sox32* for all cells in the first two cell tiers pooled from four embryos for each condition. Roman numerals correspond to treatments as defined in (H). Horizontal dashed red lines indicate threshold defining *sox32*-positive cells. (J) Plot showing the percentage of cells in (I) with high P-Smad2 (as defined by vertical dashed red lines in (I)) for each treatment condition. Means ± SD are shown. Statistical difference was tested using a Wilcoxon rank sum test. For both 5 and 5.5 hpf, n.s. (K) Plot showing the percentage of high P-Smad2 cells in (I) which are *sox32*-positive (as defined by horizontal dashed red lines in (I)) for each treatment condition. Means ± SD are shown. Statistical difference was tested using a Wilcoxon rank sum test. For both 5 and 5.5 hpf, n.s. (L, M) Schematics showing the timings of sequential treatment of embryos with SB and PD, and subsequent embryo collection for plots in (N,O). (N, O) Quantification of *sox17*-positive cells number for embryos treated with SB and PD as defined in schematics (L) and (M) respectively. Means from five embryos per condition ± SD are shown. Statistical difference was tested using a Wilcoxon rank sum test. For (N): p <0.05 (-/- vs SB/-), p < 0.01 (-/- vs -/PD), n.s (SB/- vs SB/PD), p < 0.01 (-/PD vs SB/PD). For (O): p < 0.01 for all comparisons shown. See also Figure S4.

At around 5.5 hpf once embryos have started to gastrulate, endodermal progenitors go on to express *sox17* and then migrate over the yolk towards the dorsal animal region of the embryo (Alexander and Stainier, 1999). To confirm that the maintenance of *sox32* expression in the absence of Nodal signaling resulted in cells maintaining their commitment to the endodermal lineage, we repeated the above experiment, but fixed embryos at 7 hpf and performed in situ hybridization for *sox17* (Figure 4D, E). As above, the cells continued to differentiate down the endodermal lineage even in the absence of Nodal signaling. Moreover, this model predicts that as cell switching to the endodermal lineage occurs continuously through early epiboly, the later Nodal signaling is inhibited, the more progenitors should accumulate. Repeating the above experiment but applying SB-505124 at successively later timepoints demonstrated that this is the case; the later Nodal signaling was inhibited, the more *sox17*-positive cells were present at 7 hpf (Figure 4F, G).

Although Fgf signaling is known to be inhibitory for endoderm induction, it is unclear at what level it inhibits the ability of cells to switch to the endodermal lineage. We reasoned that Fgf signaling could either work downstream of Nodal, by reducing the likelihood of cells switching fate when experiencing a particular level of Nodal signaling, or Fgf could act upstream of Nodal by inhibiting Nodal signaling, and thereby indirectly reducing the likelihood of cells switching fate. To test this, we treated embryos at 4 hpf with the Mek inhibitor PD-0325901 (Anastasaki et al., 2012) (Figure S4D), collected embryos at 5.0 and 5.5 hpf, and performed immunofluorescence for P-Smad2 with RNAscope for *sox32* (Figure 4H–K). Upon Mek inhibition, the proportion of these cells which were *sox32*-positive increased, while the proportion of cells with elevated P-Smad2 did not. Therefore, Mek inhibition increases the proportion of cells experiencing a given level of Nodal signaling being induced to endoderm, indicating that Fgf signaling acts by modulating the likelihood of cells exposed to Nodal signaling acquiring an endodermal fate.

A key prediction of this model is that inhibiting Fgf signaling should only have an effect on endoderm induction if Nodal signaling is active. Indeed, inhibiting Fgf signaling after Nodal signaling had been blocked led to the same number of progenitors as blocking Nodal signaling alone early in development, whilst blocking Fgf alone late in development led to a slight increase in progenitor numbers (Figure 4L, N). Conversely, the late inhibition of Nodal signaling after Fgf signaling inhibition still led to a reduction in *sox17*-positive cells compared to only blocking Fgf signaling early (Figure 4M, O). These observations support the model that Fgf signaling functions by modulating the effect of Nodal signaling.

### Switching to an endodermal fate is not initially associated with suppression of mesodermal markers

So far, our data indicate that cells in the first two cell tiers of the margin of the embryo are fated to differentiate to mesoderm, but during epiboly, some cells randomly switch to the endodermal fate. We next investigated whether this switch in fate also involved the suppression of their mesodermal character, and reciprocally whether we could find evidence for a unique mesodermal master regulator. To capture transcriptional changes underlying endoderm and mesoderm specification we used the same published scRNA-seq dataset (Farrell et al., 2018) as above for the cell cycle analysis to investigate gene expression at the embryo margin. To narrow down the analysis to the first few cell tiers, we focused on *gata5*- positive cells at 50% epiboly (5.3 hpf) which display mesodermal and endodermal marker expression and are devoid of more anterior ectodermal inputs (Figure S2B, C; Figure S5A, B). We performed differential gene expression analysis on cells positive or negative for *sox32* within the *gata5*-positive pool (Figure 5A, B; Supplementary File 2). Surprisingly, at 50% epiboly gene expression across the two populations was rather homogeneous. Indeed, beside *sox32*, genes highly represented in endodermal progenitors were also present in their mesodermal progenitor neighbours (Figure 5A). Furthermore, mesodermal markers were symmetrically expressed in endodermal progenitors (Figure 5B). Thus, there was no evidence for expression of a mesodermal master regulator, equivalent to Sox32 for the endoderm, and the mesodermal signature was not suppressed in the endoderm progenitors.

**Figure 5.**
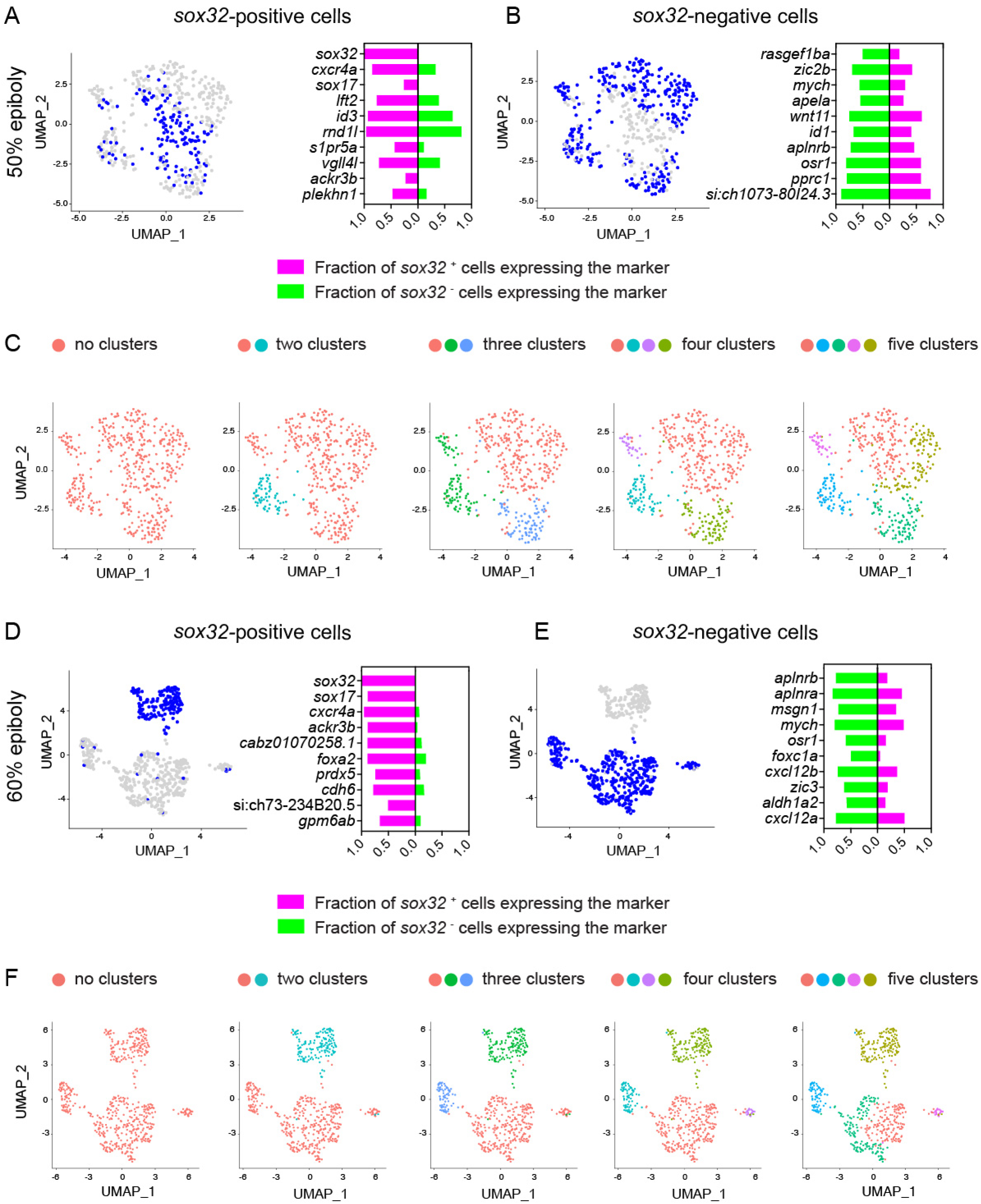
Switching to an endodermal fate is not associated with suppression of mesodermal fate. (A) Left: uniform manifold approximation and projection (UMAP) visualization of *gata5*-positive cells derived from 50% epiboly stage embryos. Blue cells are those expressing at least one read count for *sox32*. Right: stacked bar plot showing the pct (percentage of expressing cells) for the top 10 genes differentially expressed in *sox32*-positive cells (magenta) compared to the pct for the same genes in *sox32*-negative cells (green). (B) Left: as in (A) but blue cells are those expressing less than one read count for *sox32*. Right: stacked bar plot showing the pct for the top 10 genes differentially expressed in *sox32*-negative cells (green) compared to the pct for the same genes in *sox32*-positive cells (magenta). (C) UMAP visualization of *gata5*-positive cells at 50% epiboly. Cells were clustered with increasing granularity across five iterations, Find cluster resolution = 0.1 to 0.5. Color coding refers to the different clusters. (D) As for (A) but for 60% epiboly embryos. (E) As for (B), but for 60% epiboly embryos. (F) As for (C), but for 60% epiboly embryos, Find cluster resolution = 0.01 to 0.5. See also Figures S5 and S6.

As a more global approach to assess transcriptional heterogeneity at the margin, we clustered *gata5*-positive cells into different numbers of transcriptionally-defined populations and asked whether *sox32*-positive cells would define a distinct cluster (Figure 5C). Strikingly, *sox32*-positive cells distributed across multiple clusters and these cells displayed analogous levels of *sox32* counts, irrespective of their cluster allocation (Figure 5C; Figure S5E). Of note, these findings were confirmed by repeating this analysis on cells positive for *mixl1*, which is expressed in a domain of five cell tiers at the margin and defines a transcriptionally distinct cluster at 50% epiboly (Figure S2C, D; Figure S6A–C; Supplementary File 3). Together these data show that by 50% epiboly, a subset of progenitors at the margin switches on *sox32* within an otherwise transcriptionally homogeneous pool.

We have shown above that after initial induction of *sox32*, cells proceed towards the endodermal lineage and express *sox17*. We therefore asked whether *sox32*-positive cells would diverge from their mesodermal counterpart during gastrulation. We repeated the same transcriptional analysis at 60% epiboly which corresponds to mid gastrulation (Figure 5D–F, Figure S5C, D; Figure S6D, E; Supplementary File 4 and 5). We found that together with *sox32*, several endoderm-specific markers were now robustly and uniquely expressed in the endodermal progenitors (Figure 5D), and this was accompanied by a sharper suppression of mesodermal markers (Figure 5E). Consistent with this, at 60% epiboly, *sox32*-positive cells could be readily identified within a transcriptionally distinct cluster (Figure 5F; Figure S5F; Figure S6F), which uniquely expressed high levels of *sox32,* showing that by this time they have acquired a distinct identity. Therefore, the switching of cells to an endodermal fate from what would otherwise be a default mesodermal fate is initially dependent on just the onset of *sox32* expression, with the two cell fates only becoming transcriptionally distinct by mid gastrulation.

### The role of Nodal and Fgf signaling in mesoderm induction

We have shown that Nodal signaling is required for the induction, but not maintenance of endoderm progenitors, but it is not clear whether the same is true for mesoderm progenitors. We have previously shown that Fgf signaling downstream of Nodal is inhibitory for endoderm induction, but is required for the induction of mesoderm progenitors in the cell tiers further from the YSL (van Boxtel et al., 2018). To understand the relative importance of the timing of Nodal and Fgf signaling on mesoderm induction, we performed timed inhibition of Nodal and Fgf/P-Erk signaling prior to and up to mid gastrulation (4–7 hpf) (Figure 6A), and assessed the consequence of these treatments for the maintenance of paraxial mesoderm progenitors (marked by *tbx16* at 8 hpf) (Figure 6B–E), and their derivatives, such as somitic muscles in the trunk and tail, and jaw muscles in the head (marked by *myod* at 24 hpf) (Figure 6G–J).

**Figure 6.**
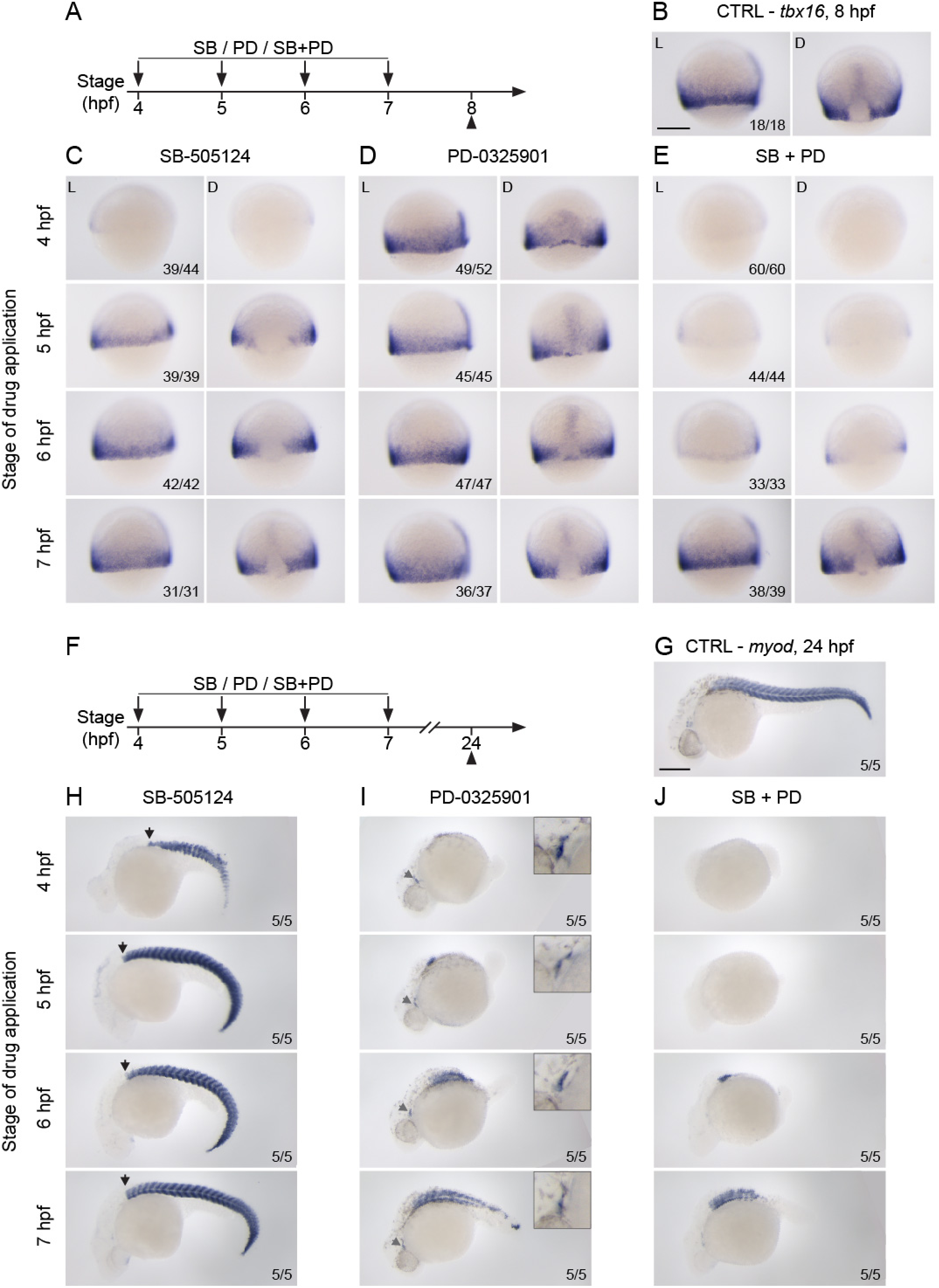
Nodal and Fgf signaling are required for mesoderm induction. (A–E) WISH showing lateral (L) and dorsal views (D) of 75% epiboly embryos stained for *tbx16*. Embryos underwent a time series of drug treatments with either 50 µM SB-505124 (left panel), 5 µM PD-0325901 (middle panel) or the combination of the two (right panel). Drug treatments were started at either 4 hpf (sphere stage), 5 hpf, 6 hpf or 7 hpf and embryos were fixed at 75% epiboly (8 hpf) (A). 75% epiboly control (CTRL) embryos are shown for comparison (B). Note that for embryos treated from sphere stage with SB-505124 or the combination of SB-505124 and PD-0325901 the dorsal side of the embryo cannot be unequivocally identified, as early Nodal inhibition prevents both *tbx16* expression and the formation of the dorsal shield. Therefore, for these samples, side-views of representative embryos are shown. Scale bar, 250 µm. (F–J) As for (A–E) but embryos were fixed at 24 hpf and stained for *myod*. 24 hpf control (CTRL) embryos shown for comparison (G). In this case, all views are lateral. In (H), the black arrow marks the most anterior position of the somites. In (I), the gray arrow indicates the jaw muscle (shown magnified, top right). Scale bar, 250 µm. Numbers in the bottom right of the images refer to the number of embryos showing the phenotype out of the total number studied.

Inhibition of Nodal signaling from 4 hpf suppressed the expression of *tbx16* at the margin (Figure 6C). This early effect was followed by the loss of both head and trunk mesoderm at 24 hpf (although tail mesoderm, which is known to only partially depend on Nodal signaling, was maintained) (Figure 6H) (Feldman et al., 1998; Gritsman et al., 1999). In contrast, *tbx16* expression was maintained if Nodal inhibition occurred from 5 hpf onwards (Figure 6C). Consistent with these results, expression of *myod* in the trunk was also restored under these conditions (Figure 6H). This suggested that after 5 hpf, mesoderm derivatives can still form independently of Nodal signaling. In contrast to Nodal inhibition, in the absence of Fgf signaling at 4 hpf *tbx16* was still expressed around the margin (Figure 6D). Also, while the early inhibition of Fgf signaling resulted in the expected loss of posterior paraxial derivatives in the embryo by 24 hpf (Figure 6I) (Draper et al., 2003), these embryos displayed normal jaw muscles in the head (Figure 6I), indicating that in the absence of Fgf signaling, anterior paraxial mesoderm derivatives are still maintained. Strikingly, loss of both Nodal and Fgf signaling abolished *tbx16* expression when treatment was performed at 4, 5 or 6 hpf (Figure 6E) and resulted in the loss of almost all mesodermal derivatives at 24 hpf (Figure 6J). Therefore, even though in the absence of Nodal signaling the formation of paraxial mesoderm derivatives can be maintained by Fgf signaling, mesoderm specification does not occur in the absence of both Nodal and Fgf signaling.

Taken together these data show that, unlike endoderm progenitors, specification of paraxial mesoderm progenitors relies on a sustained signaling input. This cannot be explained by Nodal signaling levels alone, but requires a combination of both Nodal and Fgf signaling.

### Endoderm development is robust to variation in initial progenitor number

Our proposed model for how Nodal signaling regulates the specification of endoderm versus mesoderm raises an interesting question: if the induction of endoderm is effectively a random process, how is the number of endodermal progenitors regulated? In many developmental systems, embryos are robust to small perturbations to signaling levels. For example, zebrafish embryos recover from early defects in the development of the embryonic shield in *ndr1* mutants, which are null for one of the two Nodal ligands (Dougan et al., 2003). Moreover, several recent BMP signaling studies have shown that either insufficient or excessive BMP signaling early on in development is compensated for later and therefore does not result in the dorsal–ventral patterning defects that would be expected (Rogers et al., 2020; Tuazon et al., 2020). We therefore asked whether zebrafish embryos are similarly robust to changes in the initial number of endodermal progenitors.

To perturb endodermal progenitor number, we treated embryos at sphere stage with increasing doses of the Nodal receptor inhibitor SB-505124. Increasing inhibitor dose progressively reduced levels of Nodal signaling (assayed by the number of cells in the first two cell tiers with elevated P-Smad2) (Figure 7A), and a progressive reduction in endodermal progenitor number (Figure 7B). We saw this same effect later in epiboly upon counting the number of *sox17*-positive cells at 8 hpf (Figure 7C).

**Figure 7.**
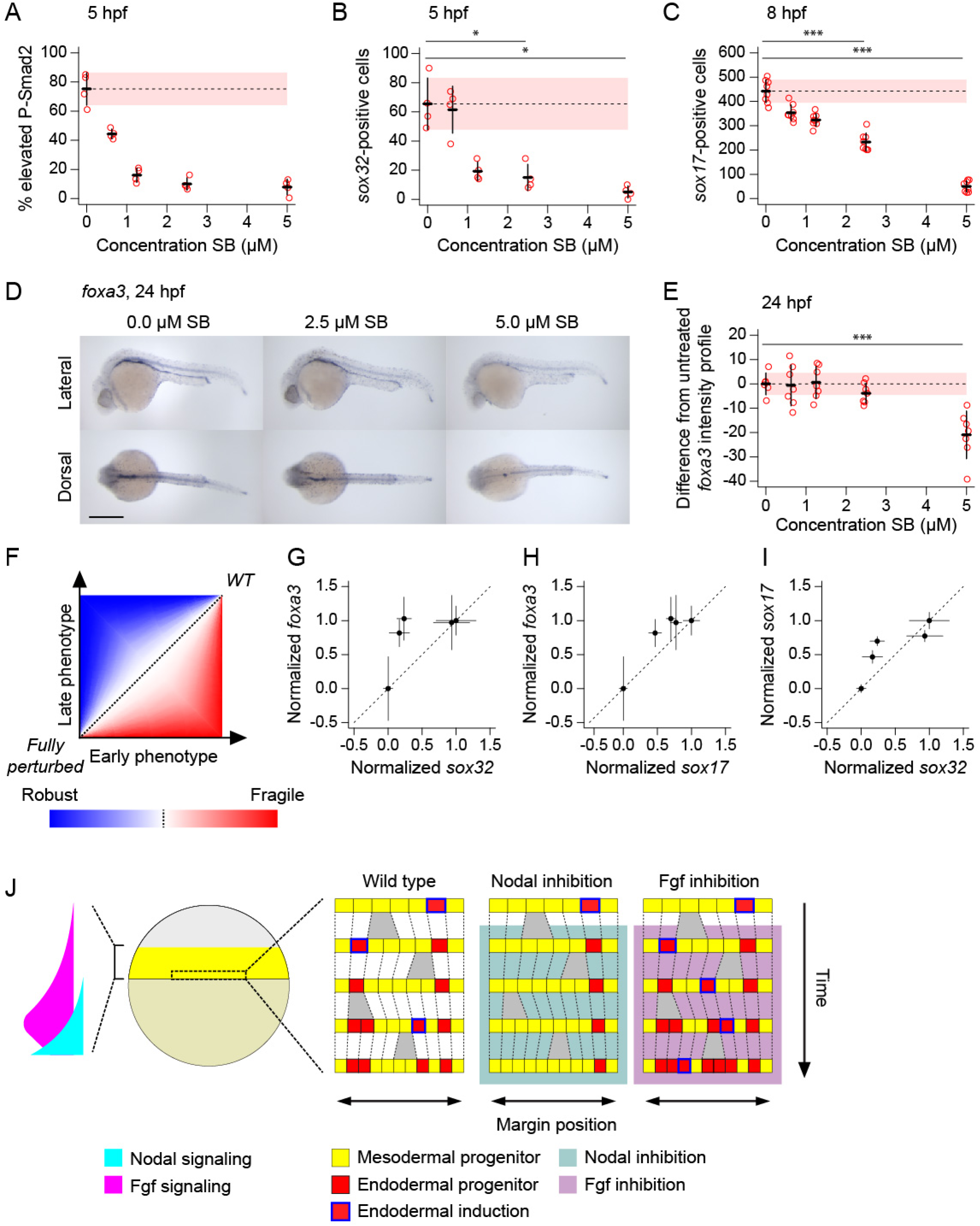
Endoderm induction is robust to variation in initial progenitor number. (A) Plot showing percentage of cells in first two cell tiers with elevated staining intensity for P-Smad2, for 5 hpf embryos treated at 4 hpf with different doses of SB-505124. Elevated P-Smad2 is defined by the mean background level for all embryos, with background for each embryo is defined by the 99^th^ percentile in P-Smad2 staining intensity for the ninth and tenth cell tiers. Four embryos per dose. Means ± SD are shown. (B) Number of *sox32*-positive cells for the same embryos as in (A). Means ± SD are shown. Statistical difference was tested using a Wilcoxon rank sum test: p < 0.05 for both 0 µM vs 2.5 µM and 0 µM vs 5 µM. (C) Number of *sox17*-positive cells for 8 hpf embryos treated at 4 hpf with different doses of SB-505124. Eight embryos per dose. Means ± SD are shown. Statistical difference was tested using a Wilcoxon rank sum test: p < 0.001 for both 0 µM vs 2.5 µM and 0 µM vs 5 µM. (D) Representative 24-hpf embryos showing *foxa3* expression on treatment with different doses of SB-505124 in dorsal and lateral views. Anterior to left; in lateral view dorsal is up. Scale bar, 250 µm. (E) Quantification of the deviation of the *foxa3* staining intensity profile from the mean untreated, for 24-hpf embryos treated at 4 hpf with different doses of SB-505124. Means ± SD are shown. Statistical difference was tested using t-test: n.s (0 µM vs 2.5 µM), p < 0.001 (0 µM vs 5 µM). Note that in A–C and E, dashed line denotes untreated mean, with pink shaded area, one SD. (F) Schematic illustrating signature of robustness and fragility in developmental systems, through plotting developmental states of embryos at early versus late time points under a range of perturbation strengths from wild type (WT) to fully perturbed. The blue region, where the deviation from WT is greater at early stages than late under a given strength perturbation (relative to fully perturbed case) represents robustness, while the red area where the converse is true, represents fragility. (G–I) Comparison of early endoderm development with later measures using quantifications in (B, C, E) to identify robustness in development as described in (F). Comparison of *sox32* counts at 5 hpf (horizontal axis) with *foxa3* expression profiles at 24 hpf (vertical axis) (G). Comparison of *sox17* counts at 8 hpf (horizontal axis) with *foxa3* expression profiles at 24 hpf (vertical axis) (H). Comparison of *sox32* counts at 5 hpf (horizontal axis) with *sox17* counts at 8 hpf (vertical axis) (I). (J) Schematic showing our model of endoderm progenitor specification in early zebrafish development. The Nodal and Fgf gradients at the margin are schematized (Economou and Hill, 2020). For details, see text. See also Figure S7.

We next asked whether this early reduction in endodermal progenitors resulted in reduced endoderm later in development. We maintained embryos until 24 hpf and visualized endodermal derivatives by in situ hybridization for *foxa3* (Ober et al., 2003) (Figure 7D). To quantitatively determine how Nodal inhibition affected gut development, we measured the staining intensity along the gut and produced a mean intensity profile for untreated (DMSO) embryos (Figure S7). Plotting the deviation from this untreated profile against Nodal inhibitor concentration revealed that there was initially no effect on the *foxa3* staining profile at lower doses of SB-505124 (Figure 7E; Figure S7E, F). Only upon treatment with 5 µM SB-505124 is the *foxa3* staining profile significantly different from the untreated embryos. Strikingly, the amount of endoderm at 24 hpf when embryos were treated with 2.5 µM SB-505124 was at wild type levels, despite the number of progenitors being reduced to around 30% of the number in untreated embryos at 5 hpf, and around 50% at 8 hpf (Figure 7B, C and E). Similar results were obtained for embryos treated with 1.25 µM SB-505124.

This reduction in the early number of endodermal progenitors without an effect on the later phenotype demonstrates that endoderm development is robust to variation in early progenitor numbers (Figure 7F-H). Indeed, even comparing the effect of perturbing the number of endodermal progenitors between 5 and 8 hpf suggests that some buffering of the early reduction in endodermal progenitor numbers occurs during epiboly. Taken together, these results indicate that zebrafish embryos are robust to significant variation in the number of early endodermal progenitors, suggesting that to reproducibly produce a viable 24 hpf embryo, the amount of early induced endoderm does not need to be tightly regulated.

## Discussion

### A model for Nodal morphogen function

Here, we describe a stochastic cell fate switching model for the role of Nodal signaling in the specification of the endoderm and mesoderm. Not only does this model differ to previous interpretations of how Nodal specifies these germ layers, but it also proposes a fundamentally different mode of action for morphogens in general (Figure 7J). In existing models, the decision as to whether a cell is fated to become endoderm or mesoderm is determined by reading out Nodal signaling relative to some absolute threshold. This is either in terms of levels of signaling, as in Wolpert’s classic solution to the French flag problem (high Nodal leads to endoderm, lower Nodal to mesoderm) (Schier, 2009; Wolpert, 1969, 2011; Zorn and Wells, 2009), or duration of signaling (a long exposure leads to endoderm, a short exposure, mesoderm) (Hagos and Dougan, 2007).

In contrast to these existing models, our present work demonstrates that endoderm and mesoderm induction are essentially independent processes. On the one hand, as we have demonstrated here and in previous studies, that Nodal works with and via Fgf to specify all cells in the margin of the embryo (up to 10 cell tiers) to a mesodermal fate (van Boxtel et al., 2018). However, endoderm induction occurs in addition to this process. High levels of Nodal signaling in the cells closest to the margin confer a competency to switch from this default mesodermal fate to the endodermal fate. In contrast to mesoderm induction, this is a stochastic process, where levels of Nodal and Fgf signaling determine the likelihood of a cell fate switch, but do not determine cell fate per se. Increasing levels of Nodal signaling make a switch in fate more likely, increasing levels of Fgf make it less likely (Figure 7J).

Consequently, in our model *sox32* expression is not simply a readout of high levels (or an extended duration) of Nodal signaling as would be the case in a classical morphogen model. Rather, *sox32* expression is regulated by a bistable switch, with Nodal signaling necessary (but not sufficient) for its transcriptional induction, but not for its maintenance and associated commitment to the endodermal lineage. Indeed, Sox32 is known to induce its own transcription in conjunction with the homeobox transcription factor Pou5fl (Lunde et al., 2004). Paraxial mesoderm fate is also maintained in the absence of Nodal signaling. However, while commitment of cells to the mesodermal fate can be lost by withdrawing both Nodal and Fgf signaling, this is not the case for endoderm where the lineage is maintained despite removal of Nodal and Fgf signaling. Thus, while endodermal cells are specified through a transcriptional switch which renders them insensitive to the withdrawal of Nodal, mesodermal progenitors require a sustained signaling input (through Nodal or Fgf).

### Nodal defines a window of competency

In our model, the switch to an endodermal fate is a probabilistic event, where at any time, all cells close to margin have the potential to switch fate (with the likelihood of switching dependent on their signaling profile). Therefore, the number of endodermal progenitors that accumulate during early epiboly is a function of the duration of exposure to Nodal: the longer the exposure, the more cells will randomly switch fate (as demonstrated in Figures 3F and 4G). Thus, rather than determining cell fate, Nodal defines a window of competency during which stochastic switching in cell fate can occur.

This is not the first time that the duration of exposure to Nodal has been implicated in this cell fate decision. As stated above, it was previously proposed that the decision between endoderm and mesoderm (and within mesoderm, different types of mesoderm) depended on the duration of exposure to Nodal (Hagos and Dougan, 2007). In particular, as the gradient of Nodal signaling grows from the margin of the embryo, only cells close to the margin that have experienced a long duration of signaling can be fated as endoderm (Economou and Hill, 2020). Our proposed role for the length of exposure to Nodal is fundamentally different to this. At any point during early epiboly between 5.0 and 5.5 hpf (late blastula to early gastrula stages) effectively any marginal cell can be induced to the endodermal fate, and all endodermal progenitors at this time have an equal endodermal potency (regardless of when they were induced). In support of this model, our scRNA-seq analysis revealed that the only endodermal marker identified during early epiboly was *sox32*. Transcriptionally, there was no other signature that distinguished endodermal progenitors from their mesodermal counterparts. Therefore, the specification of the endoderm occurs through a transcriptionally homogeneous population undergoing a transcriptional (and morphogenetic) bifurcation, with the direction that a cell takes being solely determined by whether it has by chance switched on *sox32* expression during the competency window.

### Stochasticity and robustness in early development

A major feature of our model is the importance of transcriptional and signaling variability between cells in a particular spatial location within the embryo. For example, we have demonstrated that there is considerable variation between cells within the first two cell tiers in their levels of nuclear P-Smad2, and that this variation plays a role in determining which cells are most likely to switch to an endoderm fate. It should be noted that this role for levels of Nodal signaling is very different to the role for signaling levels in classical morphogen gradient models for germ layer separation. In these models, differences in Nodal between cells are predictable and spatially structured (a gradient from the margin of the embryo), and therefore act as a source of positional information. The variation we report is spatially unstructured and therefore cannot impart positional information.

Our quantifications of the variation between signaling levels for Nodal and Fgf are based on fixed material, and therefore represent static snapshots. In many biological systems, spatially noisy processes are also temporally noisy. While this does not seem to be the case for Nodal signaling, which is integrated over time (Dubrulle et al., 2015; Economou and Hill, 2020; van Boxtel et al., 2018), in many systems signaling through P-Erk is known to stochastically fluctuate in time and as cells divide (Albeck et al., 2013; Pokrass et al., 2020; Shankaran et al., 2009; Simon et al., 2020). It is possible that part of the variability in which cells are induced to endoderm could be explained by dynamics of Erk signaling, which can be addressed in the future through the use of live reporters. Alternatively, variability could result from the inherent noisiness of a combination of molecular processes such as transcription of *sox32* itself or in the interactions with the DNA of the various transcription factors regulating *sox32* expression.

The importance of stochasticity or noise in development has become increasingly apparent in recent years (Gruenheit et al., 2018; Ohnishi et al., 2014; Priya et al., 2020). Many computational developmental models focus on how this inherent noisiness is a feature of developmental systems that needs to be buffered out to produce a precise pattern, for example by the wiring of transcriptional circuits (Diego et al., 2018; Exelby et al., 2021). In contrast, while variation and noise between cells plays an important part of our model, it is also a requirement for the patterning process. Induction of *sox32* and the endodermal lineage is a chance event. Only by allowing this to occur over hundreds of cells for an extended duration are a substantial number of endodermal progenitors induced. Without stochasticity, no endoderm would be induced.

Our model also contrasts with such buffered developmental models in that there is no requirement for a precise pattern to be initially generated. We have demonstrated that later development is robust, as the number of *sox32*-positive endodermal progenitors can be reduced to as little as one third of the number seen in untreated embryos, with little effect on the later endodermal phenotype. Moreover, as mentioned above, zebrafish embryos are remarkably resistant to dramatic changes to early BMP signaling, which are compensated for later and do not result in dorsal–ventral patterning defects that might be expected (Rogers et al., 2020; Tuazon et al., 2020). If variation in early development can be buffered out, there is no selective pressure to evolve a precise mechanism. Therefore, if robustness is a general feature of later development, we should perhaps expect to find more inherently stochastic and variable mechanisms across early development of the sort we describe here.

## STAR Methods

### Fish lines and maintenance

Zebrafish (*Danio rerio*) were housed in 28°C water (pH 7.5 and conductivity 500 µS) with a 15 h on/9 h off light cycle. All zebrafish husbandry was performed under standard conditions according to institutional (Francis Crick Institute) and national (UK) ethical and animal welfare regulations. All regulated procedures were carried out in accordance with UK Home Office regulations under project license P83B37B3C, which underwent full ethical review and approval by the Francis Crick Institute’s Animal Ethics Committee. For time courses embryos were maintained at 28°C and collected at regular intervals.

### FISH and immunofluorescence

RNAscope^®^ *in situ* hybridization (Wang et al., 2012) was performed using the RNAscope Multiplex Fluorescent v2 system, as previously described (Gross-Thebing et al., 2014; Guglielmi et al., 2021) with a few modifications. Briefly, embryos were first incubated in 2% H_2_O_2_ to inactivate endogenous peroxidases, before rehydration into PBS + 0.1% Tween-20 (PBTw), followed by hybridization overnight with specified probes at 40°C. After extensive washing in PBTw and postfixing for 10 min in 4% PFA, embryos were successively incubated with reagents Amp1-3 for 20 min at 40°C, with each amplification followed by washing with 0.2 x saline/sodium citrate buffer + 0.01% Tween-20 (SSCTw). The different probes were then visualized successively, through incubation with the HRP reagent in the appropriate channel (eg HRP-C1) for 20 min at 40°C. This was then followed by extensive washing in SSCTw and then PBTw. HRP was detected by incubating embryos with tyramide (Sigma, #T2879) coupled to either fluorescein-NHS ester (Thermo Scientific, #46410), Cy3 mono NHS ester (Sigma, #PA13101) or Cy5 mono NHS ester (Sigma, #PA15101) in PBTw for 25 min in the dark. Following the addition of 0.001% H_2_O_2_, the signal was allowed to develop for 30 min, after which the HRP was inactivated by incubating for 1 h in 3% H_2_O_2_ in PBTw to allow the detection of the next probe by repeating the process for each channel.

Immunofluorescence for P-Smad2 and P-Erk was performed as described (van Boxtel et al., 2018) with minor modifications. Embryos were first incubated in 2% H_2_O_2_ to inactivate endogenous peroxidases, before rehydration into PBS/1% Triton-X (PBTr). After incubating in acetone at −20°C, embryos were blocked in PBTr + 10% fetal bovine serum (FBS), before incubating with antibodies against P-Smad2 (Cell Signaling Technology, #8828, 1:500) or against P-Erk (Sigma, #M8159, 1:500), at 4°C overnight. For visualization, embryos were incubated for 3 h at room temperature with HRP-conjugated anti-rabbit secondary antibodies for pSmad2 (Dako, #P0448, 1:500) or anti-mouse secondary antibodies for P-Erk (Dako, #P0447, 1:500), and visualized with the tyramide system (as describe above for RNAscope assays) to increase the sensitivity of signal detection.

When combining immunofluorescence with in situ hybridizations, HRP inactivation with 3% H_2_O_2_ was followed by extensive washing in PBTr and incubation in acetone at −20°C. After that, embryos were incubated for 2 h with PBTr + 10% FBS prior to incubation with antibodies against P-Smad2 or P-Erk and were processed as for conventional immunofluorescence and RNAscope assays. For visualization, embryos were incubated for 3 h at room temperature with HRP-conjugated anti-rabbit secondary antibodies (Dako, #P0448, 1:500), and visualized with the tyramide system.

For both *in situ* hybridization and immunofluorescence, embryos were counter stained with DAPI to visualize nuclei. Embryos were then removed from the yolk, cut to allow flat-mounting, and mounted in glycerol before imaging the entire embryo on a Leica SP8 inverted confocal microscope, with an HC PL APO CS2 20x/0.75 IMM objective with the correction collar set for a glycerol immersion fluid.

### WISH

All plasmids for the generation of riboprobes, with references, can be found in the Key Resources Table. Standard WISH was performed as previously described (van Boxtel et al., 2015; van Boxtel et al., 2018). Briefly, samples were initially rehydrated to PBS/0.1% Tween (PTW). Next embryos were incubated with digoxigenin (Dig)-11-UTP-(Roche, #11209256910) labeled riboprobes in a hybridization mix containing 5% dextran sulphate overnight at 65°C. Embryos were then incubated at 4°C with anti-Dig-AP antibody overnight (Roche, #11093274910; 1:5000). After the incubation they were washed extensively in PTW before detecting alkaline phosphatase with NBT/BCIP (Sigma, # B5655).

### Drug treatments

For drug treatments, the inhibitors PD-0325901 and SB-505124 were dissolved in DMSO and directly diluted in embryo medium at 5 μM (PD-0325901) and 10 or 50 μM (SB-505124) respectively. Embryos were maintained at 28°C; time of treatment onset and durations are specified in the Figure legends.

### Image analysis

Flat mounted embryos were imaged on a Leica SP8 confocal microscopy as above. Complete embryos were captured using a tile scanning, with the voxel size set to 0.125 x 0.125 x 0.250 µm. Images were captured in 16-bit depth.

To quantify nuclear staining intensities for all cells in an embryo, nuclei were segmented using a combination of packages from the FIJI image analysis software. To identify which voxels belonged to which nuclei compared to background, a local adaptive thresholding was run on each Z-slice. This was repeated a further two times, but after reslicing the Z-stack along the X- and Y-axes, and the intersection of the foreground voxels in the three thresholded stacks was taken as nuclear voxels. As this approach did not fully separate individual nuclei, a 3-dimensional watershed was performed on the unthresholded DAPI channel (following a Gaussian blur) using the Classic Watershed function in the MorphoLibJ package. The boundaries from the water-shedding were then used to separate the thresholded nuclei. The majority of background voxels picked up by the auto-thresholding were removed by a round of erosion and dilation.

Nuclei were identified as blocks of contiguous voxels using the 3D ROI Manager from the 3D ImageJ Suite in FIJI and were used as masks to measure the mean staining intensity for all nuclei in all channels of the same image stack. Finally, background objects were excluded through a further round of filtering: objects with a mean DAPI staining intensity below 1000 were removed, as were overly flat objects (which empirically mapped to background objects) identified as those where 1.25 * volume < surface area.

To allocate a marginal position for each point, the boundary between the embryo and the YSL was marked by hand on a maximum XY projection of the Z-stack. Fitting a spline curve gave an array of regularly spaced points (at pixel increments) around the margin of the embryo. By identifying dorsal –through the location of the DFCs, or by the dorsal domain of *tbxta* – each point was allocated a marginal position (for 0 to 360 degrees where 0 degrees is dorsal). The closest marginal point was identified for each embryonic nucleus, and the associated marginal position was assigned.

### Identifying and positioning *sox32*-positive endodermal progenitors

Cells positive for *sox32* were identified relative to a staining intensity threshold. This was determined by inspection, using a value where cells clearly expressing *sox32* were recovered but background staining nuclei were excluded. As embryos within each dataset were stained together and imaged under the same settings, intensity values were comparable within a dataset. However, as different datasets were stained and imaged independently, the intensity values are not directly comparable, and so the threshold for identifying *sox32*-positive cells was identified independently for each dataset. All *sox32*-positive cells were allocated a position, φ, relative to the margin as described above.

Cells lying beyond the margin were removed as belonging to the YSL. Cells within 5 μm of the lowest nucleus in Z locally were also excluded as belonging to the YSL. DFCs were removed by excluding *sox32*-positive cells lying in the top 20% of the embryo (see Figure 2D; Figure S2G). To maintain a uniform circular distribution of *sox32*-positive cells upon removal of DFCs, the position of the remaining cells was scaled, giving a rescaled position, θ. Recording marginal positions from −180 to 180 degrees (where 0 degrees is dorsal), for cells where φ > 0, ϑ = 180-1.25*(180-φ), and where φ < 0, ϑ = −180-1.25*(−180-φ).

### Simulation

The margin of the embryo was simulated as an array of cells dividing through time. Each run of the simulation was initiated with 500 cells, proliferating to 1000 by the end of the simulation. These numbers were based on empirical counts of from the first two cell tiers of the segmented embryos (where endodermal progenitors are induced) across the early epiboly period when endoderm is induced. For simplicity it was assumed that the number of cells increases linearly through time, with each cell dividing once. Therefore, the simulation was divided into 500 time steps, with each cell in the array given an arbitrary, unique rank from 1 to 500, determining the time step each cell divides (with no cell or its daughters dividing twice during the simulation). At each time point, all cells in the array (regardless of whether or not they had yet divided) could turn on *sox32* expression, with a probability of 0.00015 (this probability was manually tuned to match the number of *sox32*-positive cells through time to the empirical data (Figure 2E)). Once induced, *sox32* expression was maintained through all remaining time steps (and in both daughters if the cell then divided). To allocate a marginal position to each cell, the array of cells was wrapped around a circle, with the two ends at 0 degrees, and all cells equally spaced.

### Sc-RNAseq analysis

The URD zebrafish cell atlas (Farrell et al., 2018) was read into R using the URD package and processed using the Seurat R toolkit https://satijalab.org/seurat/articles/get_started.html <u>(version 4.0.5)</u>. The URD R object was initially transformed into a Seurat object for downstream analysis. To investigate gene expression dynamics at the onset of gastrulation to mid-gastrulation, cells from 50% and 60% epiboly stages were initially isolated from the dataset and the raw count data were processed using Seurat’s standard data processing pipeline. For the selected stages, cells were classified into clusters (Find Cluster resolution = 0.08 for 50% epiboly and 0.05 for 60% epiboly) and uMAPs were generated using 15PCs for 50% epiboly and 20PCs for 60% epiboly. For most of the analyses in the study, cells at the margin were isolated by sub-setting cells that had at least one read of *gata5.* Alternatively, cells at the margin were obtained by isolating transcriptionally distinct clusters overlapping with the expression of *mixl1* (cluster 2 for 50% epiboly and cluster 1 and 2 for 60% epiboly). In both cases, cells were re-processed and displayed as uMAPs using 15PCs.

For cell cycle analysis, cell cycle scores were calculated for *gata5*-positive cells using the CellCycleScoring function in Seurat and cell cycle heterogeneity was visualized by performing PCA on cell cycle markers. Next, the number of cells in G1, G2/M or S phase in *sox32*-positive and -negative cells was extracted and normalized to total cell number for each population. For the analysis of *sox32* expression levels across the cell cycle, raw counts for *sox32* were extracted from *sox32*-positive cells in S phase, G2/M phase or G1 phase and plotted in GraphPad Prism. For gene enrichment analysis, the find markers function (min.diff.pct = 0.1) was used on cells positive or negative for *sox32* within the *gata5*-positive pool. After that, the ptc.1 and ptc.2 values were extracted from each gene set and the values for the top 10 genes for each population were plotted in GraphPad Prism. Finally, for clustering analysis of *gata5*-positive cells at the margin, five clustering iterations were performed using the FindNeighbors and FindClusters functions (Find cluster resolution = 0.1 to 0.5 for 50% epiboly, 0.01 to 0.5 for 60% epiboly). The same analysis was also performed on cluster-derived marginal cells for both 50 and 60% epiboly stages (Find cluster resolution = 0.05 to 0.56 for 50% epiboly, 0.01 to 0.2 for 60% epiboly). See Supplementary Files 1–5 for full details of these analyses.

## Statistical analysis

For the gene expression analysis across the phases of the cell cycle, raw counts levels were compared using a non-parametric Kruskal-Wallis test with Dunn’s multiple comparison correction (Figure 2; Figure S2). Fluorescence intensity for P-Smad2 and P-Erk between cell tiers at different time points was compared using a t-test (Figure 3D–H, Figure S3A and B). A t-test was also used to compare *foxa3* profiles (Figure 7E) and the fluorescence intensity for *tbx16* in YSL vs blastoderm cells (Figure S2A). Finally, a Wilcoxon rank sum test was used to compare the percentage of high P-Smad2 cells (Figure 4J), percentage of *sox32*-positive cells (Figure 4K) and the number of *sox17*-positive cells (Figure 4K and 4O) upon PD-0325901 and SB-505124 treatments.

## Supporting information

Key Resources Table

## Acknowledgments

We are very grateful for the support for this work from the Francis Crick Institute Aquarium, and the Light Microscopy Facility. We thank Simon Hughes, Phil Ingham, Anming Meng and Elke Ober for probes and Gavin Kelly for advice with statistical analysis. We thank Toby Andrews, James Briscoe, Matthias Carl, Foteini Papaleonidopoulou, Rashmi Priya, MC Ramel, Lilianna Solnica-Krezel, Nic Tapon, Fiona Wardle and Scott Wilcockson for helpful discussions and very useful comments on the manuscript. This research was funded in whole, or in part, by the Wellcome Trust [FC001095]. For the purpose of Open Access, the corresponding author has applied a CC BY public copyright licence to any Author Accepted Manuscript version arising from this submission. This work was supported by the Francis Crick Institute which receives its core funding from Cancer Research UK (FC001095), the UK Medical Research Council (FC001095), and the Wellcome Trust (FC001095).

The authors declare no competing interests.

## Author contributions

A.D.E., L.G. and C.S.H. conceived and designed the study. A.D.E. and L.G. planned, performed the experiments and analyzed the data, with help from P.E. A.D.E., L.G. and C.S.H wrote the manuscript. C.S.H. provided supervision and funding for the study.

## Inventory of Supplementary information

**Supplementary File 1. R markdown file for cell cycle analysis of *sox32* expression**

**Supplementary File 2. R markdown file for endoderm analysis at 50% epiboly**

**Supplementary File 3. R markdown file for cluster-based analysis at 50% epiboly**

**Supplementary File 4. R markdown file for endoderm analysis at 60% epiboly**

**Supplementary File 5. R markdown file for cluster-based analysis at 60% epiboly**

**Key Resources Table.**

**Figure S1.**
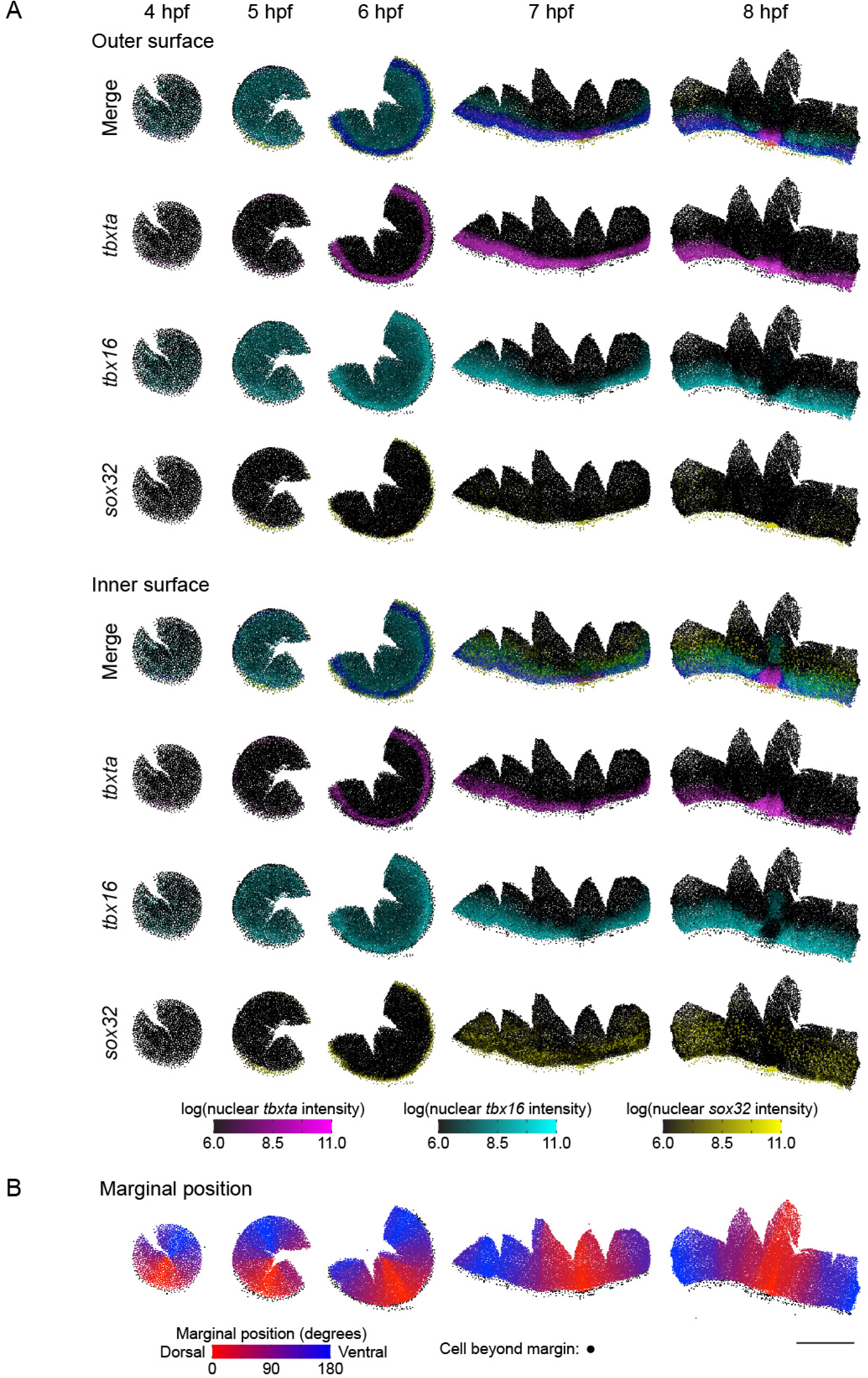
Gene expression patterns are captured through nuclear segmentation. (A) Time series of representative reconstructed embryos for each stage, showing gene expression patterns from nuclear segmentation (*tbxta*, magenta; *tbx16*, cyan; *sox32*, yellow). For each embryo, the XY positions of the centroids for each segmented nucleus are plotted, colored by the intensity of staining for each gene (nuclei below the background level of staining intensity 6.0 are colored black). The order in which nuclei are plotted is determined by their Z coordinate, giving a view of the outer surfaces of the embryos (upper panels), or by reversing this order, the inner surfaces (lower panels). (B) Reconstructed embryos in (A) with each nucleus colored by the dorsoventral position of the closest point on the embryonic margin (dorsal, red; ventral, blue). Nuclei lying beyond the margin are colored black. Scale bar, 500 μm.

**Figure S2.**
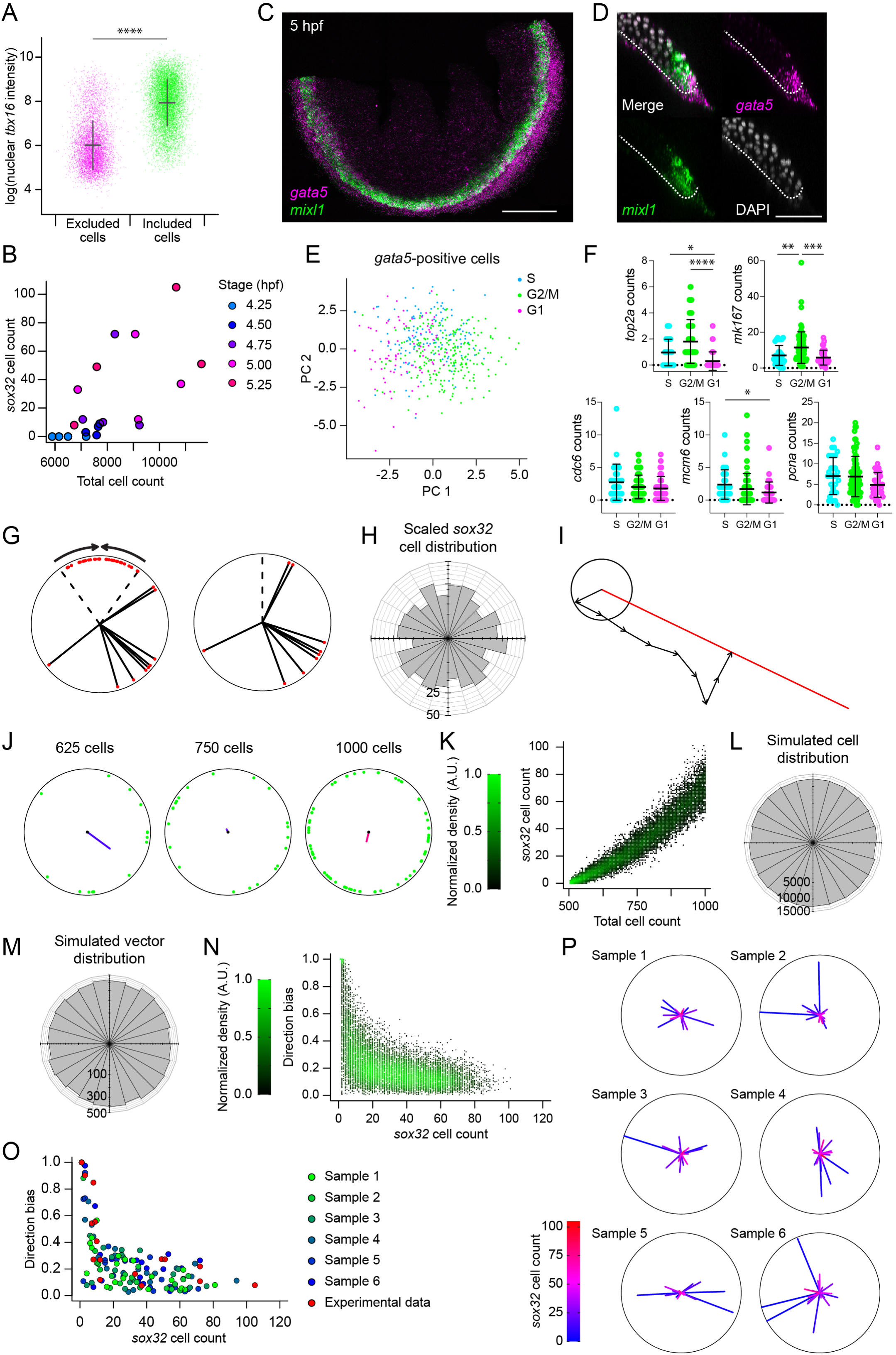
Endodermal progenitors are induced randomly in space and time. (A) Plot of nuclear *tbx16* intensity for 20 embryos collected from mass spawning across early epiboly (same embryos as used in Figure 2D), for all nuclei up to the second cell tier (the distance from the margin is less than 30 μm – this includes cells beyond the margin where the distance is negative). Cells are separated by whether they lie within the margin and are further than 5 μm from the lowest local nucleus in Z (included) or otherwise not (excluded). The lack of overlap between the two populations shows that these criteria can be used to separate the *tbx16*-positive blastodermal cells from the *tbx16*-negative YSL. Means ± SD are shown. Statistical difference was tested using a t-test. p < 0.0001. (B) Plot showing relationship between number of *sox32*-positive endodermal progenitors (YSL and DFCs removed) and total cell number for embryos collected at half hour intervals from a mass spawning. Colors indicate embryonic stage of each point. (C) Maximum projection of RNAscope in situ hybridization for *gata5* and *mixl1*, in a flat-mounted 5-hpf embryo. Scale bar, 250 µm. (D) Z-reconstruction from a 12.5 µm thick optical slice through the lateral region of the embryo, showing restriction of *gata5* expression to the most marginal cells, with *mixl1* expressed in a slightly larger domain. Dashed line marks the boundary between the embryo and YSL. Scale bar, 100 µm. (E) Principal Component Analysis (PCA) performed on *gata5*-positive cells using cell cycle markers. Color coding indicates the cell cycle phase: S phase (cyan), G2/M (green), G1 (magenta). (F) Scatterplots showing expression levels of cell cycle markers in *sox32*-positive cells. Color coding as in (E). Means ± SD are shown. Dotted line indicates zero. Statistical difference was tested using a Kruskal-Wallis test with Dunn’s multiple comparison correction. For *top2a*: p < 0.0001 (G2/M vs G1), p < 0.05 (G1 vs S), n.s (G2/M vs S). For *mk167*: p < 0.0001 (G2/M vs G1), n.s (G1 vs S), p < 0.001 (G2/M vs S). For *mcm6*: n.s (G2/M vs G1), p <0.05 (G1 vs S), n.s (G2/M vs S). For *cdc6* and *pcna* no significance was found across comparisons. (G) Cells positive for *sox32* in the top 20% of the embryo, corresponding to the dorsal side are excluded to remove DFCs (left panel). For *sox32*-positive progenitors in the ventrolateral region to conform to a uniform circular distribution, the angular positions are scaled (right panel). (H) Distribution of 489 *sox32*-positive cells around the embryonic margin as in Figure 2D, scaled to account for the removal of DFCs. The distribution of cells could not be distinguished from a uniform circular distribution (Watson’s Test for Circular Uniformity: test statistic = 0.0379, p-value > 0.10). (I) To generate an average direction vector for each embryo, the individual vectors for each cell in an embryo were summed (black arrows). A direction bias was calculated by dividing the total vector length (red line) by the length of the summed vectors, where the length of each individual vector is 1, and the total length is the number of endodermal progenitors. (J) Examples of three simulated embryos at different time steps (from 500 cells to 1000 cells), with endodermal progenitors in green, showing direction vectors and directional bias. (K) Plot showing the total number of *sox32*-positive endodermal progenitors through time (given by the total number of cells in the simulation). Based on a total of 10,000 simulations, with 100 embryos terminating at 100 equally spaced time points, ranging from 5 cell divisions to the full simulation (total cell number doubling from 500 to 1000). (L) Spatial distribution of *sox32*-positive progenitor cells pooled from all 10,000 simulations. The spatial distribution of cells is indistinguishable from uniform (Watson’s Test for Circular Uniformity: test statistic = 0.1175, p-value > 0.10). (M) Spatial distribution of direction vectors from all 10,000 simulations. The spatial distribution of vectors is indistinguishable from uniform (Watson’s Test for Circular Uniformity: test statistic = 0.0833, p-value > 0.10). (N) Plot of directional bias against the total number of *sox32*-positive cells, across all 10,000 simulations. (O) Comparison of six random subsamples of 20 simulated embryos distributed through time with the experimental data (in red). This shows the similarity in the relationship between the directional bias and the total number of *sox32*-positive cells between the experimental and simulated data. (P) Plots of directional bias for the same six embryos as in (O), showing similarity between simulated data and experimental (see Figure 2H).

**Figure S3.**
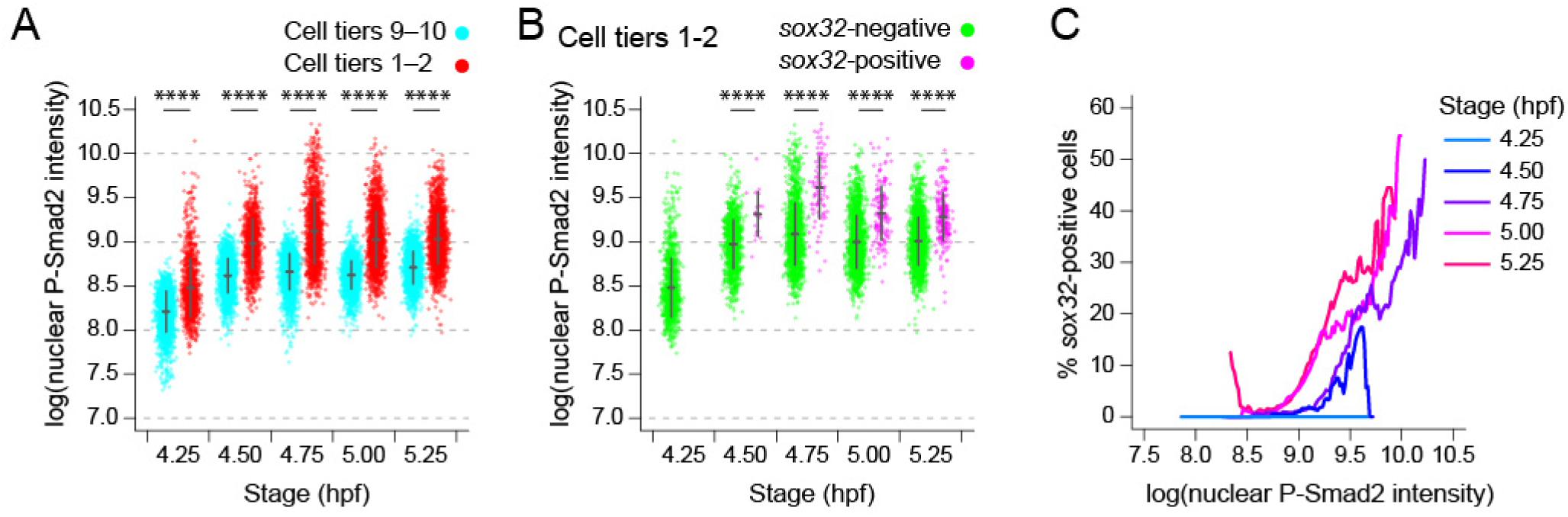
Nodal signaling levels are not deterministic for cell fate switching. (A) Plot showing P-Smad2 staining intensity for all cells in the first and second cell tiers compared to background levels (ninth and tenth cell tiers) for mass spawned dataset (Figure S2B). Means ± SD are shown. Statistical difference was tested using a t-test. For all timepoints: p < 0.0001. (B) Plot showing P-Smad2 staining intensity for all cells in the first and second cell tiers, broken down into *sox32*-positive and *sox32*-negative cells. Means ± SD are shown. Statistical difference was tested using a t-test. For all timepoints beside 4.25 hpf : p < 0.0001. (C) Traces showing the proportion of cells that are *sox32*-positive for a given P-Smad2 staining intensity for cells in the first and second cell tiers at different stages. The proportion of *sox32*-positive cells for a given level of P-Smad2 staining is based on all cells within a window of ± 10% the total range of P-Smad2 staining intensities for that stage (as this dataset was from a mass spawning and therefore more variable than the single clutch dataset, a larger smoothing window was used).

**Figure S4.**
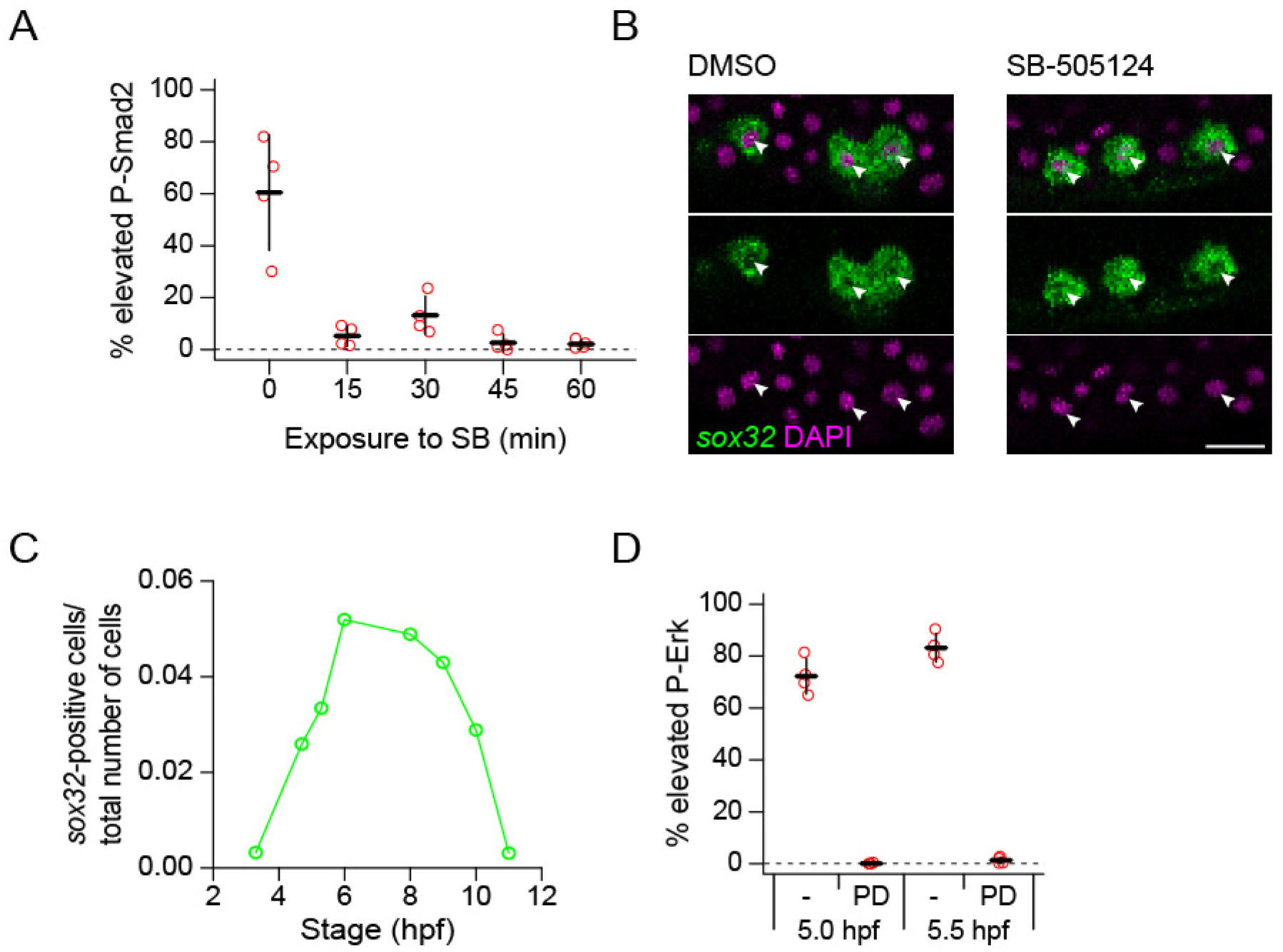
Controls for Nodal and Fgf pathway inhibitor experiments. (A) Plot showing percent of cells in the first two cell tiers with elevated staining intensity for P-Smad2, for 5.25 hpf embryos exposed to 10 μM SB-505124 for different periods of time. Upon 15 min exposure to the inhibitor, P-Smad2 is lost in the first two cell tiers. Elevated P-Smad2 is defined by the mean background level for all embryos, with background for each embryo is defined by the 99^th^ percentile in P-Smad2 staining intensity for the ninth and tenth cell tiers. Four embryos per dose. (B) Single optical slices through representative *sox32*-positive cells from embryos exposed to DMSO or 10 μM SB-505124 for 45 min, showing *sox32* transcripts present in the nuclei of inhibitor-treated embryos, comparable with control embryos (arrowheads). Scalebar, 25 μm. (C) Plot showing fraction of cells expressing > 0 read of *sox32* across gastrulation. The X axis represents developmental timing (hpf) (D) Plot showing percent of cells in the first two cell tiers with elevated staining intensity for P-Erk, for 5.0 and 5.5 hpf exposed to 5 μM PD-0325901 from 4.0 hpf. In both conditions P-Erk is lost in the first two cell tiers. Elevated P-Erk is defined by the mean background level for all embryos, with background for each embryo is defined by the 99^th^ percentile in P-Erk staining intensity for the ninth and tenth cell tiers. Four embryos per dose.

**Figure S5.**
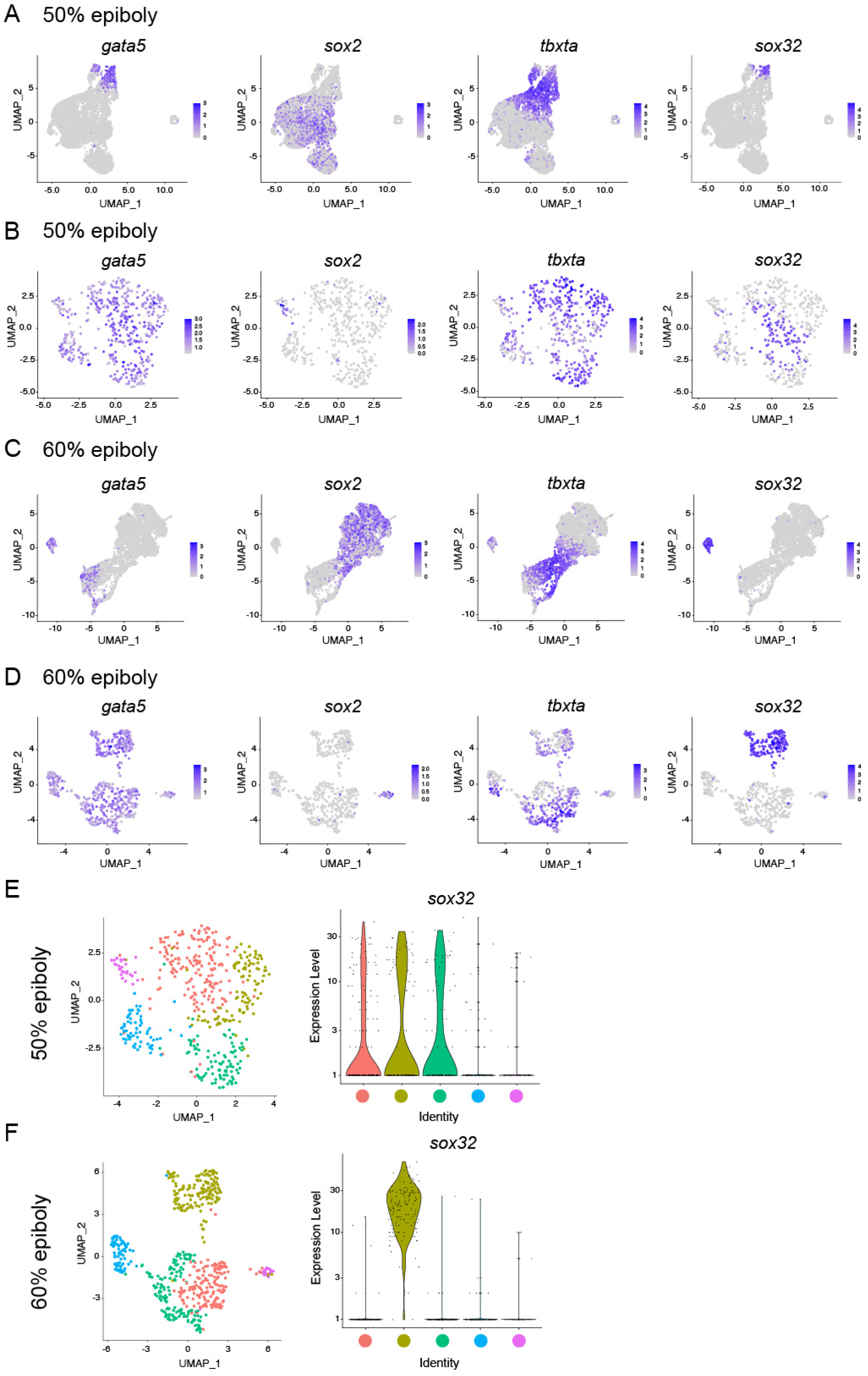
Identification of the different germ layers in scRNA-seq datasets. (A) UMAP visualization of 50% epiboly embryos showing normalized expression for (from left to right): *gata5*, *sox2*, *tbxta*, and *sox32*. (B) UMAP visualization of *gata5*-positive cells which were isolated from the 50% epiboly sample in (A). Normalized expression for the same genes as in (A) is shown. (C) UMAP visualization of 60% epiboly embryos showing normalized expression for (from left to right): *gata5*, *sox2*, *tbxta*, and *sox32*. (D) UMAP visualization of *gata5*-positive cells which were isolated from the 60% epiboly sample in (C). Normalized expression for the same genes as in (C) is shown. (E) UMAP visualization of *gata5-*positive cells at 50% epiboly showing five different clusters (left). Violin plot showing expression levels for *sox32* within each cluster (right). Color coding correlates expression levels with a specific cluster. (F) UMAP visualization of *gata5-*positive cells at 60% epiboly showing five different clusters (left). Violin plot showing expression levels for *sox32* within each cluster (right). Color coding correlates expression levels with a specific cluster.

**Figure S6.**
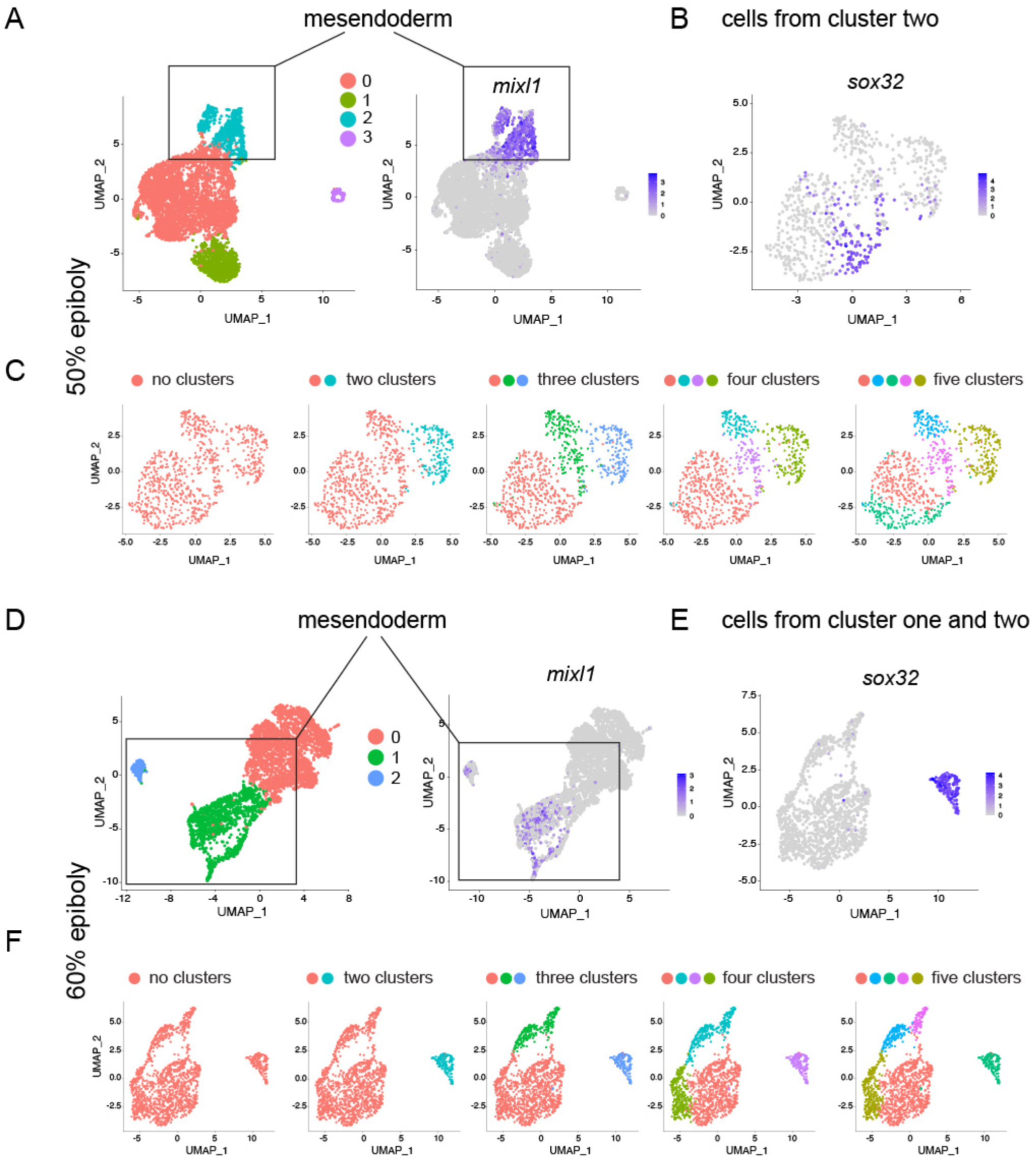
*sox32*-positive cells define a transcriptionally unique cluster at 60% epiboly, but not at 50% epiboly. (A) UMAP visualization of 50% epiboly embryos showing four clusters (left). UMAP visualization showing normalized expression of *mixl1* (right). Note that cluster two corresponds with the expression of *mixl1*. (B) UMAP visualization showing normalized expression of *sox32* within cells extracted from cluster two shown in (A) (C) UMAP visualization of cluster two cells. Cells were clustered with increasing granularity across five iterations, Find cluster resolution = 0.05 to 0.56. Color coding refers to the different clusters. Note that *sox32-*positive cells (in B) cannot by defined by a transcriptionally distinct cluster. (D) UMAP visualization of 60% epiboly embryos showing three clusters (left). UMAP visualization showing normalized expression of *mixl1* (right). Note that cluster one and two correspond to the expression of *mixl1*. (E) UMAP visualization showing normalized expression of *sox32* within cells extracted from cluster one and two shown in (D) (F) UMAP visualization of cluster one and two cells. Cells were clustered with increasing granularity across five iterations, Find cluster resolution = 0.01 to 0.2. Color coding refers to the different clusters. Note that *sox32-*positive cells in (E) define a transcriptionally distinct cluster at 60% epiboly.

**Figure S7.**
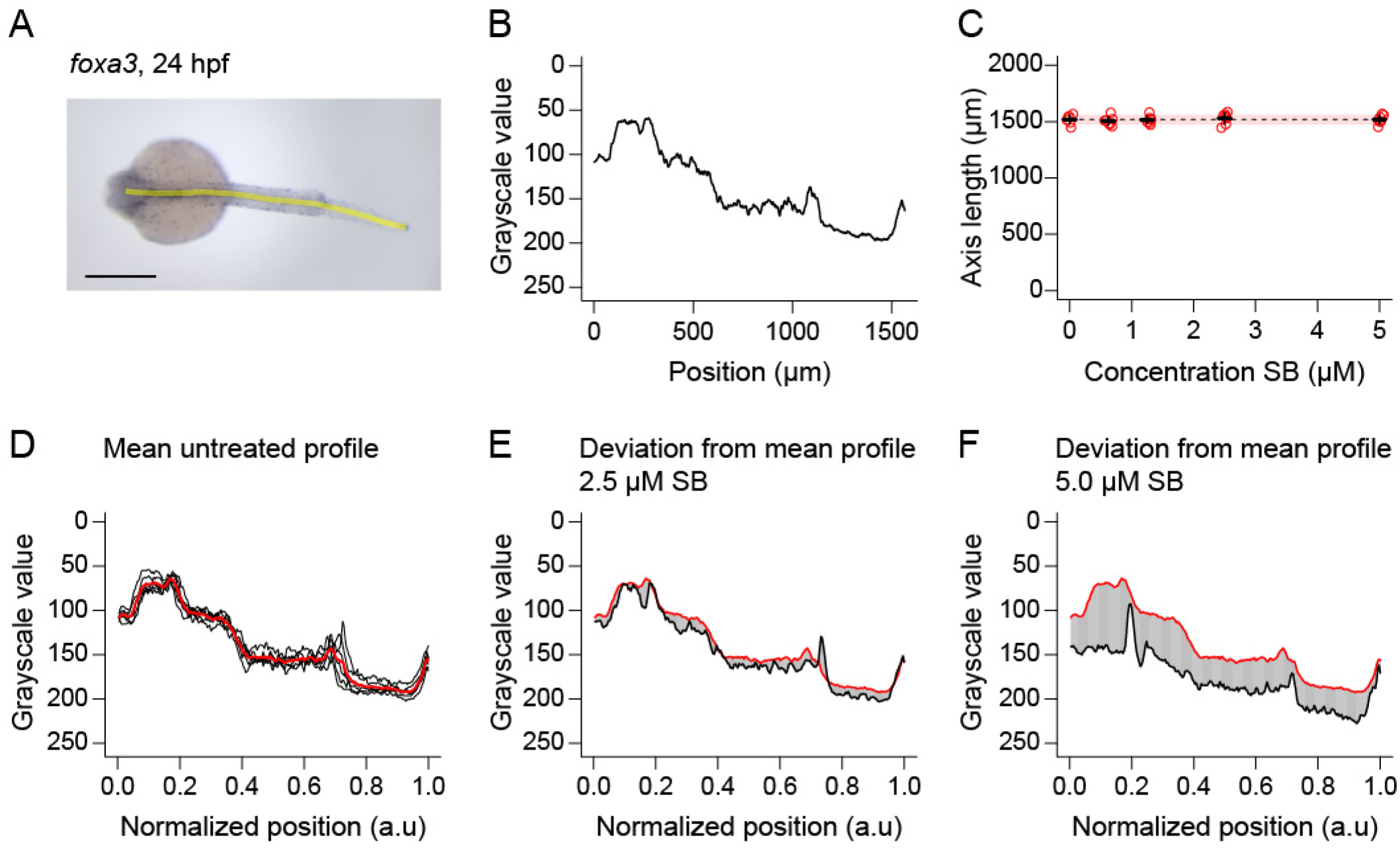
*foxa3* staining as a measure of endoderm formation at 24 hpf. (A) Dorsal view of 24-hpf embryo stained for *foxa3* expression by in situ hybridization. The yellow line marks an 18 µm wide strip along the axis of the embryo, along which the grayscale values of the staining are measured. Anterior to left; in lateral view dorsal is up. Scale bar, 250 µm. (B) Grayscale intensity profile measured from embryo in (A). Anterior end of profile at 0 µm. (C) Plot showing length of the axial profile (yellow line in (A)), for 24-hpf embryos treated at 4 hpf with different doses of SB-505124. Inhibitor treatment does not affect the gross morphology (in terms of the length of then axial profile) meaning profiles can be aligned to compare *foxa3* expression. Means ± SD are shown. Dashed line denotes untreated mean, with pink shaded area one SD. (D) Mean intensity profile (red line) made by averaging the grayscale intensity profiles for untreated six embryos (black lines). As all embryos are approximately the same length (see (C)), profiles are normalized to the length of each embryo, averaging the intensity between profiles at 250 equally spaced points along each profile. (E, F) Plots showing the intensity profiles for individual embryos treated with different doses of SB-505124 (black lines), relative to the mean profile (red line), with the offset between the lines shaded in grey. To calculate the difference from untreated *foxa3* intensity profiles for each embryo (plotted in Figure 7E), the grayscale value for each embryo profile is subtracted from the mean profile at 250 equally spaced points. Averaging the 250 values gives a score for each individual inhibitor-treated embryo.

## Notes

### Competing Interest Statement

The authors have declared no competing interest.

## References

Albeck, J.G., Mills, G.B., and Brugge, J.S. (2013). Frequency-modulated pulses of ERK activity transmit quantitative proliferation signals. Mol Cell 49, 249–261.

Alexander, J., and Stainier, D.Y. (1999). A molecular pathway leading to endoderm formation in zebrafish. Curr Biol 9, 1147–1157.

Anastasaki, C., Rauen, K.A., and Patton, E.E. (2012). Continual low-level MEK inhibition ameliorates cardio-facio-cutaneous phenotypes in zebrafish. Dis Model Mech 5, 546–552.

Briscoe, J., and Small, S. (2015). Morphogen rules: design principles of gradient-mediated embryo patterning. Development 142, 3996–4009.

Chen, Y., and Schier, A.F. (2002). Lefty proteins are long-range inhibitors of squint-mediated nodal signaling. Curr Biol 12, 2124–2128.

DaCosta Byfield, S., Major, C., Laping, N.J., and Roberts, A.B. (2004). SB-505124 is a selective inhibitor of transforming growth factor-beta type I receptors ALK4, ALK5, and ALK7. Mol Pharmacol 65, 744–752.

Diego, X., Marcon, L., Müller, P., and Sharpe, J. (2018). Key Features of Turing Systems are Determined Purely by Network Topology. Phys Rev X 8, 021071.

Dougan, S.T., Warga, R.M., Kane, D.A., Schier, A.F., and Talbot, W.S. (2003). The role of the zebrafish nodal-related genes squint and cyclops in patterning of mesendoderm. Development 130, 1837–1851.

Draper, B.W., Stock, D.W., and Kimmel, C.B. (2003). Zebrafish fgf24 functions with fgf8 to promote posterior mesodermal development. Development 130, 4639–4654.

Dubrulle, J., Jordan, B.M., Akhmetova, L., Farrell, J.A., Kim, S.H., Solnica-Krezel, L., and Schier, A.F. (2015). Response to Nodal morphogen gradient is determined by the kinetics of target gene induction. Elife 4, e05042.

Economou, A.D., and Hill, C.S. (2020). Temporal dynamics in the formation and interpretation of Nodal and BMP morphogen gradients. Curr Top Dev Biol 137, 363–389.

Exelby, K., Herrera-Delgado, E., Perez, L.G., Perez-Carrasco, R., Sagner, A., Metzis, V., Sollich, P., and Briscoe, J. (2021). Precision of tissue patterning is controlled by dynamical properties of gene regulatory networks. Development 148.

Farrell, J.A., Wang, Y., Riesenfeld, S.J., Shekhar, K., Regev, A., and Schier, A.F. (2018). Single-cell reconstruction of developmental trajectories during zebrafish embryogenesis. Science 360, eaar3131.

Feldman, B., Gates, M.A., Egan, E.S., Dougan, S.T., Rennebeck, G., Sirotkin, H.I., Schier, A.F., and Talbot, W.S. (1998). Zebrafish organizer development and germ-layer formation require nodal-related signals. Nature 395, 181–185.

Griffin, K.J., Amacher, S.L., Kimmel, C.B., and Kimelman, D. (1998). Molecular identification of spadetail: regulation of zebrafish trunk and tail mesoderm formation by T-box genes. Development 125, 3379–3388.

Gritsman, K., Zhang, J., Cheng, S., Heckscher, E., Talbot, W.S., and Schier, A.F. (1999). The EGF-CFC protein one-eyed pinhead is essential for nodal signaling. Cell 97, 121–132.

Gross-Thebing, T., Paksa, A., and Raz, E. (2014). Simultaneous high-resolution detection of multiple transcripts combined with localization of proteins in whole-mount embryos. BMC Biol 12, 55.

Gruenheit, N., Parkinson, K., Brimson, C.A., Kuwana, S., Johnson, E.J., Nagayama, K., Llewellyn, J., Salvidge, W.M., Stewart, B., Keller, T., et al. (2018). Cell Cycle Heterogeneity Can Generate Robust Cell Type Proportioning. Dev Cell 47, 494–508 e494.

Guglielmi, L., Heliot, C., Kumar, S., Alexandrov, Y., Gori, I., Papaleonidopoulou, F., Barrington, C., East, P., Economou, A.D., French, P.M.W., et al. (2021). Smad4 controls signaling robustness and morphogenesis by differentially contributing to the Nodal and BMP pathways. Nat Commun 12, 6374.

Hagos, E.G., and Dougan, S.T. (2007). Time-dependent patterning of the mesoderm and endoderm by Nodal signals in zebrafish. BMC Dev Biol 7, 22.

Hill, C.S. (2018). Spatial and temporal control of NODAL signaling. Curr Opin Cell Biol 51, 50–57.

Huang, A., and Saunders, T.E. (2020). A matter of time: Formation and interpretation of the Bicoid morphogen gradient. Curr Top Dev Biol 137, 79–117.

Kikuchi, Y., Agathon, A., Alexander, J., Thisse, C., Waldron, S., Yelon, D., Thisse, B., and Stainier, D.Y. (2001). casanova encodes a novel Sox-related protein necessary and sufficient for early endoderm formation in zebrafish. Genes Dev 15, 1493–1505.

Kikuchi, Y., Trinh, L.A., Reiter, J.F., Alexander, J., Yelon, D., and Stainier, D.Y. (2000). The zebrafish bonnie and clyde gene encodes a Mix family homeodomain protein that regulates the generation of endodermal precursors. Genes Dev 14, 1279–1289.

Kikuchi, Y., Verkade, H., Reiter, J.F., Kim, C.H., Chitnis, A.B., Kuroiwa, A., and Stainier, D.Y. (2004). Notch signaling can regulate endoderm formation in zebrafish. Dev Dyn 229, 756–762.

Lord, N.D., Carte, A.N., Abitua, P.B., and Schier, A.F. (2021). The pattern of nodal morphogen signaling is shaped by co-receptor expression. Elife 10, e54894.

Lunde, K., Belting, H.G., and Driever, W. (2004). Zebrafish pou5f1/pou2, homolog of mammalian Oct4, functions in the endoderm specification cascade. Curr Biol 14, 48–55.

Meinhardt, H., and Gierer, A. (2000). Pattern formation by local self-activation and lateral inhibition. Bioessays 22, 753–760.

Mizoguchi, T., Izawa, T., Kuroiwa, A., and Kikuchi, Y. (2006). Fgf signaling negatively regulates Nodal-dependent endoderm induction in zebrafish. Dev Biol 300, 612–622.

Nelson, A.C., Cutty, S.J., Gasiunas, S.N., Deplae, I., Stemple, D.L., and Wardle, F.C. (2017). In Vivo Regulation of the Zebrafish Endoderm Progenitor Niche by T-Box Transcription Factors. Cell Rep 19, 2782–2795.

Nowotschin, S., Hadjantonakis, A.K., and Campbell, K. (2019). The endoderm: a divergent cell lineage with many commonalities. Development 146, dev150920.

Ober, E.A., Field, H.A., and Stainier, D.Y. (2003). From endoderm formation to liver and pancreas development in zebrafish. Mech Dev 120, 5–18.

Ohnishi, Y., Huber, W., Tsumura, A., Kang, M., Xenopoulos, P., Kurimoto, K., Oles, A.K., Arauzo-Bravo, M.J., Saitou, M., Hadjantonakis, A.K., et al. (2014). Cell-to-cell expression variability followed by signal reinforcement progressively segregates early mouse lineages. Nat Cell Biol 16, 27–37.

Pages, F., and Kerridge, S. (2000). Morphogen gradients. A question of time or concentration? Trends Genet 16, 40–44.

Pokrass, M.J., Ryan, K.A., Xin, T., Pielstick, B., Timp, W., Greco, V., and Regot, S. (2020). Cell-Cycle-Dependent ERK Signaling Dynamics Direct Fate Specification in the Mammalian Preimplantation Embryo. Dev Cell 55, 328–340 e325.

Poulain, M., Furthauer, M., Thisse, B., Thisse, C., and Lepage, T. (2006). Zebrafish endoderm formation is regulated by combinatorial Nodal, FGF and BMP signalling. Development 133, 2189–2200.

Priya, R., Allanki, S., Gentile, A., Mansingh, S., Uribe, V., Maischein, H.M., and Stainier, D.Y.R. (2020). Tension heterogeneity directs form and fate to pattern the myocardial wall. Nature 588, 130–134.

Reiter, J.F., Kikuchi, Y., and Stainier, D.Y. (2001). Multiple roles for Gata5 in zebrafish endoderm formation. Development 128, 125–135.

Rogers, K.W., ElGamacy, M., Jordan, B.M., and Muller, P. (2020). Optogenetic investigation of BMP target gene expression diversity. Elife 9, e58641.

Rogers, K.W., Lord, N.D., Gagnon, J.A., Pauli, A., Zimmerman, S., Aksel, D.C., Reyon, D., Tsai, S.Q., Joung, J.K., and Schier, A.F. (2017). Nodal patterning without Lefty inhibitory feedback is functional but fragile. Elife 6, e28785.

Rogers, K.W., and Muller, P. (2019). Nodal and BMP dispersal during early zebrafish development. Dev Biol 447, 14–23.

Rogers, K.W., and Schier, A.F. (2011). Morphogen gradients: from generation to interpretation. Annu Rev Cell Dev Biol 27, 377–407.

Saijoh, Y., Adachi, H., Sakuma, R., Yeo, C.Y., Yashiro, K., Watanabe, M., Hashiguchi, H., Mochida, K., Ohishi, S., Kawabata, M., et al. (2000). Left-right asymmetric expression of lefty2 and nodal is induced by a signaling pathway that includes the transcription factor FAST2. Mol Cell 5, 35–47.

Schier, A.F. (2003). Nodal signaling in vertebrate development. Annu Rev Cell Dev Biol 19, 589–621.

Schier, A.F. (2009). Nodal morphogens. Cold Spring Harb Perspect Biol 1, a003459.

Schulte-Merker, S., van Eeden, F.J., Halpern, M.E., Kimmel, C.B., and Nusslein-Volhard, C. (1994). no tail (ntl) is the zebrafish homologue of the mouse T (Brachyury) gene. Development 120, 1009–1015.

Shankaran, H., Ippolito, D.L., Chrisler, W.B., Resat, H., Bollinger, N., Opresko, L.K., and Wiley, H.S. (2009). Rapid and sustained nuclear-cytoplasmic ERK oscillations induced by epidermal growth factor. Mol Syst Biol 5, 332.

Siefert, J.C., Clowdus, E.A., and Sansam, C.L. (2015). Cell cycle control in the early embryonic development of aquatic animal species. Comp Biochem Physiol C Toxicol Pharmacol 178, 8–15.

Simon, C.S., Rahman, S., Raina, D., Schroter, C., and Hadjantonakis, A.K. (2020). Live Visualization of ERK Activity in the Mouse Blastocyst Reveals Lineage-Specific Signaling Dynamics. Dev Cell 55, 341–353 e345.

Sun, Z., Jin, P., Tian, T., Gu, Y., Chen, Y.G., and Meng, A. (2006). Activation and roles of ALK4/ALK7-mediated maternal TGFbeta signals in zebrafish embryo. Biochem Biophys Res Commun 345, 694–703.

Tuazon, F.B., Wang, X., Andrade, J.L., Umulis, D., and Mullins, M.C. (2020). Proteolytic Restriction of Chordin Range Underlies BMP Gradient Formation. Cell Rep 32, 108039.

van Boxtel, A.L., Chesebro, J.E., Heliot, C., Ramel, M.C., Stone, R.K., and Hill, C.S. (2015). A Temporal Window for Signal Activation Dictates the Dimensions of a Nodal Signaling Domain. Dev Cell 35, 175–185.

van Boxtel, A.L., Economou, A.D., Heliot, C., and Hill, C.S. (2018). Long-Range Signaling Activation and Local Inhibition Separate the Mesoderm and Endoderm Lineages. Dev Cell 44, 179–191 e175.

Wang, F., Flanagan, J., Su, N., Wang, L.C., Bui, S., Nielson, A., Wu, X., Vo, H.T., Ma, X.J., and Luo, Y. (2012). RNAscope: a novel in situ RNA analysis platform for formalin-fixed, paraffin-embedded tissues. J Mol Diagn 14, 22–29.

Warga, R.M., and Nusslein-Volhard, C. (1999). Origin and development of the zebrafish endoderm. Development 126, 827–838.

Wolpert, L. (1969). Positional information and the spatial pattern of cellular differentiation. J Theor Biol 25, 1–47.

Wolpert, L. (2011). Positional information and patterning revisited. J Theor Biol 269, 359–365.

Zorn, A.M., and Wells, J.M. (2009). Vertebrate endoderm development and organ formation. Annu Rev Cell Dev Biol 25, 221–251.

