## Supplementary material for "Nodal signaling establishes a competency window for stochastic cell fate switching": Key Resources Table

| REAGENT or RESOURCE | SOURCE | IDENTIFIER |
|---|---|---|
| Antibodies | | |
| Anti-phospho-Smad2 (IF, Dilution: 1 in 500) | Cell Signaling Technology | Cat# 8828; RRID: AB_263108 9 |
| Anti-phospho-Erk (IF, Dilution: 1:500) | Sigma | Cat# M8159; RRID: AB_477245 |
| anti-Dig-AP (WISH, Dilution: 1 in 5000) | Roche | #11093274910 |
| HRP-conjugated anti-mouse secondary antibodies (IF, Dilution: 1:500) | Dako | # P0447 RRID:AB_2617137 |
| HRP-conjugated anti-rabbit secondary antibodies (IF, Dilution: 1:500) | Dako | #P0448 RRID: AB_2617138 |
| **Chemicals, peptides, and recombinant proteins** | | |
| Tyramide hydrochloride | Sigma | # T2879 |
| NHS-Fluorescein ester | ThermoFisher Scientific | # 46410 |
| Cy3 mono NHS ester | Sigma | # PA13101 |
| Cy5 mono NHS ester | Sigma | # PA15101 |
| Digoxigenin (Dig)-11-UTP | Roche | # 11209256910 |
| NBT/BCIP | Sigma | # B5655 |
| SB-505124 | Sigma | # S4696 |
| PD-0325901 | Merck | # 444968 |
| **Critical commercial assays** | | |
| Multiplex Fluorescent Assay v2 | ACDBio | acdbio.com |
| **Experimental models: Organisms/strains** | | |
| Zebrafish *Danio rerio*: WT | | |
| **Software and algorithms** | | |
| FIJI (ImageJ) | Schneider et al., 2012 PMID: 22930834 | https://imagej.net/Fiji/Downloads |
| R computing | The R Foundation | https://www.r-project.org/ |
| **Other** | | |
| Antisense RNA probe *tbx16*: linearize EcoR1: polymerase T7 | Osborn et al., 2020 PMID: 32345657 | |
| Antisense RNA probe *myod*: linearize Xba1: polymerase T7 | Weinberg et al., 1996 PMID: 8565839 | |
| Antisense RNA probe *sox17*: linearize NcoI: Polymerase Sp6 | Alexander and Stainier, 1999: PMID: 10531029 | |

| | | |
|---|---|---|
| Antisense RNA probe *foxa3*: linearize ApaI: polymerase T3 | Field et al, 2003 PMID: 12645931 | |
| Dr*tbxta*-C1 | ACDbio | REF: 483511 |
| Dr*sox32*-C4 | ACDbio | REF: 524941-C4 |
| Dr*mixl1*-C2 | ACDbio | REF: 850191-C2 |
| Dr*gata5*-C4 | ACDbio | REF: 850201-C4 |
| Dr*tbx16*-C3 | ACDbio | REF: 833541-C3 |
