## Supplementary File 2 for "Nodal signaling establishes a competency window for stochastic cell fate switching": Suplementary file 2.html

Endoderm analysis 50% epiboly (gata5 positive cells)


### Endoderm analysis 50% epiboly (gata5 positive cells)

###### Luca Guglielmi

#### 12/01/2022

```
# Loading required packages
library( Seurat )
```

```
## Attaching SeuratObject
```

```
library( ggplot2 )
library( dplyr )
```

```
## 
## Attaching package: 'dplyr'
```

```
## The following objects are masked from 'package:stats':
## 
##     filter, lag
```

```
## The following objects are masked from 'package:base':
## 
##     intersect, setdiff, setequal, union
```

```
library( tibble )
library( plotly )
```

```
## 
## Attaching package: 'plotly'
```

```
## The following object is masked from 'package:ggplot2':
## 
##     last_plot
```

```
## The following object is masked from 'package:stats':
## 
##     filter
```

```
## The following object is masked from 'package:graphics':
## 
##     layout
```

```
library( tidyr )
library( cowplot )
library( openxlsx )
library (readxl)
library (knitr)
```

```
# Extracting the 50% epiboly stage from the entire URD Seurat object


srat_obj_rds <- file.path("~/Documents/URD_projection/urd_srat_obj.rds")
srat_obj <- readRDS( file = srat_obj_rds )
srat_obj <- SetIdent( object = srat_obj, value = "orig.ident" )
stages <- c( "ZF50" )
zf50_srat_obj <- subset( srat_obj, idents = stages )

# This tidies up the factors and removes the empty factor levels, i.e. the other developmental stages.
zf50_srat_obj@meta.data <- droplevels( zf50_srat_obj@meta.data )
```

```
# Re-processing the 50% epiboly stage into a new Seurat object and PCA/uMAP computing

zf50_srat_obj <- zf50_srat_obj %>%
NormalizeData( ) %>%
FindVariableFeatures( ) %>%
ScaleData( ) %>%
RunPCA( )
```

```
## Centering and scaling data matrix
```

```
## PC_ 1 
## Positive:  ALDOB, HSPB1, STM, APOEB, SI:CH1073-80I24.3, SI:DKEY-56M19.5, TBX16, NNR, ZIC2B, NASP 
##     ID1, CCND1, CXCR4B, SOX11B, CITED4B, CXCR4A, SOX3, MCM6, ACIN1B, DIDO1 
##     SI:DKEY-228B2.6, APOC1, CCNA2, KRI1, MKI67, TTC31, BAZ1B, QKIA, ZC3H13, FBXO5 
## Negative:  CYT1, B3GNT5A, SI:CH73-347E22.8, ZGC:110333, FTR83, KRT97, STYK1, MGLL, PONZR5, CLDNB 
##     ZGC:193505, SI:DKEY-152P16.6, KRT92, CABZ01073832.1, SI:CH211-125O16.4, FUT9B, HSD17B14, STX11B.1, ZGC:153911, FA2H 
##     ZNF185, EVPLA, GCNT7, CEBPB, CAPN9, GRHL3, SI:DKEYP-67A8.4, PITPNC1, KALRNB, SVOPL 
## PC_ 2 
## Positive:  MIXL1, LHX1A, FOXA, GATA5, ISM1, DKK1B, OSR1, GATA6, PITX2, EFNB2A 
##     MSGN1, FSCN1A, PCDH8, CYP27C1, APLNRB, ZIC2A, EFNB2B, RND1L, MESPAB, SNAI1A 
##     TA, LFT2, SP5A, FLRT3, FP102169.1, KIRREL3L, FGF8A, TRIB3, CTH1, LFT1 
## Negative:  SOX3, ID1, SOX19A, ASB11, CXCR4B, ALDOB, SI:CH1073-80I24.3, ZIC2B, FOXD5, CABZ01070258.1 
##     FAM212AA, POLR3GLA, CITED4B, CXCR4A, MYCH, GLULB, NNR, GPD1B, MEX3B, ATP1B1A 
##     SOX2, SI:DKEY-56M19.5, CCND1, ALCAMB, PFKFB4B, SI:ZFOS-44A5.1, RGCC, CRABP2B, KAZALD2, SHISA2 
## PC_ 3 
## Positive:  EVE1, APOC1, TA, CDX4, DYNLL1, HES6, ACTB1, SI:CH211-152L15.1, WNT11, WNT8A 
##     VED, HMGN2, HSPB1, SI:CH211-114N24.6, ACTB2, MALAT1, APOEB, IM:7138239, TPBGA, MESPAB 
##     DLD, VENT, CA9, ALDH1A2, MCM6, ALDOB, SRSF5A, SP5L, TBX6L, MESPAA 
## Negative:  ZGC:110425, SI:CH211-113A14.18, SI:DKEY-108K21.10, ZGC:153405, SI:CH211-113A14.12, AKAP12B, MKI67, SI:DKEY-261M9.12, SI:CH211-113A14.24, ZGC:153409 
##     ZGC:113886, NCL, TPRB, CENPF, H1M, BAZ1B, HNRNPUB, WHSC1, RRM2, ZC3H13 
##     SI:DKEY-108K21.21, ASPM, SI:DKEY-108K21.14, MIA3, SI:ZFOS-44A5.1, MGAA, EIF3S10, USP16, SUPT6H, UACAB 
## PC_ 4 
## Positive:  CDX4, EVE1, VED, WNT11, TA, HES6, WNT8A, TBX16, TPBGA, MESPAB 
##     SEBOX, DLD, WNT5B, SI:DKEY-261J4.5, HER1, SI:CH211-106E7.2, ALDH1A2, ZGC:110425, BAMBIA, VENT 
##     TBX6L, IM:7138239, SI:DKEY-108K21.10, AKAP12B, BMP4, SI:CH211-113A14.18, RASGEF1BA, HELB, VOX, FP102169.1 
## Negative:  GSC, NOG1, FRZB, FZD8B, RIPPLY1, CHD, OTX1A, FZD8A, LFT2, FOXA2 
##     OTX1B, FOXA3, ISM1, SHISA2, SERTAD2B, CITED4B, SIX3B, KLF17, ID3, CST3 
##     PKDCCA, INSB, SLC25A33, TBR1B, TPH1B, TBX1, GATA6, ZIC3, ADMP, HER3 
## PC_ 5 
## Positive:  HMGN2, HIRIP3, CRABP2B, APOC1, SOX2, MALAT1, PKDCCB, TOP2A, MIDN, SFRP1A 
##     HSPE1, YWHAQB, ZGC:110425, AKAP12B, SP5L, SI:CH211-133N4.4, SI:CH211-137A8.4, MKI67, KRI1, KIF20BB 
##     P4HA1A, EIF3S10, SI:DKEY-108K21.10, LHX5, HISTH1L, SERPINH1B, CEP250, SLKB, PRDM14, MID1IP1B 
## Negative:  GLULB, H1M, BZW1B, CCNA1, CCNA2, ACP5A, CLDND, NANOG, ZGC:158852, BLF 
##     MKRN4, SLC16A3, ZGC:113886, CTSBA, NOTO, DDIT4, CLDN7B, TIFA, RCOR1, THY1 
##     CTH1, FAM212AA, YWHAQA, OCLNA, ARL4AB, ZGC:113424, GRA, DNAJB1B, HER7, ID1
```

```
urd_srat_obj <- zf50_srat_obj  
ElbowPlot(urd_srat_obj)
```

```
urd_srat_obj<-FindNeighbors(urd_srat_obj, dim = 1:15)
```

```
## Computing nearest neighbor graph
```

```
## Computing SNN
```

```
urd_srat_obj<-FindClusters(urd_srat_obj, resolution = 0.08)
```

```
## Modularity Optimizer version 1.3.0 by Ludo Waltman and Nees Jan van Eck
## 
## Number of nodes: 5716
## Number of edges: 187849
## 
## Running Louvain algorithm...
## Maximum modularity in 10 random starts: 0.9407
## Number of communities: 4
## Elapsed time: 0 seconds
```

```
urd_srat_obj<-RunUMAP(urd_srat_obj, dim = 1:15 )
```

```
## Warning: The default method for RunUMAP has changed from calling Python UMAP via reticulate to the R-native UWOT using the cosine metric
## To use Python UMAP via reticulate, set umap.method to 'umap-learn' and metric to 'correlation'
## This message will be shown once per session
```

```
## 13:37:04 UMAP embedding parameters a = 0.9922 b = 1.112
```

```
## 13:37:04 Read 5716 rows and found 15 numeric columns
```

```
## 13:37:04 Using Annoy for neighbor search, n_neighbors = 30
```

```
## 13:37:04 Building Annoy index with metric = cosine, n_trees = 50
```

```
## 0%   10   20   30   40   50   60   70   80   90   100%
```

```
## [----|----|----|----|----|----|----|----|----|----|
```

```
## **************************************************|
## 13:37:05 Writing NN index file to temp file /var/folders/s2/phwgmcs926x8kbjk57m5lqw14_t3jy/T//RtmpsHXR6o/file15dad56eeec98
## 13:37:05 Searching Annoy index using 1 thread, search_k = 3000
## 13:37:06 Annoy recall = 100%
## 13:37:06 Commencing smooth kNN distance calibration using 1 thread
## 13:37:07 Initializing from normalized Laplacian + noise
## 13:37:07 Commencing optimization for 500 epochs, with 225422 positive edges
## 13:37:13 Optimization finished
```

```
DimPlot (urd_srat_obj, reduction = "umap", label = TRUE,  pt.size = 1) + NoLegend()
```

```
# Expression of mesoderm, endoderm and ectoderm markers at 50% epiboly

FeaturePlot(urd_srat_obj, features = c("TA"), pt.size = 1.5)
```

```
FeaturePlot(urd_srat_obj, features = c("SOX32"), pt.size = 1.5)
```

```
FeaturePlot(urd_srat_obj, features = c("SOX2"), pt.size = 1.5)
```

```
FeaturePlot(urd_srat_obj, features = c("GATA5"), pt.size = 1.5)
```

```
# Sub-setting gata5 positive cells into a new Seurat object

SOX3250 <- subset(urd_srat_obj, subset = GATA5 > 0, slot = "counts" )
SOX3250
```

```
## An object of class Seurat 
## 17239 features across 461 samples within 1 assay 
## Active assay: RNA (17239 features, 2000 variable features)
##  3 dimensional reductions calculated: pca, tsne, umap
```

```
# Re-processing of gata5 positive cells and PCA/uMAP computing

SOX3250<- NormalizeData(SOX3250, normalization.method = "LogNormalize", scale.factor = 10000)
SOX3250<- FindVariableFeatures(SOX3250, selection.method = "vst", nfeatures = 2000)
top10 <- head(VariableFeatures(SOX3250), 10)
plot1 <- VariableFeaturePlot(SOX3250)
plot1 <- LabelPoints(plot = plot1, points = top10, repel = TRUE)
```

```
## When using repel, set xnudge and ynudge to 0 for optimal results
```

```
plot1
```

```
## Warning: Transformation introduced infinite values in continuous x-axis
```

```
## Warning: Removed 2679 rows containing missing values (geom_point).
```

```
all.genes <- rownames(SOX3250)
SOX3250 <- ScaleData(SOX3250, features = all.genes)
```

```
## Centering and scaling data matrix
```

```
SOX3250 <- RunPCA(SOX3250, features = VariableFeatures(object = SOX3250))
```

```
## PC_ 1 
## Positive:  KRT4, CAPN9, KRT92, KRT97, SI:CH211-125O16.4, KALRNB, ZGC:174935, ZGC:91849, EVPLA, KRT5 
##     SLC3A2B, PONZR5, HSD17B14, DSPA, SI:CH211-269I23.2, CLDNF, ZGC:193505, MGLL, SI:DKEYP-67A8.4, GRHL3 
##     SLC2A12, SI:CH211-69B7.6, CLDNB, EPPK1, CNTF, RASSF7B, SID1, CABZ01073832.1, STYK1, SI:DKEY-17E16.17 
## Negative:  TBX16, APOEB, ALDOB, HSPB1, APOC1, NNR, MSGN1, SI:DKEY-68O6.5, CCNA2, NOP2 
##     ZNFL2A, MESPAB, SI:CH211-152C2.3, BAMBIA, CX43.4, HES6, CCNB1, DDIT4, SI:DKEY-56M19.5, CTH1 
##     ANP32E, FP102169.1, MESPAA, EVE1, FSCN1A, APLNRB, SI:DKEY-261J4.5, IM:7138239, OSR1, ALDH1A2 
## PC_ 2 
## Positive:  GSC, CHD, NOG1, FRZB, FZD8B, RIPPLY1, OTX1A, FZD8A, KLF17, ID3 
##     SLC25A33, OTX1B, SIX3B, PKDCCA, FOXA3, FOXA2, RND1L, STM, SERTAD2B, XBP1 
##     MAGI1B, ISM1, SHISA2, GADD45BA, CST3, FOXA, LFT2, TPH1B, HER11, TBX1 
## Negative:  VED, HES6, WNT11, EVE1, TA, MESPAB, CDX4, VOX, WNT8A, BAMBIA 
##     MYCH, ID1, FP102169.1, TBX16, SI:DKEY-261J4.5, ID2A, CDCA7A, MSGN1, SI:DKEY-56M19.5, TPBGA 
##     BMP2B, ALDH1A2, HER7, POLR3GLA, DLD, VENT, PPRC1, ANP32E, SI:DKEY-27I16.2, MESPAA 
## PC_ 3 
## Positive:  SOX32, APOC1, SP5L, SOX17, HMGN2, ACKR3B, CXCR4A, SP5A, MALAT1, LMO4B 
##     SFRP1A, APOEB, FAM212AB, CDH6, FMNL2B, LMO1, CD82A, ATP1B3A, LAMB1A, RND1L 
##     IM:7138239, CDH2, SI:CH211-133N4.4, CPN1, PRDX5, IPCEF1, MSGN1, DHRS3B, SLC13A4, SZL 
## Negative:  H1M, RCOR1, RASGEF1BA, ARL4AB, MKRN4, GTF2A1, ZGC:173742, APELA, MIXL1, CITED4B 
##     SI:CH1073-80I24.3, PRICKLE1B, REEP2, ZGC:113886, SEBOX, BZW1B, OSR1, DCTPP1, SRFBP1, ACP5A 
##     ZIC2B, YWHAQA, CDH1, FGF8A, UCK2A, DNAJB1B, XBP1, CLDND, PPRC1, SI:CH211-86H15.1 
## PC_ 4 
## Positive:  SI:CH211-152C2.3, ZIC2B, GLULB, CCNA2, CITED4B, SI:CH1073-80I24.3, FOPNL, APELA, SOX19A, DYNLL1 
##     RPLP0, DDX5, CCNB1, GSC, ACTB1, OSR1, DNAJB6B, SOX3, ETS2, ID1 
##     ACTB2, CHD, CTH1, FZD8B, H2AFX, SI:DKEY-105I19.5, SPATA6L, NOTO, MEX3B, TPH1B 
## Negative:  AKAP12B, SI:DKEY-108K21.10, ZGC:110425, SI:CH211-113A14.18, MKI67, SI:DKEY-261M9.12, ZGC:153405, SI:CH211-113A14.12, EIF3S10, BX324216.1 
##     SI:CH211-113A14.24, NUCKS1A, SI:CH211-106E7.2, HNRNPUB, SI:DKEY-108K21.21, CENPF, ZNF638, CSDE1, ASPM, SI:DKEY-108K21.14 
##     SI:CH211-209J10.5, CEP350, SALL4, TPRB, ZGC:153409, SETX, CDH2, CECR2, MARCKSA, NOP14 
## PC_ 5 
## Positive:  FSCN1A, CTH1, DKK1B, GATA6, LHX1A, PITX2, FP102169.1, LFT2, FOXA, FGF8A 
##     VENT, ACTB2, CMTM7, MIXL1, IRX7, EFNB2A, RND1L, SP5A, SOX32, KRT23 
##     IRX3A, PLEKHN1, DDX4, S1PR5A, MSGN1, APLNRB, APLNRA, MESPAB, ID3, ISM1 
## Negative:  SOX3, SOX19A, ID1, FOXD5, CITED4B, CABZ01070258.1, SOX2, ZIC2B, CXCR4B, PFKFB4B 
##     FOXD3, ASB11, CRABP2B, APELA, SERPINH1B, MYCH, HIRIP3, NRARPA, ID2A, NOTO 
##     ADD3B, ACIN1A, KIF4, ALCAMB, CX43.4, PPIG, ARRDC3A, NAV2B, DLA, HSPE1
```

```
DimPlot(SOX3250, reduction = "pca")
```

```
VizDimLoadings(SOX3250, dims = 1:2, reduction = "pca")
```

```
ElbowPlot(SOX3250)
```

```
SOX3250<- FindNeighbors(SOX3250, dims = 1:15)
```

```
## Computing nearest neighbor graph
```

```
## Computing SNN
```

```
SOX3250 <- FindClusters(SOX3250, resolution = 0.1)
```

```
## Modularity Optimizer version 1.3.0 by Ludo Waltman and Nees Jan van Eck
## 
## Number of nodes: 461
## Number of edges: 16281
## 
## Running Louvain algorithm...
## Maximum modularity in 10 random starts: 0.9000
## Number of communities: 1
## Elapsed time: 0 seconds
```

```
SOX3250 <- RunUMAP(SOX3250, dims = 1:15)
```

```
## 13:37:21 UMAP embedding parameters a = 0.9922 b = 1.112
```

```
## 13:37:21 Read 461 rows and found 15 numeric columns
```

```
## 13:37:21 Using Annoy for neighbor search, n_neighbors = 30
```

```
## 13:37:21 Building Annoy index with metric = cosine, n_trees = 50
```

```
## 0%   10   20   30   40   50   60   70   80   90   100%
```

```
## [----|----|----|----|----|----|----|----|----|----|
```

```
## **************************************************|
## 13:37:21 Writing NN index file to temp file /var/folders/s2/phwgmcs926x8kbjk57m5lqw14_t3jy/T//RtmpsHXR6o/file15dad165c2c03
## 13:37:21 Searching Annoy index using 1 thread, search_k = 3000
## 13:37:21 Annoy recall = 100%
## 13:37:22 Commencing smooth kNN distance calibration using 1 thread
## 13:37:22 Initializing from normalized Laplacian + noise
## 13:37:22 Commencing optimization for 500 epochs, with 16050 positive edges
## 13:37:23 Optimization finished
```

```
# Expression of mesoderm, endoderm and ectoderm markers in gata5 positive cells

FeaturePlot(SOX3250, features = c("SOX32"), pt.size = 2)
```

```
FeaturePlot(SOX3250, features = c("TA"), pt.size = 2)
```

```
FeaturePlot(SOX3250, features = c("SOX2"), pt.size = 2)
```

```
FeaturePlot(SOX3250, features = c("GATA5"), pt.size = 2)
```

```
# clustering of gata5 positive cells with increasing granularity

DimPlot(SOX3250, reduction = "umap",label = TRUE,  pt.size = 2) + NoLegend()
```

```
SOX3250<- FindNeighbors(SOX3250, dims = 1:15)
```

```
## Computing nearest neighbor graph
```

```
## Computing SNN
```

```
SOX3250 <- FindClusters(SOX3250, resolution = 0.15)
```

```
## Modularity Optimizer version 1.3.0 by Ludo Waltman and Nees Jan van Eck
## 
## Number of nodes: 461
## Number of edges: 16281
## 
## Running Louvain algorithm...
## Maximum modularity in 10 random starts: 0.8623
## Number of communities: 2
## Elapsed time: 0 seconds
```

```
SOX3250 <- RunUMAP(SOX3250, dims = 1:15)
```

```
## 13:37:24 UMAP embedding parameters a = 0.9922 b = 1.112
```

```
## 13:37:24 Read 461 rows and found 15 numeric columns
```

```
## 13:37:24 Using Annoy for neighbor search, n_neighbors = 30
```

```
## 13:37:24 Building Annoy index with metric = cosine, n_trees = 50
```

```
## 0%   10   20   30   40   50   60   70   80   90   100%
```

```
## [----|----|----|----|----|----|----|----|----|----|
```

```
## **************************************************|
## 13:37:25 Writing NN index file to temp file /var/folders/s2/phwgmcs926x8kbjk57m5lqw14_t3jy/T//RtmpsHXR6o/file15dad538fc0e5
## 13:37:25 Searching Annoy index using 1 thread, search_k = 3000
## 13:37:25 Annoy recall = 100%
## 13:37:25 Commencing smooth kNN distance calibration using 1 thread
## 13:37:25 Initializing from normalized Laplacian + noise
## 13:37:25 Commencing optimization for 500 epochs, with 16050 positive edges
## 13:37:26 Optimization finished
```

```
DimPlot(SOX3250, reduction = "umap",label = TRUE,  pt.size = 2) + NoLegend()
```

```
SOX3250<- FindNeighbors(SOX3250, dims = 1:15)
```

```
## Computing nearest neighbor graph
```

```
## Computing SNN
```

```
SOX3250 <- FindClusters(SOX3250, resolution = 0.25)
```

```
## Modularity Optimizer version 1.3.0 by Ludo Waltman and Nees Jan van Eck
## 
## Number of nodes: 461
## Number of edges: 16281
## 
## Running Louvain algorithm...
## Maximum modularity in 10 random starts: 0.8066
## Number of communities: 3
## Elapsed time: 0 seconds
```

```
SOX3250 <- RunUMAP(SOX3250, dims = 1:15)
```

```
## 13:37:26 UMAP embedding parameters a = 0.9922 b = 1.112
```

```
## 13:37:26 Read 461 rows and found 15 numeric columns
```

```
## 13:37:26 Using Annoy for neighbor search, n_neighbors = 30
```

```
## 13:37:26 Building Annoy index with metric = cosine, n_trees = 50
```

```
## 0%   10   20   30   40   50   60   70   80   90   100%
```

```
## [----|----|----|----|----|----|----|----|----|----|
```

```
## **************************************************|
## 13:37:26 Writing NN index file to temp file /var/folders/s2/phwgmcs926x8kbjk57m5lqw14_t3jy/T//RtmpsHXR6o/file15dad56744c82
## 13:37:26 Searching Annoy index using 1 thread, search_k = 3000
## 13:37:26 Annoy recall = 100%
## 13:37:26 Commencing smooth kNN distance calibration using 1 thread
## 13:37:27 Initializing from normalized Laplacian + noise
## 13:37:27 Commencing optimization for 500 epochs, with 16050 positive edges
## 13:37:27 Optimization finished
```

```
DimPlot(SOX3250, reduction = "umap",label = TRUE,  pt.size = 2) + NoLegend()
```

```
SOX3250<- FindNeighbors(SOX3250, dims = 1:15)
```

```
## Computing nearest neighbor graph
```

```
## Computing SNN
```

```
SOX3250 <- FindClusters(SOX3250, resolution = 0.3)
```

```
## Modularity Optimizer version 1.3.0 by Ludo Waltman and Nees Jan van Eck
## 
## Number of nodes: 461
## Number of edges: 16281
## 
## Running Louvain algorithm...
## Maximum modularity in 10 random starts: 0.7832
## Number of communities: 4
## Elapsed time: 0 seconds
```

```
SOX3250 <- RunUMAP(SOX3250, dims = 1:15)
```

```
## 13:37:28 UMAP embedding parameters a = 0.9922 b = 1.112
```

```
## 13:37:28 Read 461 rows and found 15 numeric columns
```

```
## 13:37:28 Using Annoy for neighbor search, n_neighbors = 30
```

```
## 13:37:28 Building Annoy index with metric = cosine, n_trees = 50
```

```
## 0%   10   20   30   40   50   60   70   80   90   100%
```

```
## [----|----|----|----|----|----|----|----|----|----|
```

```
## **************************************************|
## 13:37:28 Writing NN index file to temp file /var/folders/s2/phwgmcs926x8kbjk57m5lqw14_t3jy/T//RtmpsHXR6o/file15dad5cdcb631
## 13:37:28 Searching Annoy index using 1 thread, search_k = 3000
## 13:37:28 Annoy recall = 100%
## 13:37:28 Commencing smooth kNN distance calibration using 1 thread
## 13:37:29 Initializing from normalized Laplacian + noise
## 13:37:29 Commencing optimization for 500 epochs, with 16050 positive edges
## 13:37:29 Optimization finished
```

```
DimPlot(SOX3250, reduction = "umap",label = TRUE,  pt.size = 2) + NoLegend()
```

```
SOX3250<- FindNeighbors(SOX3250, dims = 1:15)
```

```
## Computing nearest neighbor graph
```

```
## Computing SNN
```

```
SOX3250 <- FindClusters(SOX3250, resolution = 0.5)
```

```
## Modularity Optimizer version 1.3.0 by Ludo Waltman and Nees Jan van Eck
## 
## Number of nodes: 461
## Number of edges: 16281
## 
## Running Louvain algorithm...
## Maximum modularity in 10 random starts: 0.7070
## Number of communities: 5
## Elapsed time: 0 seconds
```

```
SOX3250 <- RunUMAP(SOX3250, dims = 1:15)
```

```
## 13:37:30 UMAP embedding parameters a = 0.9922 b = 1.112
```

```
## 13:37:30 Read 461 rows and found 15 numeric columns
```

```
## 13:37:30 Using Annoy for neighbor search, n_neighbors = 30
```

```
## 13:37:30 Building Annoy index with metric = cosine, n_trees = 50
```

```
## 0%   10   20   30   40   50   60   70   80   90   100%
```

```
## [----|----|----|----|----|----|----|----|----|----|
```

```
## **************************************************|
## 13:37:30 Writing NN index file to temp file /var/folders/s2/phwgmcs926x8kbjk57m5lqw14_t3jy/T//RtmpsHXR6o/file15dad563ba0aa
## 13:37:30 Searching Annoy index using 1 thread, search_k = 3000
## 13:37:30 Annoy recall = 100%
## 13:37:30 Commencing smooth kNN distance calibration using 1 thread
## 13:37:30 Initializing from normalized Laplacian + noise
## 13:37:30 Commencing optimization for 500 epochs, with 16050 positive edges
## 13:37:31 Optimization finished
```

```
DimPlot(SOX3250, reduction = "umap",label = TRUE,  pt.size = 2) + NoLegend()
```

```
# sox32 expression within the different clusters

VlnPlot(SOX3250, features = c("SOX32"), slot = "counts", log = TRUE)
```

```
#Comparing sox32 positive/ negative cells 

SOX3250 <- AddMetaData( object = SOX3250,
                       metadata = GetAssayData( SOX3250, slot = "counts" )[ "SOX32", ] > 0,
                       col.name = "SOX32_positive" )
SOX3250 <- SetIdent( object = SOX3250, value = "SOX32_positive" )
SOX32.markers <- FindMarkers( SOX3250, ident.2 = "FALSE", ident.1 = "TRUE", min.diff.pct = 0.1, only.pos = TRUE )

head(SOX32.markers,n = 80)
```

```
##                          p_val avg_log2FC pct.1 pct.2    p_val_adj
## SOX32             1.909539e-97  4.5019750 1.000 0.000 3.291855e-93
## CXCR4A            6.268565e-40  2.2449296 0.848 0.332 1.080638e-35
## SOX17             2.607124e-21  0.9896311 0.262 0.000 4.494421e-17
## LFT2              7.513454e-19  1.3280097 0.766 0.392 1.295244e-14
## ID3               1.586208e-18  1.0625842 0.931 0.649 2.734464e-14
## RND1L             6.806602e-17  0.9014526 0.959 0.810 1.173390e-12
## S1PR5A            3.404787e-16  0.8785415 0.434 0.111 5.869512e-12
## VGLL4L            4.280165e-16  1.2631402 0.724 0.408 7.378576e-12
## ACKR3B            6.243053e-14  0.9518130 0.228 0.019 1.076240e-09
## PLEKHN1           1.195603e-13  0.7840802 0.469 0.161 2.061100e-09
## GATA6             1.108849e-12  0.6880654 0.883 0.595 1.911545e-08
## PITX2             6.704870e-12  0.7932717 0.793 0.551 1.155852e-07
## LMO4B             2.448067e-11  0.6164483 0.269 0.054 4.220222e-07
## ISM1              1.274040e-10  0.5890314 0.848 0.566 2.196318e-06
## KRT18             4.844050e-09  1.1813358 0.503 0.247 8.350657e-05
## SP5A              7.127441e-09  0.7853685 0.731 0.516 1.228700e-04
## SFRP1A            9.842346e-09  0.7812340 0.503 0.250 1.696722e-04
## CDH6              1.723898e-08  0.4894453 0.138 0.013 2.971828e-04
## IPCEF1            2.212706e-08  0.6440476 0.483 0.237 3.814484e-04
## CMTM7             5.167596e-08  0.6111058 0.690 0.456 8.908418e-04
## FRMD4BA           8.910218e-08  0.5461026 0.545 0.278 1.536032e-03
## LMO1              1.279264e-07  0.5065024 0.379 0.155 2.205323e-03
## HER11             1.572144e-07  0.6937320 0.338 0.142 2.710218e-03
## IRX3A             1.642503e-07  0.5416849 0.490 0.256 2.831512e-03
## CPN1              3.066374e-07  0.6169176 0.290 0.108 5.286122e-03
## PLPP3             3.225428e-07  0.3390820 0.186 0.041 5.560316e-03
## TAGLN2            1.448853e-06  0.7998907 0.310 0.130 2.497677e-02
## SI:CH211-155E24.3 1.591265e-06  0.4150870 0.428 0.215 2.743182e-02
## MRPL15            1.835271e-06  0.5445580 0.283 0.114 3.163823e-02
## YWHAQB            1.996130e-06  0.5852260 0.759 0.658 3.441129e-02
## VENT              2.036844e-06  0.5713459 0.676 0.503 3.511316e-02
## FAM212AB          3.116786e-06  0.5272631 0.586 0.354 5.373028e-02
## NOCTA             5.114910e-06  0.3693388 0.186 0.054 8.817594e-02
## FGFR4             7.953047e-06  0.5315525 0.407 0.228 1.371026e-01
## GATSL2            8.556393e-06  0.5842251 0.483 0.301 1.475037e-01
## MID1IP1B          9.979724e-06  0.4898418 0.372 0.190 1.720405e-01
## VOX               1.101069e-05  0.5014089 0.869 0.759 1.898133e-01
## GSTT1B            1.195573e-05  0.3749456 0.214 0.073 2.061048e-01
## TAGAPA            1.358241e-05  0.3251496 0.138 0.032 2.341472e-01
## LYE               1.375593e-05  1.0416648 0.324 0.152 2.371385e-01
## SERTAD2B          1.467032e-05  0.3697486 0.324 0.146 2.529017e-01
## ATP1B3A           2.013993e-05  0.4439223 0.366 0.187 3.471923e-01
## ZGC:101000        2.074460e-05  0.3603884 0.172 0.051 3.576162e-01
## TIFA              2.373075e-05  0.5416758 0.745 0.630 4.090944e-01
## ZGC:112994        2.440735e-05  0.4656761 0.276 0.123 4.207583e-01
## SLC13A4           4.128166e-05  0.5215440 0.172 0.057 7.116546e-01
## GRAMD3            4.706168e-05  0.4930619 0.676 0.516 8.112963e-01
## CD63              5.866718e-05  0.3444412 0.172 0.057 1.000000e+00
## DHRS3B            8.707685e-05  0.6939694 0.200 0.079 1.000000e+00
## FZD7A             8.799355e-05  0.4398421 0.655 0.491 1.000000e+00
## SULT6B1           8.915719e-05  0.6582739 0.448 0.297 1.000000e+00
## FLRT3             9.208140e-05  0.5583285 0.738 0.608 1.000000e+00
## KRT8              9.723997e-05  0.7952764 0.248 0.114 1.000000e+00
## AGPAT2            1.567264e-04  0.2618453 0.172 0.063 1.000000e+00
## UCHL5             1.624558e-04  0.3361103 0.448 0.272 1.000000e+00
## EFNA1B            3.127976e-04  0.3776749 0.379 0.231 1.000000e+00
## FKBP7             3.388713e-04  0.6003634 0.434 0.282 1.000000e+00
## RBBP4             4.029453e-04  0.3284479 0.862 0.722 1.000000e+00
## HAS2              4.095855e-04  0.3295045 0.407 0.241 1.000000e+00
## ELOVL6            4.695705e-04  0.4200821 0.483 0.332 1.000000e+00
## FMNL2B            5.172928e-04  0.4878574 0.407 0.272 1.000000e+00
## MAPK12B           5.173299e-04  0.3543058 0.628 0.500 1.000000e+00
## CYP2AA8           5.331892e-04  0.5467142 0.621 0.478 1.000000e+00
## COMMD7            6.227060e-04  0.3343627 0.331 0.193 1.000000e+00
## SI:CH211-209J10.6 6.357296e-04  0.3549892 0.414 0.253 1.000000e+00
## DKK1B             6.536989e-04  0.4335418 0.772 0.661 1.000000e+00
## IRX7              7.062751e-04  0.3276147 0.731 0.595 1.000000e+00
## CECR2             8.147665e-04  0.3368007 0.483 0.323 1.000000e+00
## ARL6IP5A          1.001506e-03  0.3890910 0.228 0.117 1.000000e+00
## KIRREL3L          1.064309e-03  0.3844407 0.510 0.367 1.000000e+00
## CHAC1             1.166738e-03  0.2824132 0.379 0.237 1.000000e+00
## LFNG              1.486668e-03  0.3287811 0.372 0.244 1.000000e+00
## ZNF800B           1.635801e-03  0.2546924 0.214 0.104 1.000000e+00
## KRT23             2.167300e-03  0.5027673 0.455 0.335 1.000000e+00
## RGMA              2.287322e-03  0.3281138 0.228 0.117 1.000000e+00
## HER5              2.572024e-03  0.4318362 0.290 0.177 1.000000e+00
## SP5L              2.786626e-03  0.4797393 0.676 0.570 1.000000e+00
## EFNA1A            4.791862e-03  0.2571439 0.503 0.373 1.000000e+00
## SLIRP             5.816638e-03  0.3683973 0.310 0.203 1.000000e+00
## CDH2              7.068094e-03  0.3784144 0.331 0.231 1.000000e+00
```

```
SOX3250 <- AddMetaData( object = SOX3250,
                       metadata = GetAssayData( SOX3250, slot = "counts" )[ "SOX32", ] < 1,
                       col.name = "SOX32_negative" )
SOX3250 <- SetIdent( object = SOX3250, value = "SOX32_negative" )
SOX32.markers <- FindMarkers( SOX3250, ident.2 = "FALSE", ident.1 = "TRUE", min.diff.pct = 0.1, only.pos = TRUE )

head(SOX32.markers,n = 80)
```

```
##                          p_val avg_log2FC pct.1 pct.2    p_val_adj
## RASGEF1BA         2.815734e-11  0.9389797 0.500 0.186 4.854044e-07
## ZIC2B             2.690717e-10  0.7750945 0.693 0.421 4.638526e-06
## MYCH              1.395417e-09  0.8243923 0.560 0.290 2.405559e-05
## APELA             1.616250e-09  0.8037662 0.544 0.262 2.786254e-05
## WNT11             4.520187e-09  1.0471343 0.750 0.600 7.792350e-05
## ID1               4.812449e-09  1.0056132 0.668 0.407 8.296181e-05
## APLNRB            8.680267e-09  0.7066844 0.725 0.462 1.496391e-04
## OSR1              1.374534e-08  0.6558499 0.807 0.586 2.369559e-04
## PPRC1             1.436608e-08  0.6480893 0.794 0.586 2.476569e-04
## SI:CH1073-80I24.3 2.247569e-08  0.5776732 0.899 0.766 3.874584e-04
## P4HA2             4.411169e-08  0.7096082 0.538 0.283 7.604414e-04
## HES6              4.535135e-08  0.7243091 0.829 0.717 7.818119e-04
## ADD3B             6.433655e-08  0.5902940 0.535 0.276 1.109098e-03
## MIXL1             1.093050e-07  0.5617215 0.918 0.814 1.884309e-03
## WNT8A             2.626107e-07  0.6743484 0.576 0.331 4.527146e-03
## CDX4              4.560659e-07  1.0716991 0.408 0.186 7.862120e-03
## ALDH1A2           9.725713e-07  0.5229203 0.472 0.234 1.676616e-02
## CITED4B           7.360032e-06  0.5755748 0.639 0.441 1.268796e-01
## NET1              1.039560e-05  0.5600480 0.453 0.248 1.792097e-01
## CITED4A           1.747176e-05  0.3932589 0.225 0.062 3.011957e-01
## ID2A              2.799995e-05  0.5877603 0.250 0.090 4.826912e-01
## SI:CH211-86H15.1  2.890363e-05  0.5294448 0.589 0.393 4.982697e-01
## MPHOSPH10         3.960700e-05  0.4189620 0.889 0.786 6.827851e-01
## SPRY2             4.099999e-05  0.4375713 0.323 0.145 7.067989e-01
## DLD               4.627745e-05  0.3479503 0.171 0.034 7.977769e-01
## TPBGA             6.139287e-05  0.5203348 0.402 0.228 1.000000e+00
## AMOTL2A           8.684028e-05  0.4193354 0.282 0.124 1.000000e+00
## CYCSB             1.072244e-04  0.3165623 0.905 0.800 1.000000e+00
## TMEM9B            1.317444e-04  0.2961481 0.234 0.083 1.000000e+00
## PUS1              1.362282e-04  0.3065234 0.241 0.090 1.000000e+00
## MORC3B            1.399723e-04  0.4272337 0.377 0.221 1.000000e+00
## SERPINH1B         1.498959e-04  0.5280132 0.513 0.338 1.000000e+00
## FGF24             2.070539e-04  0.2573129 0.218 0.076 1.000000e+00
## DACT2             2.712931e-04  0.5524983 0.491 0.331 1.000000e+00
## GLULB             3.160492e-04  0.3849694 0.832 0.717 1.000000e+00
## DDIT4             3.327408e-04  0.3545836 0.905 0.786 1.000000e+00
## NRARPA            4.267124e-04  0.2814024 0.133 0.028 1.000000e+00
## ADMP              4.636871e-04  0.6593580 0.180 0.062 1.000000e+00
## PCDH1B            5.008117e-04  0.3602855 0.285 0.145 1.000000e+00
## SEBOX             5.283238e-04  0.4561543 0.538 0.400 1.000000e+00
## RASL11B           5.792300e-04  0.2609392 0.275 0.124 1.000000e+00
## FOXH1             6.229111e-04  0.2871710 0.297 0.145 1.000000e+00
## RTN4A             6.882175e-04  0.3567546 0.557 0.414 1.000000e+00
## PRICKLE1A         7.481331e-04  0.3705695 0.177 0.062 1.000000e+00
## HER7              7.877172e-04  0.5199763 0.320 0.179 1.000000e+00
## TXNIPA            9.202943e-04  0.3472443 0.332 0.186 1.000000e+00
## CDCA7B            1.398347e-03  0.3625134 0.794 0.676 1.000000e+00
## SIVA1             1.402128e-03  0.2705047 0.763 0.655 1.000000e+00
## TCF3B             1.405817e-03  0.3605604 0.633 0.510 1.000000e+00
## TBX6L             1.656658e-03  0.3376541 0.193 0.083 1.000000e+00
## ZGC:173742        1.725493e-03  0.3851191 0.427 0.303 1.000000e+00
## LFT1              1.887463e-03  0.4120753 0.519 0.386 1.000000e+00
## SI:DKEY-27I16.2   1.928672e-03  0.3365611 0.354 0.221 1.000000e+00
## HN1L              1.953450e-03  0.3022754 0.665 0.538 1.000000e+00
## FOXD5             1.958136e-03  0.3570972 0.294 0.159 1.000000e+00
## PRICKLE1B         2.055373e-03  0.3080926 0.269 0.145 1.000000e+00
## ARL4AB            2.137012e-03  0.3284334 0.297 0.166 1.000000e+00
## NDUFA2            2.144022e-03  0.3038009 0.310 0.186 1.000000e+00
## SI:DKEYP-118H3.6  2.204775e-03  0.2727270 0.297 0.172 1.000000e+00
## MCM6L             2.352455e-03  0.2839654 0.513 0.345 1.000000e+00
## ATP1B1A           2.732678e-03  0.2779359 0.873 0.738 1.000000e+00
## NDUFA5            2.833290e-03  0.2794624 0.342 0.207 1.000000e+00
## ELOVL4B           3.654140e-03  0.2578157 0.237 0.117 1.000000e+00
## GDF3              3.668376e-03  0.2763268 0.563 0.421 1.000000e+00
## SI:CH211-66K16.27 3.917912e-03  0.2540242 0.193 0.090 1.000000e+00
## ZGC:55413         3.924556e-03  0.3182322 0.566 0.441 1.000000e+00
## RANBP9            3.931971e-03  0.2802201 0.329 0.200 1.000000e+00
## CDC27             3.985116e-03  0.2728004 0.316 0.186 1.000000e+00
## PLK1              4.093554e-03  0.3144655 0.763 0.634 1.000000e+00
## ZC4H2             4.099718e-03  0.2560399 0.677 0.545 1.000000e+00
## CABZ01057122.1    4.209051e-03  0.3138102 0.272 0.159 1.000000e+00
## HER1              4.373876e-03  0.4089970 0.405 0.283 1.000000e+00
## ZEB1A             4.576543e-03  0.3710246 0.383 0.262 1.000000e+00
## RRM1              5.493618e-03  0.2810563 0.709 0.607 1.000000e+00
## FAM168B           5.499315e-03  0.2949272 0.285 0.166 1.000000e+00
## ETV4              5.702585e-03  0.2843583 0.585 0.476 1.000000e+00
## RDH10A            5.837024e-03  0.3051963 0.655 0.503 1.000000e+00
## CLSTN1            6.214657e-03  0.2679222 0.358 0.228 1.000000e+00
## BCL2L12           6.801724e-03  0.2839112 0.424 0.310 1.000000e+00
## PPP1R3B           7.007617e-03  0.2776729 0.503 0.379 1.000000e+00
```

```
FeaturePlot (SOX3250, features = c("SOX32_positive" ), pt.size = 2)
```

```
FeaturePlot (SOX3250, features = c("SOX32_negative" ), pt.size = 2)
```
