## Supplementary File 3 for "Nodal signaling establishes a competency window for stochastic cell fate switching": Suplementary file 3.html

Endoderm analysis 50% epiboly (cluster based)


### Endoderm analysis 50% epiboly (cluster based)

###### Luca Guglielmi

#### 03/11/2021

```
# Loading required packages
library( Seurat )
```

```
## Attaching SeuratObject
```

```
library( ggplot2 )
library( dplyr )
```

```
## 
## Attaching package: 'dplyr'
```

```
## 13:39:59 UMAP embedding parameters a = 0.9922 b = 1.112
```

```
## 13:39:59 Read 5716 rows and found 15 numeric columns
```

```
## 13:39:59 Using Annoy for neighbor search, n_neighbors = 30
```

```
## 13:39:59 Building Annoy index with metric = cosine, n_trees = 50
```

```
## 0%   10   20   30   40   50   60   70   80   90   100%
```

```
## [----|----|----|----|----|----|----|----|----|----|
```

```
## **************************************************|
## 13:39:59 Writing NN index file to temp file /var/folders/s2/phwgmcs926x8kbjk57m5lqw14_t3jy/T//Rtmp66sfad/file15ddef0b11b7
## 13:40:00 Searching Annoy index using 1 thread, search_k = 3000
## 13:40:01 Annoy recall = 100%
## 13:40:01 Commencing smooth kNN distance calibration using 1 thread
## 13:40:01 Initializing from normalized Laplacian + noise
## 13:40:02 Commencing optimization for 500 epochs, with 225422 positive edges
## 13:40:08 Optimization finished
```

```
DimPlot (urd_srat_obj, reduction = "umap", label = TRUE,  pt.size = 1) + NoLegend()
```

```
# Expression of mixl1 at 50% epiboly

FeaturePlot(urd_srat_obj, features = c("MIXL1"), pt.size = 2)
```

```
# Sub-setting cells belonging to cluster 2

SOX32502 <-subset(urd_srat_obj, idents = c("2"), slot = "counts")

SOX32502
```

```
## An object of class Seurat 
## 17239 features across 753 samples within 1 assay 
## Active assay: RNA (17239 features, 2000 variable features)
##  3 dimensional reductions calculated: pca, tsne, umap
```

```
# Re-processing of cluster 2 cells and PCA/uMAP computing


SOX32502<- NormalizeData(SOX32502, normalization.method = "LogNormalize", scale.factor = 10000)
SOX32502<- FindVariableFeatures(SOX32502, selection.method = "vst", nfeatures = 2000)
top10 <- head(VariableFeatures(SOX32502), 10)
plot1 <- VariableFeaturePlot(SOX32502)
plot1 <- LabelPoints(plot = plot1, points = top10, repel = TRUE)
```

```
## When using repel, set xnudge and ynudge to 0 for optimal results
```

```
plot1
```

```
## Warning: Transformation introduced infinite values in continuous x-axis
```

```
## Warning: Removed 2068 rows containing missing values (geom_point).
```

```
all.genes <- rownames(SOX32502)
SOX32502 <- ScaleData(SOX32502, features = all.genes)
```

```
## Centering and scaling data matrix
```

```
SOX32502 <- RunPCA(SOX32502, features = VariableFeatures(object = SOX32502))
```

```
## PC_ 1 
## Positive:  VED, WNT11, HES6, CDX4, TA, EVE1, TBX16, ID1, VOX, HER7 
##     BAMBIA, PPRC1, TPBGA, WNT8A, DLD, MYCH, SI:DKEY-56M19.5, ANP32E, BLF, AMOTL2A 
##     CDCA7A, ID2A, ASB11, ARG2, CABZ01070258.1, HER1, ALDOB, ARL5C, MESPAB, DLC 
## Negative:  GSC, RIPPLY1, FRZB, NOG1, CHD, FZD8B, ID3, OTX1A, ISM1, RND1L 
##     FZD8A, KLF17, FOXA, LFT2, OTX1B, CST3, SLC25A33, FSCN1A, SERTAD2B, SIX3B 
##     GATA6, FOXA2, PKDCCA, FOXA3, MAGI1B, EFNB2A, KRT18, TBX1, INSB, XBP1 
## PC_ 2 
## Positive:  APOC1, MSGN1, APOEB, SP5A, FP102169.1, MESPAB, VED, GATA5, IM:7138239, SOX32 
##     SP5L, MESPAA, CXCL12A, VOX, SNAI1A, RND1L, VENT, GATA6, BAMBIA, CXCR4A 
##     BMP2B, SIX4A, TP53INP2, EFNB2A, CXCL12B, FOXC1A, FAM212AB, TRIB3, CIRBPB, PLEKHN1 
## Negative:  NOTO, CHD, CITED4B, ARL4AB, GSC, ADMP, NOG1, ZIC2B, RASGEF1BA, XBP1 
##     FOXD3, GADD45BA, APELA, H1M, FOXA3, TPH1B, FOXD5, BTG2, FRZB, DACT2 
##     PRICKLE1B, ZGC:113886, RCOR1, ID1, FOXA2, FZD8B, NET1, IP6K2B, CDKN1CA, HIGD1A 
## PC_ 3 
## Positive:  CIRBPB, TA, HNRNPA0B, CX43.4, HSPB1, P4HA2, RPLP0, ADMP, DYNLL1, NOTO 
##     RASGEF1BA, SI:CH211-114N24.6, ZIC2B, APELA, MALAT1, CDKN1CA, ID1, FOXD3, FOXD5, PFKFB4B 
##     ZGC:110540, LFT1, CCNB1, IP6K2B, GLULB, CHD, SI:CH1073-80I24.3, HMGN2, MESPAA, CITED4B 
## Negative:  SI:DKEY-108K21.10, AKAP12B, ZGC:110425, SI:CH211-113A14.18, SI:DKEY-261M9.12, MKI67, ZGC:153405, SI:CH211-113A14.12, BX324216.1, TPRB 
##     STM, PLEKHN1, SI:CH211-113A14.24, SALL4, EIF3S10, CXCR4A, SOX32, SI:DKEY-108K21.14, ZGC:153409, SI:DKEY-108K21.21 
##     GATA6, CEP350, RND1L, CENPF, GOLM1, PITX2, VGLL4L, RRM2, SI:CH211-106E7.2, SOX11B 
## PC_ 4 
## Positive:  CTH1, DKK1B, CCNA2, LFT2, GATA6, ACTB2, NANOG, GTF2A1, NDR1, CDC6 
##     TIFA, TRIB3, PITX2, ID3, CMTM7, GLULB, IRX3A, CDCA7A, BLF, OSR1 
##     MKRN4, FOPNL, VGLL4L, CLDN7B, SI:DKEY-68O6.5, H1M, DCTPP1, MIXL1, MYCLA, TDGF1 
## Negative:  AKAP12B, ZGC:110425, SERPINH1B, SP5L, TOP2A, MKI67, SI:DKEY-108K21.10, CDH2, HIRIP3, ZSWIM5 
##     HSPE1, HMGN2, ADD3B, NOP2, HSPB1, CEP250, HNRNPM, SI:CH211-113A14.18, P4HA2, EIF3S10 
##     NUSAP1, CENPF, NUMA1, HISTH1L, CDX4, ARRDC3A, SFPQ, ADMP, KIF15, SI:CH211-113A14.12 
## PC_ 5 
## Positive:  APLNRB, ALDH1A2, OSR1, CCNB1, P4HA2, MIXL1, RDH10A, CTH1, WNT8A, MESPAB 
##     MSGN1, EFNB2A, MESPAA, APLNRA, FOXA, LHX1A, ZSWIM5, SNAI1A, IRX7, SI:CH73-52F24.4 
##     LFT1, FAM46BA, PCDH8, ADD3B, TA, CYP27C1, DKK1B, DDIT4, IM:7138239, EFNB2B 
## Negative:  SOX32, CXCR4A, LYE, ACKR3B, SOX17, KRT18, SULT6B1, CABZ01070258.1, VGLL4L, STM 
##     KRT4, DHRS3B, KRT8, ZGC:101000, SI:CH73-234B20.5, SOX19A, SLC43A2B, CDH6, MID1IP1A, SOX3 
##     AKAP12B, TAGLN2, CLDNE, LMO4B, FERMT1, RRM2, ARL13A, KRT5, SI:CH211-113A14.12, GPM6AB
```

```
DimPlot(SOX32502, reduction = "pca")
```

```
VizDimLoadings(SOX32502, dims = 1:2, reduction = "pca")
```

```
ElbowPlot(SOX32502)
```

```
SOX32502<- FindNeighbors(SOX32502, dims = 1:15)
```

```
## Computing nearest neighbor graph
```

```
## Computing SNN
```

```
SOX32502 <- FindClusters(SOX32502, resolution = 0.1)
```

```
## Modularity Optimizer version 1.3.0 by Ludo Waltman and Nees Jan van Eck
## 
## Number of nodes: 753
## Number of edges: 27281
## 
## Running Louvain algorithm...
## Maximum modularity in 10 random starts: 0.9043
## Number of communities: 2
## Elapsed time: 0 seconds
```

```
SOX32502 <- RunUMAP(SOX32502, dims = 1:15)
```

```
## 13:40:16 UMAP embedding parameters a = 0.9922 b = 1.112
```

```
## 13:40:16 Read 753 rows and found 15 numeric columns
```

```
## 13:40:16 Using Annoy for neighbor search, n_neighbors = 30
```

```
## 13:40:16 Building Annoy index with metric = cosine, n_trees = 50
```

```
## 0%   10   20   30   40   50   60   70   80   90   100%
```

```
## [----|----|----|----|----|----|----|----|----|----|
```

```
## **************************************************|
## 13:40:16 Writing NN index file to temp file /var/folders/s2/phwgmcs926x8kbjk57m5lqw14_t3jy/T//Rtmp66sfad/file15dde7872a074
## 13:40:16 Searching Annoy index using 1 thread, search_k = 3000
## 13:40:16 Annoy recall = 100%
## 13:40:16 Commencing smooth kNN distance calibration using 1 thread
## 13:40:16 Initializing from normalized Laplacian + noise
## 13:40:16 Commencing optimization for 500 epochs, with 27440 positive edges
## 13:40:17 Optimization finished
```

```
# Expression of mesoderm, endoderm and ectoderm markers in gata5 positive cells


FeaturePlot(SOX32502, features = c("SOX32"), pt.size = 2)
```

```
FeaturePlot(SOX32502, features = c("GATA5"), pt.size = 2)
```

```
FeaturePlot(SOX32502, features = c("TA"), pt.size = 2)
```

```
FeaturePlot(SOX32502, features = c("SOX2"), pt.size = 2)
```

```
# clustering of gata5 positive cells with increasing granularity

SOX32502<- FindNeighbors(SOX32502, dims = 1:15)
```

```
## Computing nearest neighbor graph
```

```
## Computing SNN
```

```
SOX32502 <- FindClusters(SOX32502, resolution = 0.05)
```

```
## Modularity Optimizer version 1.3.0 by Ludo Waltman and Nees Jan van Eck
## 
## Number of nodes: 753
## Number of edges: 27281
## 
## Running Louvain algorithm...
## Maximum modularity in 10 random starts: 0.9500
## Number of communities: 1
## Elapsed time: 0 seconds
```

```
SOX32502 <- RunUMAP(SOX32502, dims = 1:15)
```

```
## 13:40:19 UMAP embedding parameters a = 0.9922 b = 1.112
```

```
## 13:40:19 Read 753 rows and found 15 numeric columns
```

```
## 13:40:19 Using Annoy for neighbor search, n_neighbors = 30
```

```
## 13:40:19 Building Annoy index with metric = cosine, n_trees = 50
```

```
## 0%   10   20   30   40   50   60   70   80   90   100%
```

```
## [----|----|----|----|----|----|----|----|----|----|
```

```
## **************************************************|
## 13:40:19 Writing NN index file to temp file /var/folders/s2/phwgmcs926x8kbjk57m5lqw14_t3jy/T//Rtmp66sfad/file15dde2ae6c133
## 13:40:19 Searching Annoy index using 1 thread, search_k = 3000
## 13:40:19 Annoy recall = 100%
## 13:40:19 Commencing smooth kNN distance calibration using 1 thread
## 13:40:19 Initializing from normalized Laplacian + noise
## 13:40:20 Commencing optimization for 500 epochs, with 27440 positive edges
## 13:40:21 Optimization finished
```

```
DimPlot(SOX32502, reduction = "umap",label = TRUE,  pt.size = 2) + NoLegend()
```

```
SOX32502<- FindNeighbors(SOX32502, dims = 1:15)
```

```
## Computing nearest neighbor graph
```

```
## Computing SNN
```

```
SOX32502 <- FindClusters(SOX32502, resolution = 0.1)
```

```
## Modularity Optimizer version 1.3.0 by Ludo Waltman and Nees Jan van Eck
## 
## Number of nodes: 753
## Number of edges: 27281
## 
## Running Louvain algorithm...
## Maximum modularity in 10 random starts: 0.9043
## Number of communities: 2
## Elapsed time: 0 seconds
```

```
SOX32502 <- RunUMAP(SOX32502, dims = 1:15)
```

```
## 13:40:21 UMAP embedding parameters a = 0.9922 b = 1.112
```

```
## 13:40:21 Read 753 rows and found 15 numeric columns
```

```
## 13:40:21 Using Annoy for neighbor search, n_neighbors = 30
```

```
## 13:40:21 Building Annoy index with metric = cosine, n_trees = 50
```

```
## 0%   10   20   30   40   50   60   70   80   90   100%
```

```
## [----|----|----|----|----|----|----|----|----|----|
```

```
## **************************************************|
## 13:40:21 Writing NN index file to temp file /var/folders/s2/phwgmcs926x8kbjk57m5lqw14_t3jy/T//Rtmp66sfad/file15dde614814b5
## 13:40:21 Searching Annoy index using 1 thread, search_k = 3000
## 13:40:21 Annoy recall = 100%
## 13:40:21 Commencing smooth kNN distance calibration using 1 thread
## 13:40:22 Initializing from normalized Laplacian + noise
## 13:40:22 Commencing optimization for 500 epochs, with 27440 positive edges
## 13:40:23 Optimization finished
```

```
DimPlot(SOX32502, reduction = "umap",label = TRUE,  pt.size = 2) + NoLegend()
```

```
SOX32502<- FindNeighbors(SOX32502, dims = 1:15)
```

```
## Computing nearest neighbor graph
```

```
## Computing SNN
```

```
SOX32502 <- FindClusters(SOX32502, resolution = 0.3)
```

```
## Modularity Optimizer version 1.3.0 by Ludo Waltman and Nees Jan van Eck
## 
## Number of nodes: 753
## Number of edges: 27281
## 
## Running Louvain algorithm...
## Maximum modularity in 10 random starts: 0.8119
## Number of communities: 3
## Elapsed time: 0 seconds
```

```
SOX32502 <- RunUMAP(SOX32502, dims = 1:15)
```

```
## 13:40:23 UMAP embedding parameters a = 0.9922 b = 1.112
```

```
## 13:40:23 Read 753 rows and found 15 numeric columns
```

```
## 13:40:23 Using Annoy for neighbor search, n_neighbors = 30
```

```
## 13:40:23 Building Annoy index with metric = cosine, n_trees = 50
```

```
## 0%   10   20   30   40   50   60   70   80   90   100%
```

```
## [----|----|----|----|----|----|----|----|----|----|
```

```
## **************************************************|
## 13:40:23 Writing NN index file to temp file /var/folders/s2/phwgmcs926x8kbjk57m5lqw14_t3jy/T//Rtmp66sfad/file15dde15a6544a
## 13:40:23 Searching Annoy index using 1 thread, search_k = 3000
## 13:40:24 Annoy recall = 100%
## 13:40:24 Commencing smooth kNN distance calibration using 1 thread
## 13:40:24 Initializing from normalized Laplacian + noise
## 13:40:24 Commencing optimization for 500 epochs, with 27440 positive edges
## 13:40:25 Optimization finished
```

```
DimPlot(SOX32502, reduction = "umap",label = TRUE,  pt.size = 2) + NoLegend()
```

```
SOX32502<- FindNeighbors(SOX32502, dims = 1:15)
```

```
## Computing nearest neighbor graph
```

```
## Computing SNN
```

```
SOX32502 <- FindClusters(SOX32502, resolution = 0.43)
```

```
## Modularity Optimizer version 1.3.0 by Ludo Waltman and Nees Jan van Eck
## 
## Number of nodes: 753
## Number of edges: 27281
## 
## Running Louvain algorithm...
## Maximum modularity in 10 random starts: 0.7617
## Number of communities: 4
## Elapsed time: 0 seconds
```

```
SOX32502 <- RunUMAP(SOX32502, dims = 1:15)
```

```
## 13:40:26 UMAP embedding parameters a = 0.9922 b = 1.112
```

```
## 13:40:26 Read 753 rows and found 15 numeric columns
```

```
## 13:40:26 Using Annoy for neighbor search, n_neighbors = 30
```

```
## 13:40:26 Building Annoy index with metric = cosine, n_trees = 50
```

```
## 0%   10   20   30   40   50   60   70   80   90   100%
```

```
## [----|----|----|----|----|----|----|----|----|----|
```

```
## **************************************************|
## 13:40:26 Writing NN index file to temp file /var/folders/s2/phwgmcs926x8kbjk57m5lqw14_t3jy/T//Rtmp66sfad/file15dde29632440
## 13:40:26 Searching Annoy index using 1 thread, search_k = 3000
## 13:40:26 Annoy recall = 100%
## 13:40:26 Commencing smooth kNN distance calibration using 1 thread
## 13:40:26 Initializing from normalized Laplacian + noise
## 13:40:26 Commencing optimization for 500 epochs, with 27440 positive edges
## 13:40:27 Optimization finished
```

```
DimPlot(SOX32502, reduction = "umap",label = TRUE,  pt.size = 2) + NoLegend()
```

```
SOX32502<- FindNeighbors(SOX32502, dims = 1:15)
```

```
## Computing nearest neighbor graph
```

```
## Computing SNN
```

```
SOX32502 <- FindClusters(SOX32502, resolution = 0.56)
```

```
## Modularity Optimizer version 1.3.0 by Ludo Waltman and Nees Jan van Eck
## 
## Number of nodes: 753
## Number of edges: 27281
## 
## Running Louvain algorithm...
## Maximum modularity in 10 random starts: 0.7306
## Number of communities: 5
## Elapsed time: 0 seconds
```

```
SOX32502 <- RunUMAP(SOX32502, dims = 1:15)
```

```
## 13:40:28 UMAP embedding parameters a = 0.9922 b = 1.112
```

```
## 13:40:28 Read 753 rows and found 15 numeric columns
```

```
## 13:40:28 Using Annoy for neighbor search, n_neighbors = 30
```

```
## 13:40:28 Building Annoy index with metric = cosine, n_trees = 50
```

```
## 0%   10   20   30   40   50   60   70   80   90   100%
```

```
## [----|----|----|----|----|----|----|----|----|----|
```

```
## **************************************************|
## 13:40:28 Writing NN index file to temp file /var/folders/s2/phwgmcs926x8kbjk57m5lqw14_t3jy/T//Rtmp66sfad/file15dde3f7d4997
## 13:40:28 Searching Annoy index using 1 thread, search_k = 3000
## 13:40:28 Annoy recall = 100%
## 13:40:28 Commencing smooth kNN distance calibration using 1 thread
## 13:40:28 Initializing from normalized Laplacian + noise
## 13:40:28 Commencing optimization for 500 epochs, with 27440 positive edges
## 13:40:29 Optimization finished
```

```
DimPlot(SOX32502, reduction = "umap",label = TRUE,  pt.size = 2) + NoLegend()
```

```
#Comparing sox32 positive/ negative cells 

SOX32502 <- AddMetaData( object = SOX32502,
                       metadata = GetAssayData( SOX32502, slot = "counts" )[ "SOX32", ] > 0,
                       col.name = "SOX32_positive" )
SOX32502 <- SetIdent( object = SOX32502, value = "SOX32_positive" )
SOX32.markers <- FindMarkers( SOX32502, ident.2 = "FALSE", ident.1 = "TRUE", min.diff.pct = 0.1, only.pos = TRUE )

head(SOX32.markers,n = 80)
```

```
##                           p_val avg_log2FC pct.1 pct.2     p_val_adj
## SOX32             3.898079e-162  4.5186707 1.000 0.000 6.719898e-158
## CXCR4A             7.001121e-60  2.4479862 0.845 0.303  1.206923e-55
## SOX17              1.799543e-35  1.0189291 0.250 0.000  3.102232e-31
## ID3                1.838495e-33  1.3232889 0.923 0.545  3.169382e-29
## GATA6              4.885625e-32  1.0966555 0.875 0.407  8.422329e-28
## RND1L              2.933781e-31  1.2133550 0.952 0.708  5.057545e-27
## LFT2               7.374260e-31  1.5109702 0.738 0.308  1.271249e-26
## S1PR5A             4.575992e-28  0.9139751 0.399 0.068  7.888553e-24
## ACKR3B             5.102963e-28  1.1051120 0.220 0.005  8.796998e-24
## GATA5              2.290389e-27  1.0180953 0.851 0.492  3.948402e-23
## PITX2              6.014349e-24  1.0440002 0.780 0.419  1.036814e-19
## ISM1               5.064019e-22  0.8689818 0.833 0.441  8.729863e-18
## PLEKHN1            1.208693e-21  0.8332581 0.435 0.118  2.083666e-17
## SFRP1A             4.030245e-19  0.9938628 0.500 0.179  6.947740e-15
## FRMD4BA            1.427329e-16  0.7718358 0.512 0.193  2.460573e-12
## VGLL4L             2.165505e-16  1.2060348 0.685 0.398  3.733115e-12
## IPCEF1             8.570192e-16  0.7830087 0.458 0.173  1.477415e-11
## SP5A               2.865858e-15  0.9535169 0.696 0.424  4.940452e-11
## LMO4B              3.857835e-14  0.5312290 0.232 0.046  6.650523e-10
## LMO1               5.215647e-14  0.6094206 0.339 0.101  8.991254e-10
## KRT18              1.108157e-13  1.3260663 0.494 0.226  1.910353e-09
## CPN1               1.233056e-13  0.6323490 0.274 0.070  2.125665e-09
## CDH6               7.590786e-12  0.5152396 0.149 0.021  1.308576e-07
## HER11              5.489457e-11  0.7144998 0.327 0.123  9.463276e-07
## CMTM7              5.988798e-11  0.7201939 0.631 0.397  1.032409e-06
## SERTAD2B           2.610151e-10  0.4838467 0.310 0.108  4.499639e-06
## IRX3A              2.818025e-10  0.6041343 0.452 0.221  4.857993e-06
## GRAMD3             6.759774e-10  0.6896870 0.643 0.427  1.165317e-05
## DKK1B              7.877398e-10  0.6585869 0.762 0.530  1.357985e-05
## MID1IP1B           9.976281e-10  0.5581428 0.345 0.142  1.719811e-05
## FGFR4              1.536587e-09  0.6151553 0.369 0.168  2.648922e-05
## YWHAQB             2.068391e-09  0.6018264 0.768 0.631  3.565699e-05
## TRIB3              2.696366e-09  0.5798502 0.774 0.579  4.648265e-05
## PLPP3              4.442363e-09  0.3587362 0.173 0.041  7.658189e-05
## EFNB2A             6.293720e-09  0.5255912 0.851 0.638  1.084974e-04
## ATP1B3A            9.819407e-09  0.5369494 0.345 0.154  1.692768e-04
## NOCTA              1.193980e-08  0.3533650 0.167 0.041  2.058302e-04
## TAGLN2             1.510265e-08  0.7283416 0.292 0.118  2.603546e-04
## FLRT3              1.892572e-08  0.7059611 0.720 0.549  3.262605e-04
## ZGC:101000         2.160149e-08  0.5148346 0.167 0.043  3.723881e-04
## MRPL15             2.228580e-08  0.5283057 0.256 0.097  3.841849e-04
## HAS2               3.127085e-08  0.4628612 0.375 0.173  5.390781e-04
## FZD7A              4.367012e-08  0.5789488 0.625 0.409  7.528293e-04
## CD63               4.498852e-08  0.3671628 0.161 0.041  7.755570e-04
## DHRS3B             9.091271e-08  0.7272075 0.190 0.062  1.567244e-03
## LYE                1.794716e-07  1.0703863 0.310 0.142  3.093911e-03
## FAM212AB           2.600268e-07  0.4992692 0.548 0.330  4.482602e-03
## KRT23              3.341541e-07  0.7668058 0.446 0.268  5.760483e-03
## LFNG               5.042725e-07  0.4565634 0.345 0.176  8.693153e-03
## ELOVL6             7.228258e-07  0.5285109 0.464 0.275  1.246079e-02
## GSTT1B             7.236041e-07  0.3564988 0.202 0.072  1.247421e-02
## MAPK12B            9.019677e-07  0.4597841 0.613 0.444  1.554902e-02
## ZGC:112994         1.312251e-06  0.4642943 0.262 0.116  2.262190e-02
## GATSL2             1.402478e-06  0.5285828 0.446 0.275  2.417731e-02
## TIFA               1.532077e-06  0.5715322 0.714 0.600  2.641148e-02
## VOX                1.722207e-06  0.4190188 0.857 0.708  2.968913e-02
## AGPAT2             1.735302e-06  0.2614554 0.161 0.051  2.991487e-02
## SLC13A4            2.336793e-06  0.4585909 0.149 0.046  4.028398e-02
## VENT               2.603552e-06  0.5653924 0.607 0.462  4.488264e-02
## EFNA1A             2.704857e-06  0.4309316 0.464 0.282  4.662903e-02
## SI:CH211-209J10.6  6.935385e-06  0.4088220 0.393 0.221  1.195591e-01
## EFNA1B             9.829982e-06  0.4132447 0.351 0.195  1.694591e-01
## FMNL2B             1.197407e-05  0.5215801 0.387 0.243  2.064210e-01
## SCML2              1.958498e-05  0.3803132 0.488 0.313  3.376255e-01
## HER5               2.084950e-05  0.5092889 0.298 0.164  3.594245e-01
## SI:CH211-155E24.3  2.269764e-05  0.3284472 0.399 0.234  3.912847e-01
## IRX7               2.291920e-05  0.3672601 0.685 0.518  3.951041e-01
## HS6ST2             2.635765e-05  0.5932951 0.524 0.398  4.543795e-01
## KIRREL3L           2.667131e-05  0.4426952 0.494 0.333  4.597866e-01
## ARL4D              2.801369e-05  0.4475317 0.381 0.234  4.829280e-01
## NCKAP5L            5.361197e-05  0.3037842 0.262 0.132  9.242168e-01
## SASH1A             5.491009e-05  0.3416634 0.565 0.400  9.465951e-01
## RAP2B              6.118720e-05  0.4427303 0.631 0.506  1.000000e+00
## ARL4CB             9.392196e-05  0.2545526 0.321 0.176  1.000000e+00
## KRT8               1.869125e-04  0.8028815 0.232 0.123  1.000000e+00
## FZD8A              2.350400e-04  0.2963567 0.286 0.156  1.000000e+00
## SULT6B1            3.301145e-04  0.7607696 0.405 0.301  1.000000e+00
## RGMA               4.595186e-04  0.3462153 0.214 0.111  1.000000e+00
## CECR2              6.261122e-04  0.3357837 0.458 0.321  1.000000e+00
## SI:CH211-209A2.1   7.003590e-04  0.3797138 0.369 0.236  1.000000e+00
```

```
SOX32502 <- AddMetaData( object = SOX32502,
                       metadata = GetAssayData( SOX32502, slot = "counts" )[ "SOX32", ] < 1,
                       col.name = "SOX32_negative" )
SOX32502 <- SetIdent( object = SOX32502, value = "SOX32_negative" )
SOX32.markers <- FindMarkers( SOX32502, ident.2 = "FALSE", ident.1 = "TRUE", min.diff.pct = 0.1, only.pos = TRUE )

head(SOX32.markers,n = 80)
```

```
##                          p_val avg_log2FC pct.1 pct.2    p_val_adj
## TA                5.810659e-32  1.4298784 0.933 0.810 1.001699e-27
## RASGEF1BA         5.011123e-21  1.2486031 0.597 0.190 8.638676e-17
## APELA             5.059845e-19  1.1771592 0.617 0.262 8.722666e-15
## WNT11             9.319149e-17  1.1072810 0.802 0.589 1.606528e-12
## HES6              1.594424e-16  0.9205216 0.850 0.679 2.748628e-12
## CDX4              4.318961e-16  1.3775291 0.508 0.167 7.445456e-12
## ID1               7.220878e-16  1.2262897 0.699 0.393 1.244807e-11
## ZIC2B             1.353809e-13  0.7743060 0.720 0.417 2.333832e-09
## PPRC1             5.976757e-13  0.7833300 0.768 0.536 1.030333e-08
## MYCH              1.244346e-12  0.7901067 0.554 0.262 2.145129e-08
## DLD               4.245796e-10  0.6070986 0.248 0.030 7.319328e-06
## CITED4B           1.069701e-09  0.7000502 0.656 0.423 1.844057e-05
## NET1              2.244380e-09  0.6636208 0.497 0.250 3.869086e-05
## P4HA2             2.245365e-09  0.7749930 0.525 0.292 3.870785e-05
## ADD3B             2.695490e-09  0.5722948 0.528 0.280 4.646755e-05
## SI:CH1073-80I24.3 1.722602e-08  0.4751626 0.896 0.762 2.969593e-04
## ASB11             3.489185e-08  0.5725981 0.817 0.679 6.015006e-04
## AMOTL2A           4.043990e-08  0.5533080 0.325 0.119 6.971435e-04
## PRICKLE1B         4.193558e-08  0.6080152 0.350 0.143 7.229274e-04
## TPBGA             5.734470e-08  0.6045034 0.443 0.226 9.885653e-04
## SPRY2             1.617500e-07  0.4646509 0.356 0.143 2.788408e-03
## MORC3B            3.648701e-07  0.5329174 0.409 0.214 6.289996e-03
## ARL4AB            3.871871e-07  0.5395136 0.357 0.155 6.674718e-03
## HER7              4.194538e-07  0.7405232 0.369 0.173 7.230964e-03
## FOXD3             5.016064e-07  0.5726486 0.203 0.042 8.647192e-03
## MPHOSPH10         5.026233e-07  0.4030817 0.884 0.768 8.664724e-03
## OSR1              5.134896e-07  0.5245960 0.728 0.560 8.852048e-03
## MIXL1             6.557737e-07  0.4752880 0.920 0.792 1.130488e-02
## CITED4A           6.827110e-07  0.4363954 0.229 0.060 1.176925e-02
## ID2A              7.933869e-07  0.6577650 0.248 0.077 1.367720e-02
## ZEB1A             1.008581e-06  0.5394509 0.443 0.250 1.738693e-02
## SEBOX             1.409924e-06  0.5572514 0.566 0.387 2.430568e-02
## EPHA2A            1.437939e-06  0.3005526 0.138 0.006 2.478863e-02
## ETV4              1.821651e-06  0.4664846 0.607 0.446 3.140345e-02
## ADMP              1.937195e-06  1.1245916 0.238 0.077 3.339530e-02
## FOXD5             2.239187e-06  0.5087576 0.345 0.155 3.860134e-02
## APLNRB            2.830886e-06  0.4895863 0.646 0.435 4.880164e-02
## CYCSB             4.027720e-06  0.3477772 0.897 0.780 6.943387e-02
## DACT2             4.709471e-06  0.5168841 0.525 0.333 8.118657e-02
## WNT8A             5.464916e-06  0.4905074 0.503 0.315 9.420968e-02
## DLC               5.557922e-06  0.4649952 0.183 0.042 9.581302e-02
## NOTO              6.707736e-06  1.3380981 0.385 0.214 1.156347e-01
## BCL2L12           6.827857e-06  0.4138374 0.487 0.304 1.177054e-01
## ALDH1A2           9.843983e-06  0.4750650 0.410 0.226 1.697004e-01
## RASL11B           1.056541e-05  0.4099851 0.275 0.107 1.821371e-01
## FGF24             1.411624e-05  0.3539022 0.221 0.071 2.433499e-01
## TCF3B             1.627623e-05  0.4324917 0.643 0.488 2.805860e-01
## IRF2BPL           1.952762e-05  0.3051423 0.133 0.018 3.366366e-01
## TMEM9B            2.199901e-05  0.3652807 0.222 0.077 3.792409e-01
## DDIT4             2.461743e-05  0.3311488 0.909 0.780 4.243799e-01
## GLULB             3.460841e-05  0.3875942 0.831 0.702 5.966143e-01
## SI:CH211-86H15.1  3.517290e-05  0.5212022 0.559 0.411 6.063456e-01
## SERPINH1B         3.827504e-05  0.6387474 0.503 0.333 6.598234e-01
## NRARPA            3.901756e-05  0.3001116 0.137 0.024 6.726237e-01
## PCDH1B            5.809137e-05  0.3668186 0.289 0.143 1.000000e+00
## HER1              7.362482e-05  0.4667488 0.421 0.268 1.000000e+00
## ATP1B1A           8.502019e-05  0.3394173 0.851 0.720 1.000000e+00
## PRICKLE1A         9.524479e-05  0.3926152 0.188 0.065 1.000000e+00
## SIVA1             1.142704e-04  0.2938511 0.768 0.643 1.000000e+00
## PPP1R3B           1.392463e-04  0.3555329 0.485 0.333 1.000000e+00
## TXNIPA            1.618300e-04  0.2733301 0.332 0.179 1.000000e+00
## PUS1              1.781837e-04  0.2618464 0.217 0.089 1.000000e+00
## LFT1              1.837640e-04  0.4853613 0.496 0.363 1.000000e+00
## H1M               2.034278e-04  0.3483280 0.935 0.833 1.000000e+00
## ZGC:55413         2.058967e-04  0.3769076 0.556 0.405 1.000000e+00
## PIM2              2.226180e-04  0.3865050 0.311 0.173 1.000000e+00
## SMARCA5           2.743504e-04  0.3294085 0.696 0.536 1.000000e+00
## HOMEZA            3.776132e-04  0.2906268 0.260 0.131 1.000000e+00
## CHD4A             3.858549e-04  0.3029094 0.863 0.738 1.000000e+00
## CDCA7B            4.011189e-04  0.3196117 0.793 0.667 1.000000e+00
## DZIP1             6.017216e-04  0.2775535 0.277 0.143 1.000000e+00
## SOX13             6.433112e-04  0.2594486 0.154 0.054 1.000000e+00
## TBX6L             6.898617e-04  0.3515496 0.173 0.071 1.000000e+00
## ABCF2A            7.043195e-04  0.3752030 0.516 0.405 1.000000e+00
## HIGD1A            7.096005e-04  0.3598531 0.721 0.595 1.000000e+00
## ZC4H2             7.690717e-04  0.2682001 0.644 0.506 1.000000e+00
## CDC27             7.773218e-04  0.3032994 0.306 0.173 1.000000e+00
## ARL5C             8.561986e-04  0.3961307 0.441 0.327 1.000000e+00
## GDF3              1.032707e-03  0.2875938 0.525 0.387 1.000000e+00
## BTG3              1.183905e-03  0.3676619 0.391 0.286 1.000000e+00
```

```
FeaturePlot (SOX32502, features = c("SOX32_positive" ), pt.size = 2)
```

```
FeaturePlot (SOX32502, features = c("SOX32_negative" ), pt.size = 2)
```
