## Supplementary File 5 for "Nodal signaling establishes a competency window for stochastic cell fate switching": Suplementary file 5.html

Endoderm analysis 60% epiboly (cluster based)


### Endoderm analysis 60% epiboly (cluster based)

###### Luca Guglielmi

#### 14/01/2022

```
# Loading required packages
library( Seurat )
```

```
## Attaching SeuratObject
```

```
library( ggplot2 )
library( dplyr )
```

```
## 
## Attaching package: 'dplyr'
```

```
## The following objects are masked from 'package:stats':
## 
##     filter, lag
```

```
## The following objects are masked from 'package:base':
## 
##     intersect, setdiff, setequal, union
```

```
library( tibble )
library( plotly )
```

```
## 
## Attaching package: 'plotly'
```

```
## The following object is masked from 'package:ggplot2':
## 
##     last_plot
```

```
## The following object is masked from 'package:stats':
## 
##     filter
```

```
## The following object is masked from 'package:graphics':
## 
##     layout
```

```
library( tidyr )
library( cowplot )
library( openxlsx )
library (readxl)
library (knitr)
```

```
# Extracting the 60% epiboly stage from the entire URD Seurat object

srat_obj_rds <- file.path( "~/Documents/URD_projection/urd_srat_obj.rds" )
srat_obj <- readRDS( file = srat_obj_rds )
srat_obj <- SetIdent( object = srat_obj, value = "orig.ident" )
stages <- c( "ZF60" )
zf60_srat_obj <- subset( srat_obj, idents = stages )

# This tidies up the factors and removes the empty factor levels, i.e. the other developmental stages.
zf60_srat_obj@meta.data <- droplevels( zf60_srat_obj@meta.data )
```

```
# Re-processing the 60% epiboly stage into a new Seurat object and PCA/uMAP computing

zf60_srat_obj <- zf60_srat_obj%>% 
    NormalizeData( ) %>%
    FindVariableFeatures( ) %>%
    ScaleData( ) %>%
    RunPCA( )
```

```
## Centering and scaling data matrix
```

```
## PC_ 1 
## Positive:  SOX19A, ALDOB, SOX3, SI:CH211-152C2.3, SI:CH1073-80I24.3, APELA, ALCAMB, POLR3GLA, SOX2, CRABP2B 
##     CXCR4B, ID1, PFKFB4B, GPD1B, ASB11, CABZ01070258.1, MEX3B, HER3, SOX11A, FOXD5 
##     MDKB, HIRIP3, SI:CH211-170D8.2, SI:DKEY-27I16.2, POU5F3, HER8A, PRDM14, SI:DKEY-56M19.5, TFAP2A, TFAP2C 
## Negative:  APLNRA, FSCN1A, TBX16, SNAI1A, FOXA, ISM1, CYP27C1, TUBA8L2, PCDH8, APLNRB 
##     CXCL12B, ELOVL6, MESPAB, MSGN1, KIRREL3L, FLRT3, EFNB2A, CXCL12A, EFNB2B, CST3 
##     LHX1A, ALDH1A2, ASPH, RND1L, CMTM6, RDH10A, RIPPLY1, ZSWIM5, INSB, SASH1A 
## PC_ 2 
## Positive:  HSPB1, CX43.4, TBX16, MSGN1, RBM38, MESPAB, SNAI1A, ANP32E, SP5L, HES6 
##     ALDH1A2, IM:7138239, TA, CYP27C1, WNT8A, DDIT4, CXCL12A, FOXC1A, APLNRB, CXCL12B 
##     VRTN, SIX4A, CDX4, TBX6L, MESPAA, TP53INP2, FSCN1A, PCDH8, HER1, EFNB2B 
## Negative:  KRT8, KRT4, ZGC:101000, KRT5, MID1IP1A, KRT18, ZGC:193505, KRT92, ZNF185, CAPN9 
##     CLDNE, HSD17B14, PONZR5, SI:DKEY-152P16.6, ALOX5B.3, ZGC:162964, KALRNB, PPL, SI:CH211-125O16.4, KRT97 
##     SI:CH211-195B11.3, SI:CH211-202F5.3, KRT23, WASLB, CEBPB, CLDNB, MGLL, ELOVL1A, ABCB5, TRIM16 
## PC_ 3 
## Positive:  GSC, FRZB, FZD8B, SHISA2, NOG1, RRBP1B, FZD8A, SIX3B, OTX1A, SIX7 
##     KLF17, LFT2, OTX1B, ZGC:174153, SI:CH211-163L21.7, FOXA3, CHD, TBX1, CITED4B, ZGC:174855 
##     TMEM120A, RIPPLY1, DHRS3B, ID3, PITX2, RDH8A, B3GNT5B, XBP1, CDC42EP4A, PKDCCA 
## Negative:  CDX4, HES6, WNT8A, RBM38, TA, EVE1, VED, IM:7138239, MSGN1, DLD 
##     WNT11, TBX6L, MESPAB, HER1, ALDH1A2, ANP32E, HER7, SP5L, DDIT4, HELB 
##     VOX, VRTN, TOB1A, TPBGA, SERPINH1B, SNAI1A, FGF24, MESPAA, TBX16, ZNFL2A 
## PC_ 4 
## Positive:  APOEB, APOC1, MARCKSL1B, SOX32, VED, BAMBIA, GATA2A, SOX17, ACKR3B, GATA5 
##     LIMA1A, FAM212AB, VGLL4L, FOXI1, ID3, VOX, PRDX5, S1PR5A, GPM6AB, DLX3B 
##     TFAP2A, SI:CH73-234B20.5, CDH6, DNMT3BB.1, BMP7A, LMO4B, MID1IP1B, KLF2B, CXCR4A, CPN1 
## Negative:  ADMP, NOTO, FOXD3, CHD, FOXB1A, P4HA2, CX43.4, SHHA, RASGEF1BA, ARL4AB 
##     FGF8A, ETV4, TWIST2, ZEB1A, EPHA2A, DACT2, TDGF1, FGF24, TPH1B, EPHA4A 
##     HER3, NTD5, ZIC3, TA, FOXD5, FOXA3, LFT2, NOG1, WNT8A, LFT1 
## PC_ 5 
## Positive:  IRX7, SFRP1A, ZIC3, IRX1B, HER3, ZIC2A, OTX1A, PKDCCB, FZD7A, HESX1 
##     HER11, RDH10A, MYCN, ZIC2B, CPN1, MIDN, LHX5, TWIST1A, CXCL12A, LRATB 
##     SIX4A, HER5, FOXC1A, MEIS1B, ISM1, SHISA2, HAS2, STM, APLNRA, CYP26A1 
## Negative:  CDX4, VED, BAMBIA, EVE1, FZD8B, ZGC:174153, BMP4, KLF17, PRDM1A, ZGC:174855 
##     SIX3B, TA, LFT2, APOC1, GSC, DLX3B, RRBP1B, FRZB, TMEM120A, B3GNT5B 
##     MSX1B, SZL, ATP1B3A, RGCC, FOXI1, SI:CH211-163L21.8, HER7, SI:CH211-163L21.7, SIX7, UBE2E2
```

```
urd_srat_obj2 <- zf60_srat_obj 
ElbowPlot(urd_srat_obj2)
```

```
urd_srat_obj2<-FindNeighbors(urd_srat_obj2, dim = 1:20)
```

```
## Computing nearest neighbor graph
```

```
## Computing SNN
```

```
urd_srat_obj2<-FindClusters(urd_srat_obj2, resolution = 0.05)
```

```
## Modularity Optimizer version 1.3.0 by Ludo Waltman and Nees Jan van Eck
## 
## Number of nodes: 4101
## Number of edges: 140204
## 
## Running Louvain algorithm...
## Maximum modularity in 10 random starts: 0.9620
## Number of communities: 3
## Elapsed time: 0 seconds
```

```
urd_srat_obj2<-RunUMAP(urd_srat_obj2, dim = 1:20 )
```

```
## Warning: The default method for RunUMAP has changed from calling Python UMAP via reticulate to the R-native UWOT using the cosine metric
## To use Python UMAP via reticulate, set umap.method to 'umap-learn' and metric to 'correlation'
## This message will be shown once per session
```

```
## 13:48:15 UMAP embedding parameters a = 0.9922 b = 1.112
```

```
## 13:48:15 Read 4101 rows and found 20 numeric columns
```

```
## 13:48:15 Using Annoy for neighbor search, n_neighbors = 30
```

```
## 13:48:15 Building Annoy index with metric = cosine, n_trees = 50
```

```
## 0%   10   20   30   40   50   60   70   80   90   100%
```

```
## [----|----|----|----|----|----|----|----|----|----|
```

```
## **************************************************|
## 13:48:15 Writing NN index file to temp file /var/folders/s2/phwgmcs926x8kbjk57m5lqw14_t3jy/T//RtmpLNkX55/file15e4c4cb01744
## 13:48:15 Searching Annoy index using 1 thread, search_k = 3000
## 13:48:16 Annoy recall = 100%
## 13:48:16 Commencing smooth kNN distance calibration using 1 thread
## 13:48:16 Initializing from normalized Laplacian + noise
## 13:48:16 Commencing optimization for 500 epochs, with 160084 positive edges
## 13:48:21 Optimization finished
```

```
DimPlot (urd_srat_obj2, reduction = "umap", label = TRUE,  pt.size = 1) + NoLegend()
```

```
# Expression of mixl1 at 50% epiboly
FeaturePlot (urd_srat_obj2, features = c("MIXL1"), pt.size = 2)
```

```
# Sub-setting cells belonging to cluster 1 and 2

SOX32602 <-subset(urd_srat_obj2, idents = c("1","2"), slot = "counts")
SOX32602
```

```
## An object of class Seurat 
## 17239 features across 1520 samples within 1 assay 
## Active assay: RNA (17239 features, 2000 variable features)
##  3 dimensional reductions calculated: pca, tsne, umap
```

```
# Re-processing of cluster 1 and 2 cells and PCA/uMAP computing

SOX32602<- NormalizeData(SOX32602, normalization.method = "LogNormalize", scale.factor = 10000)
```

```
SOX32602<- FindVariableFeatures(SOX32602, selection.method = "vst", nfeatures = 2000)
top10 <- head(VariableFeatures(SOX32602), 10)
plot1 <- VariableFeaturePlot(SOX32602)
plot1 <- LabelPoints(plot = plot1, points = top10, repel = TRUE)
```

```
## When using repel, set xnudge and ynudge to 0 for optimal results
```

```
plot1
```

```
## Warning: Transformation introduced infinite values in continuous x-axis
```

```
## Warning: Removed 1450 rows containing missing values (geom_point).
```

```
all.genes <- rownames(SOX32602)
SOX32602 <- ScaleData(SOX32602, features = all.genes)
```

```
## Centering and scaling data matrix
```

```
SOX32602 <- RunPCA(SOX32602, features = VariableFeatures(object = SOX32602))
```

```
## PC_ 1 
## Positive:  FRZB, SHISA2, GSC, FZD8B, NOG1, FZD8A, ID3, RRBP1B, KLF17, OTX1A 
##     XBP1, OTX1B, SIX7, LFT2, CHD, SIX3B, FOXA, RIPPLY1, FOXA3, DHRS3B 
##     ELOVL6, CST3, PITX2, TBX1, ELL2, ZGC:174153, KRT18, CDC42EP4A, SI:CH211-163L21.7, BTG2 
## Negative:  CDX4, HES6, TA, ANP32E, WNT8A, VED, EVE1, RBM38, ID1, SP5L 
##     DLD, WNT11, HER7, HSPB1, SI:CH211-222L21.1, DDIT4, VOX, NRARPA, IM:7138239, CX43.4 
##     POLR3GLA, APELA, HER1, SERPINH1B, TBX6L, ETV4, UBE2E2, FGF24, ATP1B1A, TPBGA 
## PC_ 2 
## Positive:  NOTO, ADMP, RASGEF1BA, P4HA2, TA, APELA, SOX3, LFT2, FOXD3, CITED4B 
##     NOG1, SHHA, GSC, FZD8B, ARL4AB, TWIST2, ETV4, GADD45BA, CDKN1CA, EPHA2A 
##     RRBP1B, SIX3B, SIX7, CHD, CX43.4, FRZB, KLF17, FOXD5, DACT2, ZGC:174153 
## Negative:  SOX32, GATA5, SOX17, APOEB, CXCL12A, ACKR3B, APOC1, CXCR4A, FAM212AB, ASPH 
##     FSCN1A, CPN1, ACTB2, MARCKSL1B, PRDX5, VOX, CDH6, LIMA1A, FLRT3, GATA2A 
##     IER5L, SFRP1A, IRX7, CMTM6, ZIC2A, RND1L, AKAP12B, TBX16, SI:CH73-234B20.5, S1PR5A 
## PC_ 3 
## Positive:  CABZ01070258.1, CXCR4A, ACKR3B, SOX32, SOX17, SI:CH211-152C2.3, FOXD5, ARL5C, PRDX5, SI:CH73-234B20.5 
##     GPM6AB, SOX3, PLPP3, APELA, CDH6, ASB11, LMO4A, GPD1B, VGLL4L, SOX19A 
##     FOXA2, SOX2, DHRS3B, SSUH2RS1, IQCA1, ARG2, GSTT1B, GATSL2, DNMT3BB.1, CYP2AA8 
## Negative:  MSGN1, APLNRA, ALDH1A2, APLNRB, CYP27C1, EFNB2B, FOXC1A, RDH10A, SIX4A, MESPAA 
##     CXCL12A, ZIC3, CXCL12B, ZIC2A, PCDH8, IM:7138239, EFNB2A, MESPAB, ISM1, LPAR1 
##     GAS1B, MEIS1B, KIRREL3L, MYF5, FOXA, HAS2, ZSWIM5, PCDH10B, RIPPLY1, TUBA8L2 
## PC_ 4 
## Positive:  APOC1, APOEB, ZGC:174153, PRDM1A, ZGC:174855, BAMBIA, VED, KLF17, B3GNT5B, CCND1 
##     EVE1, FZD8B, ATP1B3A, TMEM120A, SI:CH211-163L21.8, ICN, RAB33A, SIX7, RDH8A, PPRC1 
##     OSR1, MLLT1B, BMP4, CHST11, CTSLB, SI:CH211-163L21.7, SI:DKEY-261J4.5, FAM174B, ELL2, FOXP1B 
## Negative:  NOTO, ADMP, FOXA2, CDKN1CA, SHHA, CHD, TWIST2, EPHA4A, INSB, NTD5 
##     FADD, TPH1B, SHHB, HSPB1, RASL11B, ARL4AB, MIDN, BCL2L12, FOXD3, XBP1 
##     EPHA2A, CDH6, PPP1R14C, SP5L, HMMR, ZIC3, FGF8A, FOXA3, SALL1A, IRX7 
## PC_ 5 
## Positive:  MALAT1, HMGN2, P4HA1B, PLOD1A, EVA1BA, HISTH1L, ACTB1, TDH, NID2A, SI:DKEY-85K7.7 
##     ARL4AA, MEIS1B, PCDH10B, HSPB1, ZGC:113208, NTD5, TWIST1A, ACTB2, EGLN2, ZFAND5A 
##     EGLN3, WU:FB55G09, PRR12B, TWIST2, ANGPTL5, GABARAPL1, FN1A, ZGC:158343, SI:CH211-66K16.27, FSTL1B 
## Negative:  RRM2, ZGC:153409, MIXL1, BX324216.1, FGF3, DLEU7, LHX1A, NCL, HIST2H2AB, EFNB2A 
##     POU5F3, SNAI1A, ZNFL2A, TBX16, CTH1, CCNG1, MARCKSL1B, RND1L, H1M, TRIB3 
##     PITX2, FP102169.1, SI:CH211-113A14.18, MAP1LC3B, HSP90AA1.2, DKK1B, ELOVL6, SI:CH1073-80I24.3, WNT11, DDX4
```

```
DimPlot(SOX32602, reduction = "pca")
```

```
VizDimLoadings(SOX32602, dims = 1:2, reduction = "pca")
```

```
ElbowPlot(SOX32602)
```

```
SOX32602<- FindNeighbors(SOX32602, dims = 1:15)
```

```
## Computing nearest neighbor graph
```

```
## Computing SNN
```

```
SOX32602 <- FindClusters(SOX32602, resolution = 0.01)
```

```
## Modularity Optimizer version 1.3.0 by Ludo Waltman and Nees Jan van Eck
## 
## Number of nodes: 1520
## Number of edges: 50945
## 
## Running Louvain algorithm...
## Maximum modularity in 10 random starts: 0.9900
## Number of communities: 1
## Elapsed time: 0 seconds
```

```
SOX32602 <- RunUMAP(SOX32602, dims = 1:15)
```

```
## 13:48:30 UMAP embedding parameters a = 0.9922 b = 1.112
```

```
## 13:48:30 Read 1520 rows and found 15 numeric columns
```

```
## 13:48:30 Using Annoy for neighbor search, n_neighbors = 30
```

```
## 13:48:30 Building Annoy index with metric = cosine, n_trees = 50
```

```
## 0%   10   20   30   40   50   60   70   80   90   100%
```

```
## [----|----|----|----|----|----|----|----|----|----|
```

```
## **************************************************|
## 13:48:31 Writing NN index file to temp file /var/folders/s2/phwgmcs926x8kbjk57m5lqw14_t3jy/T//RtmpLNkX55/file15e4c7ff950b
## 13:48:31 Searching Annoy index using 1 thread, search_k = 3000
## 13:48:31 Annoy recall = 100%
## 13:48:31 Commencing smooth kNN distance calibration using 1 thread
## 13:48:31 Initializing from normalized Laplacian + noise
## 13:48:31 Commencing optimization for 500 epochs, with 56392 positive edges
## 13:48:33 Optimization finished
```

```
# Expression of mesoderm, endoderm and ectoderm markers in gata5 positive cells

FeaturePlot(SOX32602, features = c("SOX32"), pt.size = 2)
```

```
FeaturePlot(SOX32602, features = c("GATA5"), pt.size = 2)
```

```
FeaturePlot(SOX32602, features = c("TA"), pt.size = 2)
```

```
FeaturePlot(SOX32602, features = c("SOX2"), pt.size = 2)
```

```
# clustering of gata5 positive cells with increasing granularity

SOX32602<- FindNeighbors(SOX32602, dims = 1:15)
```

```
## Computing nearest neighbor graph
```

```
## Computing SNN
```

```
SOX32602 <- FindClusters(SOX32602, resolution = 0.01)
```

```
## Modularity Optimizer version 1.3.0 by Ludo Waltman and Nees Jan van Eck
## 
## Number of nodes: 1520
## Number of edges: 50945
## 
## Running Louvain algorithm...
## Maximum modularity in 10 random starts: 0.9900
## Number of communities: 1
## Elapsed time: 0 seconds
```

```
SOX32602 <- RunUMAP(SOX32602, dims = 1:15)
```

```
## 13:48:35 UMAP embedding parameters a = 0.9922 b = 1.112
```

```
## 13:48:35 Read 1520 rows and found 15 numeric columns
```

```
## 13:48:35 Using Annoy for neighbor search, n_neighbors = 30
```

```
## 13:48:35 Building Annoy index with metric = cosine, n_trees = 50
```

```
## 0%   10   20   30   40   50   60   70   80   90   100%
```

```
## [----|----|----|----|----|----|----|----|----|----|
```

```
## **************************************************|
## 13:48:35 Writing NN index file to temp file /var/folders/s2/phwgmcs926x8kbjk57m5lqw14_t3jy/T//RtmpLNkX55/file15e4c30b0e44f
## 13:48:35 Searching Annoy index using 1 thread, search_k = 3000
## 13:48:36 Annoy recall = 100%
## 13:48:36 Commencing smooth kNN distance calibration using 1 thread
## 13:48:36 Initializing from normalized Laplacian + noise
## 13:48:36 Commencing optimization for 500 epochs, with 56392 positive edges
## 13:48:38 Optimization finished
```

```
DimPlot(SOX32602, reduction = "umap",label = TRUE,  pt.size = 2) + NoLegend()
```

```
# clustering of gata5 positive cells with increasing granularity

SOX32602<- FindNeighbors(SOX32602, dims = 1:15)
```

```
## Computing nearest neighbor graph
```

```
## Computing SNN
```

```
SOX32602 <- FindClusters(SOX32602, resolution = 0.02)
```

```
## Modularity Optimizer version 1.3.0 by Ludo Waltman and Nees Jan van Eck
## 
## Number of nodes: 1520
## Number of edges: 50945
## 
## Running Louvain algorithm...
## Maximum modularity in 10 random starts: 0.9819
## Number of communities: 2
## Elapsed time: 0 seconds
```

```
SOX32602 <- RunUMAP(SOX32602, dims = 1:15)
```

```
## 13:48:39 UMAP embedding parameters a = 0.9922 b = 1.112
```

```
## 13:48:39 Read 1520 rows and found 15 numeric columns
```

```
## 13:48:39 Using Annoy for neighbor search, n_neighbors = 30
```

```
## 13:48:39 Building Annoy index with metric = cosine, n_trees = 50
```

```
## 0%   10   20   30   40   50   60   70   80   90   100%
```

```
## [----|----|----|----|----|----|----|----|----|----|
```

```
## **************************************************|
## 13:48:39 Writing NN index file to temp file /var/folders/s2/phwgmcs926x8kbjk57m5lqw14_t3jy/T//RtmpLNkX55/file15e4c3ab26f12
## 13:48:39 Searching Annoy index using 1 thread, search_k = 3000
## 13:48:39 Annoy recall = 100%
## 13:48:39 Commencing smooth kNN distance calibration using 1 thread
## 13:48:40 Initializing from normalized Laplacian + noise
## 13:48:40 Commencing optimization for 500 epochs, with 56392 positive edges
## 13:48:42 Optimization finished
```

```
DimPlot(SOX32602, reduction = "umap",label = TRUE,  pt.size = 2) + NoLegend()
```

```
SOX32602<- FindNeighbors(SOX32602, dims = 1:15)
```

```
## Computing nearest neighbor graph
```

```
## Computing SNN
```

```
SOX32602 <- FindClusters(SOX32602, resolution = 0.1)
```

```
## Modularity Optimizer version 1.3.0 by Ludo Waltman and Nees Jan van Eck
## 
## Number of nodes: 1520
## Number of edges: 50945
## 
## Running Louvain algorithm...
## Maximum modularity in 10 random starts: 0.9335
## Number of communities: 3
## Elapsed time: 0 seconds
```

```
SOX32602 <- RunUMAP(SOX32602, dims = 1:15)
```

```
## 13:48:42 UMAP embedding parameters a = 0.9922 b = 1.112
```

```
## 13:48:42 Read 1520 rows and found 15 numeric columns
```

```
## 13:48:42 Using Annoy for neighbor search, n_neighbors = 30
```

```
## 13:48:42 Building Annoy index with metric = cosine, n_trees = 50
```

```
## 0%   10   20   30   40   50   60   70   80   90   100%
```

```
## [----|----|----|----|----|----|----|----|----|----|
```

```
## **************************************************|
## 13:48:42 Writing NN index file to temp file /var/folders/s2/phwgmcs926x8kbjk57m5lqw14_t3jy/T//RtmpLNkX55/file15e4c2fe92f2d
## 13:48:42 Searching Annoy index using 1 thread, search_k = 3000
## 13:48:43 Annoy recall = 100%
## 13:48:43 Commencing smooth kNN distance calibration using 1 thread
## 13:48:43 Initializing from normalized Laplacian + noise
## 13:48:43 Commencing optimization for 500 epochs, with 56392 positive edges
## 13:48:45 Optimization finished
```

```
DimPlot(SOX32602, reduction = "umap",label = TRUE,  pt.size = 2) + NoLegend()
```

```
SOX32602<- FindNeighbors(SOX32602, dims = 1:15)
```

```
## Computing nearest neighbor graph
```

```
## Computing SNN
```

```
SOX32602 <- FindClusters(SOX32602, resolution = 0.15)
```

```
## Modularity Optimizer version 1.3.0 by Ludo Waltman and Nees Jan van Eck
## 
## Number of nodes: 1520
## Number of edges: 50945
## 
## Running Louvain algorithm...
## Maximum modularity in 10 random starts: 0.9110
## Number of communities: 4
## Elapsed time: 0 seconds
```

```
SOX32602 <- RunUMAP(SOX32602, dims = 1:15)
```

```
## 13:48:46 UMAP embedding parameters a = 0.9922 b = 1.112
```

```
## 13:48:46 Read 1520 rows and found 15 numeric columns
```

```
## 13:48:46 Using Annoy for neighbor search, n_neighbors = 30
```

```
## 13:48:46 Building Annoy index with metric = cosine, n_trees = 50
```

```
## 0%   10   20   30   40   50   60   70   80   90   100%
```

```
## [----|----|----|----|----|----|----|----|----|----|
```

```
## **************************************************|
## 13:48:46 Writing NN index file to temp file /var/folders/s2/phwgmcs926x8kbjk57m5lqw14_t3jy/T//RtmpLNkX55/file15e4c3520f12d
## 13:48:46 Searching Annoy index using 1 thread, search_k = 3000
## 13:48:46 Annoy recall = 100%
## 13:48:46 Commencing smooth kNN distance calibration using 1 thread
## 13:48:47 Initializing from normalized Laplacian + noise
## 13:48:47 Commencing optimization for 500 epochs, with 56392 positive edges
## 13:48:49 Optimization finished
```

```
DimPlot(SOX32602, reduction = "umap",label = TRUE,  pt.size = 2) + NoLegend()
```

```
SOX32602<- FindNeighbors(SOX32602, dims = 1:15)
```

```
## Computing nearest neighbor graph
```

```
## Computing SNN
```

```
SOX32602 <- FindClusters(SOX32602, resolution = 0.2)
```

```
## Modularity Optimizer version 1.3.0 by Ludo Waltman and Nees Jan van Eck
## 
## Number of nodes: 1520
## Number of edges: 50945
## 
## Running Louvain algorithm...
## Maximum modularity in 10 random starts: 0.8950
## Number of communities: 5
## Elapsed time: 0 seconds
```

```
SOX32602 <- RunUMAP(SOX32602, dims = 1:15)
```

```
## 13:48:49 UMAP embedding parameters a = 0.9922 b = 1.112
```

```
## 13:48:49 Read 1520 rows and found 15 numeric columns
```

```
## 13:48:49 Using Annoy for neighbor search, n_neighbors = 30
```

```
## 13:48:49 Building Annoy index with metric = cosine, n_trees = 50
```

```
## 0%   10   20   30   40   50   60   70   80   90   100%
```

```
## [----|----|----|----|----|----|----|----|----|----|
```

```
## **************************************************|
## 13:48:50 Writing NN index file to temp file /var/folders/s2/phwgmcs926x8kbjk57m5lqw14_t3jy/T//RtmpLNkX55/file15e4c6d42470c
## 13:48:50 Searching Annoy index using 1 thread, search_k = 3000
## 13:48:50 Annoy recall = 100%
## 13:48:50 Commencing smooth kNN distance calibration using 1 thread
## 13:48:50 Initializing from normalized Laplacian + noise
## 13:48:50 Commencing optimization for 500 epochs, with 56392 positive edges
## 13:48:52 Optimization finished
```

```
DimPlot(SOX32602, reduction = "umap",label = TRUE,  pt.size = 2) + NoLegend()
```

```
#Comparing sox32 positive/ negative cells 


SOX32602 <- AddMetaData( object = SOX32602,
                       metadata = GetAssayData( SOX32602, slot = "counts" )[ "SOX32", ] > 0,
                       col.name = "SOX32_positive" )
SOX32602 <- SetIdent( object = SOX32602, value = "SOX32_positive" )
SOX32.markers <- FindMarkers( SOX32602, ident.2 = "FALSE", ident.1 = "TRUE", min.diff.pct = 0.1, only.pos = TRUE )

head(SOX32.markers,n = 80)
```

```
##                           p_val avg_log2FC pct.1 pct.2     p_val_adj
## SOX32              0.000000e+00  4.8341975 1.000 0.000  0.000000e+00
## SOX17             6.008658e-274  2.8620429 0.859 0.002 1.035833e-269
## ACKR3B            4.188287e-205  2.8799652 0.843 0.043 7.220188e-201
## CXCR4A            4.341631e-181  3.9460890 0.927 0.108 7.484538e-177
## SI:CH73-234B20.5  8.470194e-130  1.2459664 0.476 0.008 1.460177e-125
## PRDX5             8.920162e-123  2.5209888 0.723 0.086 1.537747e-118
## GPM6AB            4.231691e-121  1.5406837 0.623 0.044 7.295012e-117
## CDH6              1.422802e-106  2.2378060 0.764 0.131 2.452769e-102
## PLPP3             5.937047e-103  1.1845705 0.524 0.035  1.023487e-98
## VGLL4L             2.111851e-96  1.6459737 0.529 0.044  3.640621e-92
## FOXA2              7.413711e-93  1.9158445 0.869 0.195  1.278050e-88
## CABZ01070258.1     1.805606e-88  2.0973207 0.874 0.280  3.112685e-84
## S1PR5A             2.089275e-77  0.9547645 0.476 0.047  3.601701e-73
## GATA5              1.930731e-76  1.5680483 0.880 0.273  3.328388e-72
## SSUH2RS1           1.725255e-70  0.5765721 0.288 0.008  2.974168e-66
## DHRS3B             1.521729e-67  1.8214266 0.874 0.325  2.623309e-63
## IQCA1              1.022641e-66  0.5583330 0.236 0.002  1.762930e-62
## GSTT1B             8.472432e-66  1.1962014 0.482 0.066  1.460563e-61
## GATSL2             6.046004e-64  1.3621421 0.649 0.157  1.042271e-59
## FLRT3              5.801458e-54  1.3116664 0.937 0.556  1.000113e-49
## ST6GALNAC          2.005714e-48  0.5745459 0.246 0.015  3.457651e-44
## CPN1               2.861984e-47  1.1524273 0.696 0.211  4.933774e-43
## LMO4B              5.065150e-44  0.6958264 0.309 0.035  8.731812e-40
## SLC43A2B           7.070120e-44  0.7139410 0.230 0.015  1.218818e-39
## LHFP               8.860896e-44  0.7180132 0.304 0.034  1.527530e-39
## ID3                2.091423e-41  1.2245503 0.921 0.489  3.605403e-37
## FKBP7              8.838172e-39  1.3217290 0.738 0.359  1.523612e-34
## NT5C3A             1.673365e-38  0.4891534 0.157 0.005  2.884714e-34
## NT5C2B             2.094491e-38  0.7401639 0.230 0.020  3.610693e-34
## ASPH               1.101402e-36  0.9504073 0.984 0.735  1.898707e-32
## ABTB2              1.218207e-36  0.3289206 0.136 0.002  2.100067e-32
## JPH3               1.264310e-35  0.2505201 0.115 0.000  2.179543e-31
## TMEM223            2.803317e-32  0.5903609 0.330 0.062  4.832639e-28
## ANGPTL7            4.426207e-32  0.4353854 0.115 0.002  7.630337e-28
## TSPAN18A           5.595294e-32  0.2889390 0.120 0.002  9.645728e-28
## CXXC5B             5.810980e-32  0.6951079 0.445 0.114  1.001755e-27
## FAM212AB           7.786703e-32  1.0012809 0.785 0.391  1.342350e-27
## TP53I11A           2.330711e-31  0.4851323 0.251 0.035  4.017912e-27
## ELOVL6             4.282119e-31  0.8873273 0.770 0.330  7.381945e-27
## HER5               5.026772e-31  0.9701495 0.304 0.055  8.665652e-27
## CCDC39             1.411926e-30  0.2845062 0.120 0.003  2.434019e-26
## GRAMD3             1.531919e-30  0.9497051 0.712 0.328  2.640874e-26
## COMMD7             1.984824e-30  0.7136509 0.529 0.172  3.421639e-26
## ARL5C              8.565950e-30  0.9532271 0.628 0.255  1.476684e-25
## CMTM7              1.604626e-29  0.9152846 0.518 0.175  2.766215e-25
## FSCN1A             5.525503e-29  0.7391764 0.984 0.882  9.525415e-25
## CYP2AA8            1.018827e-28  1.1999701 0.497 0.180  1.756356e-24
## LIMA1A             1.737137e-28  0.9663856 0.634 0.273  2.994651e-24
## LFNG               1.228421e-26  0.7499353 0.539 0.196  2.117675e-22
## TRIP10A            4.636259e-26  0.2575163 0.126 0.007  7.992447e-22
## GPD1B              7.669563e-26  0.7293517 0.325 0.078  1.322156e-21
## SFRP1A             1.084391e-25  1.0146983 0.796 0.461  1.869382e-21
## TSKU               1.301219e-25  0.7348832 0.607 0.250  2.243172e-21
## RBMS2B             3.454352e-25  0.4215561 0.246 0.043  5.954957e-21
## AHI1               6.455146e-25  1.0107785 0.613 0.285  1.112803e-20
## KRT18              2.379019e-24  0.8976116 0.471 0.160  4.101191e-20
## HMGCS1             6.010877e-24  0.3325464 0.173 0.020  1.036215e-19
## GATA2A             1.637999e-23  0.7734142 0.372 0.109  2.823746e-19
## MRPL15             2.813014e-23  0.6353886 0.403 0.129  4.849356e-19
## DNMT3BB.1          4.330204e-23  0.7294870 0.346 0.098  7.464839e-19
## ARL6IP5A           6.313091e-23  0.6699101 0.330 0.090  1.088314e-18
## HER11              6.450099e-23  0.6798824 0.330 0.084  1.111933e-18
## SI:CH211-209J10.5  5.389140e-22  0.6113073 0.382 0.121  9.290338e-18
## GNAIA              1.313113e-21  0.7626429 0.880 0.637  2.263675e-17
## PREX1              5.236396e-21  0.3198522 0.162 0.021  9.027024e-17
## PHLDB1B            1.248969e-20  0.6511156 0.403 0.143  2.153097e-16
## CHAC1              1.356644e-20  0.8013247 0.560 0.272  2.338719e-16
## LMO4A              1.612634e-20  0.9724957 0.576 0.274  2.780020e-16
## AOC2               1.717858e-20  0.6635086 0.476 0.193  2.961415e-16
## SLC25A36B          1.887664e-20  0.4456135 0.283 0.071  3.254145e-16
## SP5A               5.212494e-20  0.7540196 0.759 0.454  8.985818e-16
## SI:CH211-155E24.3  5.218430e-20  0.6787180 0.529 0.236  8.996051e-16
## TPD52L2A           8.548861e-20  0.4551744 0.293 0.080  1.473738e-15
## IER5L              8.662638e-20  0.6750990 0.592 0.280  1.493352e-15
## CSRP2              1.910757e-19  0.5342976 0.183 0.032  3.293954e-15
## AHCYL2             2.187465e-19  0.7050455 0.654 0.369  3.770972e-15
## RAP2B              2.200700e-19  0.7405625 0.634 0.337  3.793786e-15
## TUBB5              2.502498e-19  0.7288465 0.414 0.160  4.314056e-15
## ZGC:113372         3.030721e-19  0.6538905 0.487 0.201  5.224660e-15
## TBX1               4.962744e-19  0.5172183 0.366 0.117  8.555274e-15
```

```
SOX32602 <- AddMetaData( object = SOX32602,
                       metadata = GetAssayData( SOX32602, slot = "counts" )[ "SOX32", ] < 1,
                       col.name = "SOX32_negative" )
SOX32602 <- SetIdent( object = SOX32602, value = "SOX32_negative" )
SOX32.markers <- FindMarkers( SOX32602, ident.2 = "FALSE", ident.1 = "TRUE", min.diff.pct = 0.1, only.pos = TRUE )

head(SOX32.markers,n = 80)
```

```
##                          p_val avg_log2FC pct.1 pct.2    p_val_adj
## WNT8A             2.405083e-35  1.6641485 0.631 0.136 4.146123e-31
## RBM38             8.529271e-35  1.6764787 0.669 0.225 1.470361e-30
## DDIT4             7.039740e-32  1.3334953 0.771 0.440 1.213581e-27
## TA                4.014448e-31  1.9649048 0.787 0.576 6.920508e-27
## ADD3B             7.044270e-27  1.0304063 0.758 0.382 1.214362e-22
## P4HA2             9.253705e-27  1.3137604 0.576 0.152 1.595246e-22
## SERPINH1B         1.431402e-26  1.5887751 0.582 0.188 2.467594e-22
## ALDH1A2           8.278532e-26  1.5204682 0.550 0.152 1.427136e-21
## CDX4              6.919955e-25  1.7896234 0.710 0.398 1.192931e-20
## APLNRB            2.436607e-23  1.2700267 0.546 0.183 4.200467e-19
## MYCH              2.366786e-22  0.9861619 0.728 0.471 4.080102e-18
## TOB1A             1.189717e-21  1.2092891 0.427 0.068 2.050954e-17
## HER1              1.902084e-21  1.2907234 0.431 0.068 3.279002e-17
## IM:7138239        2.261991e-21  1.3028390 0.611 0.293 3.899446e-17
## ATP1B1A           5.442307e-21  0.9293851 0.877 0.738 9.381993e-17
## MSGN1             2.553642e-20  2.2669515 0.599 0.351 4.402224e-16
## HER7              2.631173e-20  1.4553311 0.447 0.094 4.535879e-16
## FOXC1A            4.158223e-20  1.2237880 0.412 0.063 7.168361e-16
## TBX6L             6.912518e-19  0.9872913 0.350 0.026 1.191649e-14
## PPRC1             1.726825e-18  0.8987292 0.691 0.408 2.976873e-14
## DLD               1.662575e-17  1.0428223 0.375 0.063 2.866114e-13
## LFT1              6.056648e-17  1.0188485 0.417 0.099 1.044105e-12
## POLR3GLA          1.318144e-16  0.8104645 0.845 0.702 2.272348e-12
## CYP27C1           1.428897e-16  0.9511109 0.655 0.403 2.463275e-12
## RASGEF1BA         1.832539e-16  1.0582003 0.338 0.047 3.159114e-12
## DLC               8.232396e-16  0.9478629 0.298 0.021 1.419183e-11
## MESPAA            1.189932e-15  1.2833973 0.339 0.052 2.051323e-11
## DACT2             1.259917e-15  0.9098597 0.496 0.204 2.171971e-11
## ETV4              2.516477e-15  0.8784111 0.629 0.424 4.338155e-11
## SKILB             6.275951e-15  0.8070857 0.376 0.094 1.081911e-10
## SURF6             1.473000e-14  0.7138053 0.781 0.623 2.539304e-10
## FGF24             5.100791e-14  0.8439301 0.287 0.037 8.793254e-10
## APLNRA            1.647613e-13  0.9713134 0.630 0.440 2.840320e-09
## FGF8A             3.394888e-13  0.9409320 0.388 0.126 5.852447e-09
## OSR1              1.049884e-12  0.9676722 0.406 0.168 1.809896e-08
## SOX3              2.399471e-12  1.2040576 0.273 0.042 4.136448e-08
## RRS1              6.014605e-12  0.6158605 0.796 0.665 1.036858e-07
## CITED4B           1.073026e-11  0.9561082 0.657 0.424 1.849789e-07
## EFNB2B            1.085975e-11  0.8931144 0.535 0.346 1.872112e-07
## APELA             2.673044e-11  1.0804408 0.366 0.141 4.608061e-07
## RHOV              6.949140e-11  0.6410164 0.574 0.356 1.197962e-06
## ADMP              1.436984e-10  1.1801585 0.252 0.047 2.477216e-06
## AMOTL2A           1.909151e-10  0.5897149 0.245 0.047 3.291185e-06
## WNT5B             4.764914e-10  0.6072002 0.284 0.079 8.214236e-06
## CXCL12B           5.916044e-10  0.8124344 0.540 0.346 1.019867e-05
## WNT11             7.207976e-10  0.9036281 0.637 0.513 1.242583e-05
## ZEB1A             1.051687e-09  0.6615726 0.436 0.230 1.813003e-05
## ZIC3              1.353800e-09  0.8342387 0.391 0.188 2.333817e-05
## CRABP2B           1.820467e-09  0.7419540 0.473 0.262 3.138303e-05
## SOX2              2.093878e-09  0.7035797 0.189 0.016 3.609635e-05
## CYP26A1           2.399555e-09  0.7929947 0.441 0.236 4.136593e-05
## FOXD3             2.557815e-09  1.0326105 0.178 0.010 4.409418e-05
## SI:DKEY-261H17.1  2.869193e-09  0.5231373 0.211 0.031 4.946202e-05
## UBE2E2            3.418866e-09  0.6627095 0.332 0.131 5.893784e-05
## ID1               3.829448e-09  0.6538591 0.808 0.665 6.601585e-05
## NRARPA            4.144395e-09  0.6171029 0.311 0.110 7.144523e-05
## PPP1R3CA          4.795755e-09  0.6665053 0.481 0.288 8.267403e-05
## SIX4A             6.214121e-09  0.6937614 0.386 0.194 1.071252e-04
## ZNF503            6.972767e-09  0.4890564 0.203 0.031 1.202035e-04
## PCDH1B            1.136134e-08  0.5385834 0.312 0.115 1.958581e-04
## RRP1              1.359274e-08  0.5241332 0.754 0.644 2.343252e-04
## ID2A              2.295791e-08  0.6583274 0.266 0.084 3.957714e-04
## ZNF703            3.600265e-08  0.4711508 0.214 0.047 6.206497e-04
## TCF3B             4.858109e-08  0.6097003 0.603 0.487 8.374894e-04
## PPP1R14C          5.355765e-08  0.5119843 0.210 0.047 9.232803e-04
## ITGB5             5.656227e-08  0.4836214 0.251 0.079 9.750770e-04
## SI:CH1073-80I24.3 7.240478e-08  0.6628028 0.489 0.314 1.248186e-03
## GAS1B             8.199811e-08  0.6648997 0.357 0.178 1.413565e-03
## IRAK3             8.627087e-08  0.4805346 0.183 0.031 1.487224e-03
## SOX19A            1.037717e-07  0.6442338 0.206 0.047 1.788921e-03
## SI:DKEY-239H2.3   1.394145e-07  0.4504008 0.758 0.654 2.403367e-03
## TDH               1.507628e-07  0.5978670 0.321 0.141 2.599000e-03
## CITED4A           1.549132e-07  0.4366438 0.220 0.058 2.670548e-03
## RIPPLY2           1.753476e-07  0.4900660 0.163 0.021 3.022818e-03
## LEF1              2.493072e-07  0.4784850 0.295 0.126 4.297806e-03
## TIMM50            2.636823e-07  0.3828794 0.223 0.063 4.545619e-03
## NT5DC2            3.913261e-07  0.4962515 0.336 0.162 6.746071e-03
## HN1L              5.722389e-07  0.4645022 0.622 0.482 9.864826e-03
## BCL2L12           6.102379e-07  0.5605066 0.446 0.283 1.051989e-02
## TWISTNB           9.585960e-07  0.5437339 0.623 0.513 1.652524e-02
```

```
FeaturePlot (SOX32602, features = c("SOX32_positive" ), pt.size = 2)
```

```
FeaturePlot (SOX32602, features = c("SOX32_negative" ), pt.size = 2)
```
