## Supplementary File 1 for "Nodal signaling establishes a competency window for stochastic cell fate switching": Supplementary file 1.html

Cell cycle analysis sox32


### Cell cycle analysis sox32

###### Luca Guglielmi

#### 12/01/2022

```
# Loading required packages

library(FactoMineR)
library( Seurat )
```

```
## Attaching SeuratObject
```

```
library( ggplot2 )
library( dplyr )
```

```
## 
## Attaching package: 'dplyr'
```

```
## The following objects are masked from 'package:stats':
## 
##     filter, lag
```

```
## The following objects are masked from 'package:base':
## 
##     intersect, setdiff, setequal, union
```

```
library( tibble )
library( plotly )
```

```
## 
## Attaching package: 'plotly'
```

```
## The following object is masked from 'package:ggplot2':
## 
##     last_plot
```

```
## The following object is masked from 'package:stats':
## 
##     filter
```

```
## The following object is masked from 'package:graphics':
## 
##     layout
```

```
library( tidyr )
library( cowplot )
library( reshape2 )
```

```
## 
## Attaching package: 'reshape2'
```

```
## The following object is masked from 'package:tidyr':
## 
##     smiths
```

```
library( openxlsx )
library (readxl)
library (knitr)
library(tidyverse)
```

```
## Registered S3 method overwritten by 'cli':
##   method     from         
##   print.boxx spatstat.geom
```

```
## ── Attaching packages ─────────────────────────────────────── tidyverse 1.3.1 ──
```

```
## ✓ readr   2.1.0     ✓ stringr 1.4.0
## ✓ purrr   0.3.4     ✓ forcats 0.5.1
```

```
## ── Conflicts ────────────────────────────────────────── tidyverse_conflicts() ──
## x plotly::filter() masks dplyr::filter(), stats::filter()
## x dplyr::lag()     masks stats::lag()
```

```
# Extracting the 60% epiboly stage from the entire URD Seurat object

srat_obj_rds <- file.path("~/Documents/URD_projection/urd_srat_obj.rds")
srat_obj <- readRDS( file = srat_obj_rds )
srat_obj <- SetIdent( object = srat_obj, value = "orig.ident" )
stages <- c( "ZF50" )
zf50_srat_obj <- subset( srat_obj, idents = stages )

# This tidies up the factors and removes the empty factor levels, i.e. the other developmental stages.

zf50_srat_obj@meta.data <- droplevels( zf50_srat_obj@meta.data )
```

```
## 13:31:39 UMAP embedding parameters a = 0.9922 b = 1.112
```

```
## 13:31:39 Read 5716 rows and found 15 numeric columns
```

```
## 13:31:39 Using Annoy for neighbor search, n_neighbors = 30
```

```
## 13:31:39 Building Annoy index with metric = cosine, n_trees = 50
```

```
## 0%   10   20   30   40   50   60   70   80   90   100%
```

```
## [----|----|----|----|----|----|----|----|----|----|
```

```
## **************************************************|
## 13:31:39 Writing NN index file to temp file /var/folders/s2/phwgmcs926x8kbjk57m5lqw14_t3jy/T//RtmpquxPVo/file15ca54cbf0ce6
## 13:31:39 Searching Annoy index using 1 thread, search_k = 3000
## 13:31:41 Annoy recall = 100%
## 13:31:41 Commencing smooth kNN distance calibration using 1 thread
## 13:31:41 Initializing from normalized Laplacian + noise
## 13:31:41 Commencing optimization for 500 epochs, with 225422 positive edges
## 13:31:48 Optimization finished
```

```
DimPlot (urd_srat_obj, reduction = "umap", label = TRUE,  pt.size = 1) + NoLegend()
```

```
# Sub-setting gata5 positive cells into a new Seurat object
SOX3250C <- subset(urd_srat_obj, subset = GATA5 > 0, slot = "counts" )
SOX3250C
```

```
## An object of class Seurat 
## 17239 features across 461 samples within 1 assay 
## Active assay: RNA (17239 features, 2000 variable features)
##  3 dimensional reductions calculated: pca, tsne, umap
```

```
# Re-processing of gata5 positive cells and PCA/uMAP computing

SOX3250C<- NormalizeData(SOX3250C, normalization.method = "LogNormalize", scale.factor = 10000)
SOX3250C<- FindVariableFeatures(SOX3250C, selection.method = "vst", nfeatures = 2000)
all.genes <- rownames(SOX3250C)
SOX3250C <- ScaleData(SOX3250C, features = all.genes)
```

```
## Centering and scaling data matrix
```

```
SOX3250C <- RunPCA(SOX3250C, features = VariableFeatures(object = SOX3250C))
```

```
## PC_ 1 
## Positive:  KRT4, CAPN9, KRT92, KRT97, SI:CH211-125O16.4, KALRNB, ZGC:174935, ZGC:91849, EVPLA, KRT5 
##     SLC3A2B, PONZR5, HSD17B14, DSPA, SI:CH211-269I23.2, CLDNF, ZGC:193505, MGLL, SI:DKEYP-67A8.4, GRHL3 
##     SLC2A12, SI:CH211-69B7.6, CLDNB, EPPK1, CNTF, RASSF7B, SID1, CABZ01073832.1, STYK1, SI:DKEY-17E16.17 
## Negative:  TBX16, APOEB, ALDOB, HSPB1, APOC1, NNR, MSGN1, SI:DKEY-68O6.5, CCNA2, NOP2 
##     ZNFL2A, MESPAB, SI:CH211-152C2.3, BAMBIA, CX43.4, HES6, CCNB1, DDIT4, SI:DKEY-56M19.5, CTH1 
##     ANP32E, FP102169.1, MESPAA, EVE1, FSCN1A, APLNRB, SI:DKEY-261J4.5, IM:7138239, OSR1, ALDH1A2 
## PC_ 2 
## Positive:  GSC, CHD, NOG1, FRZB, FZD8B, RIPPLY1, OTX1A, FZD8A, KLF17, ID3 
##     SLC25A33, OTX1B, SIX3B, PKDCCA, FOXA3, FOXA2, RND1L, STM, SERTAD2B, XBP1 
##     MAGI1B, ISM1, SHISA2, GADD45BA, CST3, FOXA, LFT2, TPH1B, HER11, TBX1 
## Negative:  VED, HES6, WNT11, EVE1, TA, MESPAB, CDX4, VOX, WNT8A, BAMBIA 
##     MYCH, ID1, FP102169.1, TBX16, SI:DKEY-261J4.5, ID2A, CDCA7A, MSGN1, SI:DKEY-56M19.5, TPBGA 
##     BMP2B, ALDH1A2, HER7, POLR3GLA, DLD, VENT, PPRC1, ANP32E, SI:DKEY-27I16.2, MESPAA 
## PC_ 3 
## Positive:  SOX32, APOC1, SP5L, SOX17, HMGN2, ACKR3B, CXCR4A, SP5A, MALAT1, LMO4B 
##     SFRP1A, APOEB, FAM212AB, CDH6, FMNL2B, LMO1, CD82A, ATP1B3A, LAMB1A, RND1L 
##     IM:7138239, CDH2, SI:CH211-133N4.4, CPN1, PRDX5, IPCEF1, MSGN1, DHRS3B, SLC13A4, SZL 
## Negative:  H1M, RCOR1, RASGEF1BA, ARL4AB, MKRN4, GTF2A1, ZGC:173742, APELA, MIXL1, CITED4B 
##     SI:CH1073-80I24.3, PRICKLE1B, REEP2, ZGC:113886, SEBOX, BZW1B, OSR1, DCTPP1, SRFBP1, ACP5A 
##     ZIC2B, YWHAQA, CDH1, FGF8A, UCK2A, DNAJB1B, XBP1, CLDND, PPRC1, SI:CH211-86H15.1 
## PC_ 4 
## Positive:  SI:CH211-152C2.3, ZIC2B, GLULB, CCNA2, CITED4B, SI:CH1073-80I24.3, FOPNL, APELA, SOX19A, DYNLL1 
##     RPLP0, DDX5, CCNB1, GSC, ACTB1, OSR1, DNAJB6B, SOX3, ETS2, ID1 
##     ACTB2, CHD, CTH1, FZD8B, H2AFX, SI:DKEY-105I19.5, SPATA6L, NOTO, MEX3B, TPH1B 
## Negative:  AKAP12B, SI:DKEY-108K21.10, ZGC:110425, SI:CH211-113A14.18, MKI67, SI:DKEY-261M9.12, ZGC:153405, SI:CH211-113A14.12, EIF3S10, BX324216.1 
##     SI:CH211-113A14.24, NUCKS1A, SI:CH211-106E7.2, HNRNPUB, SI:DKEY-108K21.21, CENPF, ZNF638, CSDE1, ASPM, SI:DKEY-108K21.14 
##     SI:CH211-209J10.5, CEP350, SALL4, TPRB, ZGC:153409, SETX, CDH2, CECR2, MARCKSA, NOP14 
## PC_ 5 
## Positive:  FSCN1A, CTH1, DKK1B, GATA6, LHX1A, PITX2, FP102169.1, LFT2, FOXA, FGF8A 
##     VENT, ACTB2, CMTM7, MIXL1, IRX7, EFNB2A, RND1L, SP5A, SOX32, KRT23 
##     IRX3A, PLEKHN1, DDX4, S1PR5A, MSGN1, APLNRB, APLNRA, MESPAB, ID3, ISM1 
## Negative:  SOX3, SOX19A, ID1, FOXD5, CITED4B, CABZ01070258.1, SOX2, ZIC2B, CXCR4B, PFKFB4B 
##     FOXD3, ASB11, CRABP2B, APELA, SERPINH1B, MYCH, HIRIP3, NRARPA, ID2A, NOTO 
##     ADD3B, ACIN1A, KIF4, ALCAMB, CX43.4, PPIG, ARRDC3A, NAV2B, DLA, HSPE1
```

```
VizDimLoadings(SOX3250C, dims = 1:2, reduction = "pca")
```

```
DimPlot(SOX3250C, reduction = "pca")
```

```
ElbowPlot(SOX3250C)
```

```
SOX3250C<- FindNeighbors(SOX3250C, dims = 1:15)
```

```
## Computing nearest neighbor graph
```

```
## Computing SNN
```

```
SOX3250C <- FindClusters(SOX3250C, resolution = 0.1)
```

```
## Modularity Optimizer version 1.3.0 by Ludo Waltman and Nees Jan van Eck
## 
## Number of nodes: 461
## Number of edges: 16281
## 
## Running Louvain algorithm...
## Maximum modularity in 10 random starts: 0.9000
## Number of communities: 1
## Elapsed time: 0 seconds
```

```
SOX3250C <- RunUMAP(SOX3250C, dims = 1:15)
```

```
## 13:31:52 UMAP embedding parameters a = 0.9922 b = 1.112
```

```
## 13:31:52 Read 461 rows and found 15 numeric columns
```

```
## 13:31:52 Using Annoy for neighbor search, n_neighbors = 30
```

```
## 13:31:52 Building Annoy index with metric = cosine, n_trees = 50
```

```
## 0%   10   20   30   40   50   60   70   80   90   100%
```

```
## [----|----|----|----|----|----|----|----|----|----|
```

```
## **************************************************|
## 13:31:52 Writing NN index file to temp file /var/folders/s2/phwgmcs926x8kbjk57m5lqw14_t3jy/T//RtmpquxPVo/file15ca569c40507
## 13:31:52 Searching Annoy index using 1 thread, search_k = 3000
## 13:31:53 Annoy recall = 100%
## 13:31:53 Commencing smooth kNN distance calibration using 1 thread
## 13:31:53 Initializing from normalized Laplacian + noise
## 13:31:53 Commencing optimization for 500 epochs, with 16050 positive edges
## 13:31:54 Optimization finished
```

```
DimPlot(SOX3250C, reduction = "umap")
```

```
# Visualizing expression of sox32 in gata5 positive cells

FeaturePlot(SOX3250C, features = c("SOX32"))
```

```
# Cell cycle analysis: A list of cell cycle markers, from Tirosh et al, 2015, is loaded with Seurat. We can segregate this list into markers of G2/M phase and markers of S phase

s.genes <- cc.genes$s.genes
g2m.genes <- cc.genes$g2m.genes

# We assign scores with the CellCycleScoring function on our gata5 positive cells. This stores S and G2/M scores in object meta data, along with the predicted classification of each cell in either G2M, S or G1 phase

SOX3250C <- CellCycleScoring(SOX3250C, s.features = s.genes, g2m.features = g2m.genes, set.ident = TRUE)
```

```
## Warning: The following features are not present in the object: UNG, CDCA7,
## MLF1IP, CLSPN, not searching for symbol synonyms
```

```
## Warning: The following features are not present in the object: HMGB2, BIRC5,
## TMPO, FAM64A, KIF20B, HJURP, CDCA3, HN1, CDC25C, RANGAP1, PSRC1, CENPA, not
## searching for symbol synonyms
```

```
# Running a PCA on cell cycle genes 

SOX3250C <- RunPCA(SOX3250C, features = c(s.genes, g2m.genes))
```

```
## Warning in PrepDR(object = object, features = features, verbose = verbose): The
## following 16 features requested have not been scaled (running reduction without
## them): UNG, CDCA7, MLF1IP, CLSPN, HMGB2, BIRC5, TMPO, FAM64A, KIF20B, HJURP,
## CDCA3, HN1, CDC25C, RANGAP1, PSRC1, CENPA
```

```
## Warning in irlba(A = t(x = object), nv = npcs, ...): You're computing too large
## a percentage of total singular values, use a standard svd instead.
```

```
## PC_ 1 
## Positive:  TOP2A, CDK1, ANP32E, NUSAP1, MKI67, LBR, CENPF, RAD51, AURKB, TACC3 
##     HMMR, UBE2C, CTCF, KIF2C, NUF2, TPX2, G2E3, DTL, AURKA, GMNN 
##     ANLN, KIF11, UHRF1, DLGAP5, NDC80, CCNE2, SMC4, CBX5, E2F8, CKAP2L 
## Negative:  EXO1, PCNA, CDC6, NEK2, CDC20, TIPIN, BUB1, CKS1B, ATAD2, CKAP5 
##     TYMS, CCNB2, MCM5, GINS2, MCM2, MCM4, MSH2, RPA2, CKS2, GAS2L3 
##     USP1, BRIP1, POLD3, CDC45, UBR7, RFC2, BLM, RAD51AP1, WDR76, GTSE1 
## PC_ 2 
## Positive:  PCNA, CDC6, SLBP, MCM6, PRIM1, CCNB2, CCNE2, RFC2, CDC20, UBE2C 
##     MCM5, MCM4, ANP32E, DTL, UBR7, MCM2, RRM1, GMNN, AURKA, NUSAP1 
##     RPA2, NCAPD2, RAD51AP1, CHAF1B, CDCA8, FEN1, CKS1B, HELLS, LBR, NDC80 
## Negative:  MKI67, RRM2, CTCF, CENPF, CENPE, DLGAP5, HMMR, NUF2, ATAD2, E2F8 
##     SMC4, CASP8AP2, BUB1, TPX2, POLD3, ECT2, KIF2C, TACC3, GAS2L3, CKAP2L 
##     NASP, TOP2A, CBX5, CKAP2, CKAP5, CDK1, CKS2, G2E3, UHRF1, MSH2 
## PC_ 3 
## Positive:  SMC4, RRM2, CDCA2, CDC6, TYMS, CASP8AP2, USP1, PCNA, POLD3, FEN1 
##     CTCF, MCM4, NASP, GMNN, DLGAP5, UHRF1, POLA1, HELLS, CCNB2, CHAF1B 
##     MKI67, CENPF, WDR76, CDC20, MSH2, TIPIN, TTK, RRM1, MCM5, PRIM1 
## Negative:  UBE2C, KIF11, AURKB, NUSAP1, TOP2A, AURKA, G2E3, TACC3, GINS2, TPX2 
##     NDC80, CDK1, CKS1B, KIF23, RAD51AP1, CKS2, BRIP1, HMMR, NEK2, NUF2 
##     CENPE, RAD51, E2F8, TUBB4B, NCAPD2, BLM, CKAP2L, RPA2, CDC45, MCM6 
## PC_ 4 
## Positive:  GMNN, GTSE1, NDC80, MSH2, TIPIN, TPX2, POLD3, DLGAP5, UBR7, LBR 
##     ATAD2, DTL, UBE2C, EXO1, KIF2C, GINS2, CCNE2, USP1, KIF23, ANLN 
##     RAD51AP1, RRM1, CCNB2, NUSAP1, PRIM1, CDC20, HELLS, CDCA2, AURKA, KIF11 
## Negative:  MCM2, BRIP1, RFC2, RPA2, MCM5, MCM6, CKAP2, NEK2, TOP2A, HMMR 
##     CKS1B, CDC45, CKAP5, CHAF1B, RRM2, DSCC1, SLBP, BUB1, NUF2, SMC4 
##     NCAPD2, ECT2, UHRF1, CENPF, FEN1, CENPE, CTCF, MCM4, TUBB4B, CKS2 
## PC_ 5 
## Positive:  HELLS, RFC2, LBR, CDCA8, CKS1B, ATAD2, KIF2C, BRIP1, NASP, CBX5 
##     TTK, DLGAP5, RAD51, TPX2, DSCC1, POLA1, RAD51AP1, CENPF, CCNB2, NUSAP1 
##     USP1, WDR76, MKI67, CDC20, CTCF, MCM2, RRM1, CKAP2, CENPE, TUBB4B 
## Negative:  TACC3, TIPIN, HMMR, BLM, SLBP, SMC4, CDCA2, ECT2, TYMS, EXO1 
##     CKAP2L, RPA2, CDC6, POLD3, AURKB, CDC45, CKS2, KIF11, UBR7, NEK2 
##     NUF2, RRM2, G2E3, DTL, GTSE1, PRIM1, CKAP5, NDC80, ANLN, GAS2L3
```

```
VizDimLoadings(SOX3250C, dims = 1:2, reduction = "pca")
```

```
DimPlot(SOX3250C, reduction = "pca", dims = 1:2, cols = c('S' = '#00A9FF', 'G2M' = 'green', 'G1' = 'magenta'))
```

```
# We visualize sox32 expression across gata5 positive cells in different phases of the cell cycle 

FeaturePlot(SOX3250C, features = c("SOX32"),
        label = TRUE,
        split.by = "Phase")  + NoLegend()
```

```
# Isolating only sox32 positive cells for downsrteam analysis

SOX32cycle <- subset(SOX3250C, subset = SOX32 > 0, slot = "counts" )

SOX32cycle
```

```
## An object of class Seurat 
## 17239 features across 145 samples within 1 assay 
## Active assay: RNA (17239 features, 2000 variable features)
##  3 dimensional reductions calculated: pca, tsne, umap
```

```
# We visualize the distribution of sox32 positivecells across the cell cycle phases

as_tibble(SOX32cycle[[]]) %>%
  ggplot(aes(x=S.Score, y=G2M.Score, color=Phase)) + 
  geom_point() +
  coord_cartesian(xlim=c(-0.80,0.80), ylim=c(-0.80,0.80))
```

```
# We visualize wether the expression levels of sox32 and other cell cycle markers correlate with a specific phase of the cell cycle

VlnPlot(SOX32cycle, features = c("SOX32", "TOP2A", "MKI67", "PCNA","CDC6" ), slot = "counts", log = TRUE)
```

```
# We extract the number of sox32 positive cells in each phase of the cell cycle

G1<-subset(x = SOX32cycle, idents = "G1")
G2M<-subset(x = SOX32cycle, idents = "G2M")
S<-subset(x = SOX32cycle, idents = "S")

G1
```

```
## An object of class Seurat 
## 17239 features across 36 samples within 1 assay 
## Active assay: RNA (17239 features, 2000 variable features)
##  3 dimensional reductions calculated: pca, tsne, umap
```

```
G2M
```

```
## An object of class Seurat 
## 17239 features across 78 samples within 1 assay 
## Active assay: RNA (17239 features, 2000 variable features)
##  3 dimensional reductions calculated: pca, tsne, umap
```

```
S
```

```
## An object of class Seurat 
## 17239 features across 31 samples within 1 assay 
## Active assay: RNA (17239 features, 2000 variable features)
##  3 dimensional reductions calculated: pca, tsne, umap
```

```
# We use the Fetch data function to extract raw counts of the genes listed above for each cell, this for subsequent plotting using Prism

df<-FetchData(G1, vars = "SOX32",slot = "counts")
df1<-FetchData(G2M, vars = "SOX32",slot = "counts")
df2<-FetchData(S, vars = "SOX32",slot = "counts")
```

```
write.infile(df, file="/Users/gugliel/Desktop/SOX32sox32positive G1.csv", sep = ";")
write.infile(df1, file="/Users/gugliel/Desktop/SOX32sox32positive G2M.csv", sep = ";")
write.infile(df2, file="/Users/gugliel/Desktop/SOX32sox32positive S.csv", sep = ";")
```

```
dfTOP2<-FetchData(G1, vars = "TOP2A",slot = "counts")
df1TOP2<-FetchData(G2M, vars = "TOP2A",slot = "counts")
df2TOP2<-FetchData(S, vars = "TOP2A",slot = "counts")
```

```
write.infile(dfTOP2, file="/Users/gugliel/Desktop/TOP2sox32positive G1.csv", sep = ";")
write.infile(df1TOP2, file="/Users/gugliel/Desktop/TOP2sox32positive G2M.csv", sep = ";")
write.infile(df2TOP2, file="/Users/gugliel/Desktop/TOP2sox32positive S.csv", sep = ";")
```

```
dfMKI67<-FetchData(G1, vars = "MKI67",slot = "counts")
df1MKI67<-FetchData(G2M, vars = "MKI67",slot = "counts")
df2MKI67<-FetchData(S, vars = "MKI67",slot = "counts")
```

```
write.infile(dfMKI67, file="/Users/gugliel/Desktop/MKI67sox32positive G1.csv", sep = ";")
write.infile(df1MKI67, file="/Users/gugliel/Desktop/MKI67sox32positive G2M.csv", sep = ";")
write.infile(df2MKI67, file="/Users/gugliel/Desktop/MKI67sox32positive S.csv", sep = ";")
```

```
dfMCM6<-FetchData(G1, vars = "MCM6",slot = "counts")
df1MCM6<-FetchData(G2M, vars = "MCM6",slot = "counts")
df2MCM6<-FetchData(S, vars = "MCM6",slot = "counts")
```

```
write.infile(dfMCM6, file="/Users/gugliel/Desktop/MCM6sox32positive G1.csv", sep = ";")
write.infile(df1MCM6, file="/Users/gugliel/Desktop/MCM6sox32positive G2M.csv", sep = ";")
write.infile(df2MCM6, file="/Users/gugliel/Desktop/MCM6sox32positive S.csv", sep = ";")
```

```
dfPCNA<-FetchData(G1, vars = "PCNA",slot = "counts")
df1PCNA<-FetchData(G2M, vars = "PCNA",slot = "counts")
df2PCNA<-FetchData(S, vars = "PCNA",slot = "counts")
```

```
write.infile(dfPCNA, file="/Users/gugliel/Desktop/PCNAsox32positive G1.csv", sep = ";")
write.infile(df1PCNA, file="/Users/gugliel/Desktop/PCNAsox32positive G2M.csv", sep = ";")
write.infile(df2PCNA, file="/Users/gugliel/Desktop/PCNAsox32positive S.csv", sep = ";")
```

```
dfCDC6<-FetchData(G1, vars = "CDC6",slot = "counts")
df1CDC6<-FetchData(G2M, vars = "CDC6",slot = "counts")
df2CDC6<-FetchData(S, vars = "CDC6",slot = "counts")
```

```
write.infile(dfCDC6, file="/Users/gugliel/Desktop/CDC6sox32positive G1.csv", sep = ";")
write.infile(df1CDC6, file="/Users/gugliel/Desktop/CDC6sox32positive G2M.csv", sep = ";")
write.infile(df2CDC6, file="/Users/gugliel/Desktop/CDC6sox32positive S.csv", sep = ";")
```

```
# As a last control we isolate sox32 negative cells and also check their distribution across the cell cycle phases

SOX32negativecycle <- subset(SOX3250C, subset = SOX32 < 1, slot = "counts" )

SOX32negativecycle
```

```
## An object of class Seurat 
## 17239 features across 316 samples within 1 assay 
## Active assay: RNA (17239 features, 2000 variable features)
##  3 dimensional reductions calculated: pca, tsne, umap
```

```
G1negative<-subset(x = SOX32negativecycle, idents = "G1")
G2Mnegative<-subset(x = SOX32negativecycle, idents = "G2M")
Snegative<-subset(x = SOX32negativecycle, idents = "S")


as_tibble(SOX32negativecycle[[]]) %>%
  ggplot(aes(x=S.Score, y=G2M.Score, color=Phase)) + 
  geom_point() +
  coord_cartesian(xlim=c(-0.80,0.80), ylim=c(-0.80,0.80))
```

```
G1negative
```

```
## An object of class Seurat 
## 17239 features across 51 samples within 1 assay 
## Active assay: RNA (17239 features, 2000 variable features)
##  3 dimensional reductions calculated: pca, tsne, umap
```

```
G2Mnegative
```

```
## An object of class Seurat 
## 17239 features across 168 samples within 1 assay 
## Active assay: RNA (17239 features, 2000 variable features)
##  3 dimensional reductions calculated: pca, tsne, umap
```

```
Snegative
```

```
## An object of class Seurat 
## 17239 features across 97 samples within 1 assay 
## Active assay: RNA (17239 features, 2000 variable features)
##  3 dimensional reductions calculated: pca, tsne, umap
```
