## Supplementary File 4 for "Nodal signaling establishes a competency window for stochastic cell fate switching": Supplementary file 4.html

Endoderm analysis 60% epiboly (gata5 positive cells)


### Endoderm analysis 60% epiboly (gata5 positive cells)

###### Luca Guglielmi

#### 14/01/2021

```
# Loading required packages

library( Seurat )
```

```
## Attaching SeuratObject
```

```
library( ggplot2 )
library( dplyr )
```

```
urd_srat_obj2 <- zf60_srat_obj  
ElbowPlot(urd_srat_obj2)
```

```
urd_srat_obj2<-FindNeighbors(urd_srat_obj2, dim = 1:20)
```

```
## Computing nearest neighbor graph
```

```
## Computing SNN
```

```
urd_srat_obj2<-FindClusters(urd_srat_obj2, resolution = 0.1)
```

```
## Modularity Optimizer version 1.3.0 by Ludo Waltman and Nees Jan van Eck
## 
## Number of nodes: 4101
## Number of edges: 140204
## 
## Running Louvain algorithm...
## Maximum modularity in 10 random starts: 0.9385
## Number of communities: 3
## Elapsed time: 0 seconds
```

```
urd_srat_obj2<-RunUMAP(urd_srat_obj2, dim = 1:20 )
```

```
## Warning: The default method for RunUMAP has changed from calling Python UMAP via reticulate to the R-native UWOT using the cosine metric
## To use Python UMAP via reticulate, set umap.method to 'umap-learn' and metric to 'correlation'
## This message will be shown once per session
```

```
## 13:45:02 UMAP embedding parameters a = 0.9922 b = 1.112
```

```
## 13:45:02 Read 4101 rows and found 20 numeric columns
```

```
## 13:45:02 Using Annoy for neighbor search, n_neighbors = 30
```

```
## 13:45:02 Building Annoy index with metric = cosine, n_trees = 50
```

```
## 0%   10   20   30   40   50   60   70   80   90   100%
```

```
## [----|----|----|----|----|----|----|----|----|----|
```

```
## **************************************************|
## 13:45:02 Writing NN index file to temp file /var/folders/s2/phwgmcs926x8kbjk57m5lqw14_t3jy/T//RtmpapQvMC/file15e2634f1f9b6
## 13:45:02 Searching Annoy index using 1 thread, search_k = 3000
## 13:45:03 Annoy recall = 100%
## 13:45:03 Commencing smooth kNN distance calibration using 1 thread
## 13:45:03 Initializing from normalized Laplacian + noise
## 13:45:04 Commencing optimization for 500 epochs, with 160084 positive edges
## 13:45:08 Optimization finished
```

```
DimPlot (urd_srat_obj2, reduction = "umap", label = TRUE,  pt.size = 1) + NoLegend()
```

```
# Expression of mesoderm, endoderm and ectoderm markers at 60% epiboly

FeaturePlot(urd_srat_obj2, features = c("SOX32"), pt.size = 1.5)
```

```
FeaturePlot(urd_srat_obj2, features = c("GATA5"), pt.size = 1.5)
```

```
FeaturePlot(urd_srat_obj2, features = c("SOX2"), pt.size = 1.5)
```

```
FeaturePlot(urd_srat_obj2, features = c("TA"), pt.size = 1.5)
```

```
# Sub-setting gata5 positive cells into a new Seurat object

SOX3260 <- subset(urd_srat_obj2, subset = GATA5 > 0, slot = "counts" )
SOX3260
```

```
## An object of class Seurat 
## 17239 features across 549 samples within 1 assay 
## Active assay: RNA (17239 features, 2000 variable features)
##  3 dimensional reductions calculated: pca, tsne, umap
```

```
# Re-processing of gata5 positive cells and PCA/uMAP computing

SOX3260<- NormalizeData(SOX3260, normalization.method = "LogNormalize", scale.factor = 10000)
SOX3260<- FindVariableFeatures(SOX3260, selection.method = "vst", nfeatures = 2000)
top10 <- head(VariableFeatures(SOX3260), 10)
plot1 <- VariableFeaturePlot(SOX3260)
plot1 <- LabelPoints(plot = plot1, points = top10, repel = TRUE)
```

```
## When using repel, set xnudge and ynudge to 0 for optimal results
```

```
plot1
```

```
## Warning: Transformation introduced infinite values in continuous x-axis
```

```
## Warning: Removed 2882 rows containing missing values (geom_point).
```

```
all.genes <- rownames(SOX3260)
SOX3260 <- ScaleData(SOX3260, features = all.genes)
```

```
## Centering and scaling data matrix
```

```
SOX3260 <- RunPCA(SOX3260, features = VariableFeatures(object = SOX3260))
```

```
## PC_ 1 
## Positive:  VED, HES6, MSGN1, CDX4, MESPAB, ID1, RBM38, ALDH1A2, CXCL12A, IM:7138239 
##     SI:DKEY-261J4.5, WNT8A, EVE1, ANP32E, VOX, WNT11, SIX4A, BAMBIA, TA, HSPB1 
##     SI:CH73-1A9.3, SP5L, CRABP2B, TBX6L, HER7, SI:CH211-222L21.1, MESPAA, SNAI1A, SKILB, CYP27C1 
## Negative:  GSC, FRZB, NOG1, FZD8B, FZD8A, KLF17, SHISA2, LFT2, RRBP1B, CHD 
##     OTX1B, SIX7, SIX3B, FOXA3, DHRS3B, OTX1A, ELL2, XBP1, CITED4B, ID3 
##     ZGC:174153, SI:CH211-163L21.7, B3GNT5B, BTG2, TMEM120A, RDH8A, ZGC:174855, PKDCCA, KRT18, GADD45BA 
## PC_ 2 
## Positive:  APLNRB, OSR1, APLNRA, MSGN1, MYCH, EFNB2B, FOXA, ALDH1A2, CYP27C1, RIPPLY1 
##     LFT1, P4HA2, RBM38, ZIC3, FOXC1A, PPRC1, PCDH8, KLF17, DACT2, FZD8B 
##     DDIT4, MESPAA, XBP1, SIX3B, FGF17, CXCL12B, CITED4B, FRZB, RRBP1B, RDH10A 
## Negative:  SOX32, CXCR4A, SOX17, ACKR3B, CABZ01070258.1, CDH6, PRDX5, SI:CH73-234B20.5, FOXA2, PLPP3 
##     ARL5C, GPM6AB, GSTT1B, SI:CH211-152C2.3, GATSL2, MARCKSL1B, LMO4A, SSUH2RS1, FLRT3, IQCA1 
##     VGLL4L, S1PR5A, CA9, FKBP7, ST6GALNAC, GPD1B, CYP2AA8, DNMT3BB.1, DHRS3B, TUBB5 
## PC_ 3 
## Positive:  MALAT1, HMGN2, MDKB, TDH, SOX19A, C7ORF50, HISTH1L, PPAN, P4HA1B, SI:DKEY-85K7.7 
##     TFAP2A, PRDM1A, APELA, APOC1, CRABP2B, SOX3, SOX2, EVA1BA, NRARPA, DLX3B 
##     TUBA1A, APOEB, MSX1B, PRR12B, ATP1B3A, ZGC:113208, NAT8L, ALCAMB, ACTB1, ZGC:174153 
## Negative:  RRM2, FSCN1A, ELOVL6, ISM1, SNAI1A, EFNB2A, LHX1A, RND1L, TBX16, KIRREL3L 
##     BX324216.1, HNRNPUB, FOXA, GID8B, PITX2, ZGC:153409, AKAP12B, DLEU7, EFNB2B, MARCKSL1B 
##     NCL, CTH1, HIST2H2AB, MIXL1, STM, XBP1, DDX4, APLNRA, FLRT3, FOXA3 
## PC_ 4 
## Positive:  TBX16, TA, BMP4, CDX4, BAMBIA, HES6, SI:DKEY-261J4.5, PPRC1, EVE1, VED 
##     FP102169.1, NUSAP1, RAB33A, DDIT4, BX324216.1, LFT1, TCF3B, SNAI1A, SI:CH211-163L21.8, SI:DKEY-261J4.3 
##     TOP2A, FLRT3, UBTF, EFNB2A, MKI67, TMEM120A, HER7, ICN, NANOG, HELB 
## Negative:  IRX1B, TWIST1A, IRX7, RIPPLY1, EVA1BA, PKDCCB, SFRP1A, APLNRA, SI:CH211-222L21.1, HMGN2 
##     INSB, HER11, EGLN3, SI:CH73-281N10.2, SHISA2, SI:CH211-51C14.1, HER5, H1FX, GNAIA, NTD5 
##     MCL1B, CYP26A1, ZGC:158343, KAZALD2, ZBTB18, HSPB1, MEIS1B, TBX6, ZIC2B, ZGC:65851 
## PC_ 5 
## Positive:  SOX19A, SOX3, SOX2, APELA, SI:CH1073-80I24.3, ALCAMB, CXCR4B, MDKB, TFAP2A, PRDM14 
##     ALDOB, HER3, SI:CH211-170D8.2, TFAP2C, SI:DKEY-27I16.2, PFKFB4B, MSX1B, HER8A, POLR3GLA, THY1 
##     NOTO, KLF2B, ADMP, CCND2A, SI:CH211-152C2.3, FOXB1A, FOXD5, SOX11A, H1M, EPCAM 
## Negative:  ASPH, CXCL12A, TUBA8L2, FSCN1A, APOC1, NID2A, PCDH10B, MEIS1B, TWIST1A, WU:FB55G09 
##     CXCL12B, ARL4AA, APOEB, APLNRB, ATP1B3A, APLNRA, CPN1, ACTB1, RDH10A, IM:7138239 
##     PHLDB1B, ZIC2A, CXXC5B, SI:DKEY-261J4.5, CDH2, EPHB3A, SIX4A, NUSAP1, RIPPLY1, PTTG1IPB
```

```
DimPlot(SOX3260, reduction = "pca")
```

```
VizDimLoadings(SOX3260, dims = 1:2, reduction = "pca")
```

```
ElbowPlot(SOX3260)
```

```
SOX3260<- FindNeighbors(SOX3260, dims = 1:15)
```

```
## Computing nearest neighbor graph
```

```
## Computing SNN
```

```
SOX3260 <- FindClusters(SOX3260, resolution = 0.5)
```

```
## Modularity Optimizer version 1.3.0 by Ludo Waltman and Nees Jan van Eck
## 
## Number of nodes: 549
## Number of edges: 18426
## 
## Running Louvain algorithm...
## Maximum modularity in 10 random starts: 0.7735
## Number of communities: 5
## Elapsed time: 0 seconds
```

```
SOX3260 <- RunUMAP(SOX3260, dims = 1:15)
```

```
## 13:45:16 UMAP embedding parameters a = 0.9922 b = 1.112
```

```
## 13:45:16 Read 549 rows and found 15 numeric columns
```

```
## 13:45:16 Using Annoy for neighbor search, n_neighbors = 30
```

```
## 13:45:16 Building Annoy index with metric = cosine, n_trees = 50
```

```
## 0%   10   20   30   40   50   60   70   80   90   100%
```

```
## [----|----|----|----|----|----|----|----|----|----|
```

```
## **************************************************|
## 13:45:16 Writing NN index file to temp file /var/folders/s2/phwgmcs926x8kbjk57m5lqw14_t3jy/T//RtmpapQvMC/file15e26319c7965
## 13:45:16 Searching Annoy index using 1 thread, search_k = 3000
## 13:45:16 Annoy recall = 100%
## 13:45:16 Commencing smooth kNN distance calibration using 1 thread
## 13:45:17 Initializing from normalized Laplacian + noise
## 13:45:17 Commencing optimization for 500 epochs, with 19834 positive edges
## 13:45:18 Optimization finished
```

```
# Expression of mesoderm, endoderm and ectoderm markers in gata5 positive cells

FeaturePlot(SOX3260, features = c("SOX32"), pt.size = 2)
```

```
FeaturePlot(SOX3260, features = c("GATA5"), pt.size = 2)
```

```
FeaturePlot(SOX3260, features = c("TA"), pt.size = 2)
```

```
FeaturePlot(SOX3260, features = c("SOX2"), pt.size = 2)
```

```
# clustering of gata5 positive cells with increasing granularity

SOX3260<- FindNeighbors(SOX3260, dims = 1:15)
```

```
## Computing nearest neighbor graph
```

```
## Computing SNN
```

```
SOX3260 <- FindClusters(SOX3260, resolution = 0.01)
```

```
## Modularity Optimizer version 1.3.0 by Ludo Waltman and Nees Jan van Eck
## 
## Number of nodes: 549
## Number of edges: 18426
## 
## Running Louvain algorithm...
## Maximum modularity in 10 random starts: 0.9900
## Number of communities: 1
## Elapsed time: 0 seconds
```

```
SOX3260 <- RunUMAP(SOX3260, dims = 1:15)
```

```
## 13:45:19 UMAP embedding parameters a = 0.9922 b = 1.112
```

```
## 13:45:19 Read 549 rows and found 15 numeric columns
```

```
## 13:45:19 Using Annoy for neighbor search, n_neighbors = 30
```

```
## 13:45:19 Building Annoy index with metric = cosine, n_trees = 50
```

```
## 0%   10   20   30   40   50   60   70   80   90   100%
```

```
## [----|----|----|----|----|----|----|----|----|----|
```

```
## **************************************************|
## 13:45:19 Writing NN index file to temp file /var/folders/s2/phwgmcs926x8kbjk57m5lqw14_t3jy/T//RtmpapQvMC/file15e267be24e5c
## 13:45:19 Searching Annoy index using 1 thread, search_k = 3000
## 13:45:19 Annoy recall = 100%
## 13:45:19 Commencing smooth kNN distance calibration using 1 thread
## 13:45:20 Initializing from normalized Laplacian + noise
## 13:45:20 Commencing optimization for 500 epochs, with 19834 positive edges
## 13:45:20 Optimization finished
```

```
DimPlot(SOX3260, reduction = "umap",label = TRUE,  pt.size = 2) + NoLegend()
```

```
SOX3260<- FindNeighbors(SOX3260, dims = 1:15)
```

```
## Computing nearest neighbor graph
```

```
## Computing SNN
```

```
SOX3260 <- FindClusters(SOX3260, resolution = 0.05)
```

```
## Modularity Optimizer version 1.3.0 by Ludo Waltman and Nees Jan van Eck
## 
## Number of nodes: 549
## Number of edges: 18426
## 
## Running Louvain algorithm...
## Maximum modularity in 10 random starts: 0.9562
## Number of communities: 2
## Elapsed time: 0 seconds
```

```
SOX3260 <- RunUMAP(SOX3260, dims = 1:15)
```

```
## 13:45:21 UMAP embedding parameters a = 0.9922 b = 1.112
```

```
## 13:45:21 Read 549 rows and found 15 numeric columns
```

```
## 13:45:21 Using Annoy for neighbor search, n_neighbors = 30
```

```
## 13:45:21 Building Annoy index with metric = cosine, n_trees = 50
```

```
## 0%   10   20   30   40   50   60   70   80   90   100%
```

```
## [----|----|----|----|----|----|----|----|----|----|
```

```
## **************************************************|
## 13:45:21 Writing NN index file to temp file /var/folders/s2/phwgmcs926x8kbjk57m5lqw14_t3jy/T//RtmpapQvMC/file15e2637003dce
## 13:45:21 Searching Annoy index using 1 thread, search_k = 3000
## 13:45:21 Annoy recall = 100%
## 13:45:21 Commencing smooth kNN distance calibration using 1 thread
## 13:45:22 Initializing from normalized Laplacian + noise
## 13:45:22 Commencing optimization for 500 epochs, with 19834 positive edges
## 13:45:22 Optimization finished
```

```
DimPlot(SOX3260, reduction = "umap",label = TRUE,  pt.size = 2) + NoLegend()
```

```
SOX3260<- FindNeighbors(SOX3260, dims = 1:15)
```

```
## Computing nearest neighbor graph
```

```
## Computing SNN
```

```
SOX3260 <- FindClusters(SOX3260, resolution = 0.1)
```

```
## Modularity Optimizer version 1.3.0 by Ludo Waltman and Nees Jan van Eck
## 
## Number of nodes: 549
## Number of edges: 18426
## 
## Running Louvain algorithm...
## Maximum modularity in 10 random starts: 0.9341
## Number of communities: 3
## Elapsed time: 0 seconds
```

```
SOX3260 <- RunUMAP(SOX3260, dims = 1:15)
```

```
## 13:45:23 UMAP embedding parameters a = 0.9922 b = 1.112
```

```
## 13:45:23 Read 549 rows and found 15 numeric columns
```

```
## 13:45:23 Using Annoy for neighbor search, n_neighbors = 30
```

```
## 13:45:23 Building Annoy index with metric = cosine, n_trees = 50
```

```
## 0%   10   20   30   40   50   60   70   80   90   100%
```

```
## [----|----|----|----|----|----|----|----|----|----|
```

```
## **************************************************|
## 13:45:23 Writing NN index file to temp file /var/folders/s2/phwgmcs926x8kbjk57m5lqw14_t3jy/T//RtmpapQvMC/file15e2666aaf062
## 13:45:23 Searching Annoy index using 1 thread, search_k = 3000
## 13:45:23 Annoy recall = 100%
## 13:45:23 Commencing smooth kNN distance calibration using 1 thread
## 13:45:24 Initializing from normalized Laplacian + noise
## 13:45:24 Commencing optimization for 500 epochs, with 19834 positive edges
## 13:45:24 Optimization finished
```

```
DimPlot(SOX3260, reduction = "umap",label = TRUE,  pt.size = 2) + NoLegend()
```

```
SOX3260<- FindNeighbors(SOX3260, dims = 1:15)
```

```
## Computing nearest neighbor graph
```

```
## Computing SNN
```

```
SOX3260 <- FindClusters(SOX3260, resolution = 0.2)
```

```
## Modularity Optimizer version 1.3.0 by Ludo Waltman and Nees Jan van Eck
## 
## Number of nodes: 549
## Number of edges: 18426
## 
## Running Louvain algorithm...
## Maximum modularity in 10 random starts: 0.8935
## Number of communities: 4
## Elapsed time: 0 seconds
```

```
SOX3260 <- RunUMAP(SOX3260, dims = 1:15)
```

```
## 13:45:25 UMAP embedding parameters a = 0.9922 b = 1.112
```

```
## 13:45:25 Read 549 rows and found 15 numeric columns
```

```
## 13:45:25 Using Annoy for neighbor search, n_neighbors = 30
```

```
## 13:45:25 Building Annoy index with metric = cosine, n_trees = 50
```

```
## 0%   10   20   30   40   50   60   70   80   90   100%
```

```
## [----|----|----|----|----|----|----|----|----|----|
```

```
## **************************************************|
## 13:45:25 Writing NN index file to temp file /var/folders/s2/phwgmcs926x8kbjk57m5lqw14_t3jy/T//RtmpapQvMC/file15e26cfb35f
## 13:45:25 Searching Annoy index using 1 thread, search_k = 3000
## 13:45:25 Annoy recall = 100%
## 13:45:25 Commencing smooth kNN distance calibration using 1 thread
## 13:45:26 Initializing from normalized Laplacian + noise
## 13:45:26 Commencing optimization for 500 epochs, with 19834 positive edges
## 13:45:26 Optimization finished
```

```
DimPlot(SOX3260, reduction = "umap",label = TRUE,  pt.size = 2) + NoLegend()
```

```
SOX3260<- FindNeighbors(SOX3260, dims = 1:15)
```

```
## Computing nearest neighbor graph
```

```
## Computing SNN
```

```
SOX3260 <- FindClusters(SOX3260, resolution = 0.5)
```

```
## Modularity Optimizer version 1.3.0 by Ludo Waltman and Nees Jan van Eck
## 
## Number of nodes: 549
## Number of edges: 18426
## 
## Running Louvain algorithm...
## Maximum modularity in 10 random starts: 0.7735
## Number of communities: 5
## Elapsed time: 0 seconds
```

```
SOX3260 <- RunUMAP(SOX3260, dims = 1:15)
```

```
## 13:45:27 UMAP embedding parameters a = 0.9922 b = 1.112
```

```
## 13:45:27 Read 549 rows and found 15 numeric columns
```

```
## 13:45:27 Using Annoy for neighbor search, n_neighbors = 30
```

```
## 13:45:27 Building Annoy index with metric = cosine, n_trees = 50
```

```
## 0%   10   20   30   40   50   60   70   80   90   100%
```

```
## [----|----|----|----|----|----|----|----|----|----|
```

```
## **************************************************|
## 13:45:27 Writing NN index file to temp file /var/folders/s2/phwgmcs926x8kbjk57m5lqw14_t3jy/T//RtmpapQvMC/file15e2633b0b47a
## 13:45:27 Searching Annoy index using 1 thread, search_k = 3000
## 13:45:27 Annoy recall = 100%
## 13:45:27 Commencing smooth kNN distance calibration using 1 thread
## 13:45:27 Initializing from normalized Laplacian + noise
## 13:45:27 Commencing optimization for 500 epochs, with 19834 positive edges
## 13:45:28 Optimization finished
```

```
DimPlot(SOX3260, reduction = "umap",label = TRUE,  pt.size = 2) + NoLegend()
```

```
# sox32 expression within the different clusters


VlnPlot(SOX3260, features = c("SOX32"), slot = "counts", log = TRUE)
```

### Comparing sox32 positive/ negative cells (Phil)

```
#Comparing sox32 positive/ negative cells 

SOX3260 <- AddMetaData( object = SOX3260,
                       metadata = GetAssayData( SOX3260, slot = "counts" )[ "SOX32", ] > 0,
                       col.name = "SOX32_positive" )
SOX3260 <- SetIdent( object = SOX3260, value = "SOX32_positive" )
SOX32.markers <- FindMarkers( SOX3260, ident.2 = "FALSE", ident.1 = "TRUE", min.diff.pct = 0.1, only.pos = TRUE )

head(SOX32.markers,n = 80)
```

```
##                           p_val avg_log2FC pct.1 pct.2     p_val_adj
## SOX32             6.513054e-116  4.8459274 1.000 0.000 1.122785e-111
## SOX17              1.406482e-98  2.8313221 0.882 0.000  2.424634e-94
## CXCR4A             9.783008e-98  4.0878681 0.953 0.069  1.686493e-93
## ACKR3B             2.871377e-92  2.9401648 0.882 0.032  4.949967e-88
## CABZ01070258.1     2.851464e-79  2.7571481 0.888 0.113  4.915639e-75
## FOXA2              4.656408e-62  2.2236806 0.888 0.190  8.027182e-58
## PRDX5              2.736938e-61  2.5662586 0.741 0.084  4.718207e-57
## CDH6               3.856570e-55  2.2263639 0.782 0.158  6.648341e-51
## SI:CH73-234B20.5   1.880308e-48  1.2385184 0.506 0.008  3.241463e-44
## GPM6AB             2.474366e-44  1.4266320 0.659 0.090  4.265560e-40
## PLPP3              1.627414e-43  1.1928515 0.541 0.040  2.805500e-39
## FLRT3              1.344656e-37  1.2277083 0.965 0.678  2.318053e-33
## VGLL4L             2.002368e-37  1.5890223 0.576 0.090  3.451882e-33
## ARL5C              4.661068e-35  1.3506893 0.653 0.161  8.035215e-31
## GATSL2             1.628387e-34  1.3568964 0.682 0.206  2.807176e-30
## DHRS3B             1.198497e-32  1.3417046 0.888 0.369  2.066089e-28
## S1PR5A             2.800919e-32  0.9445497 0.524 0.079  4.828505e-28
## SSUH2RS1           2.280965e-28  0.6203560 0.312 0.005  3.932155e-24
## GSTT1B             3.338181e-28  1.1509928 0.494 0.095  5.754691e-24
## IQCA1              7.798098e-23  0.5310774 0.247 0.003  1.344314e-18
## LMO4A              4.564535e-22  1.1618475 0.624 0.232  7.868802e-18
## FKBP7              8.837972e-22  1.1614178 0.753 0.441  1.523578e-17
## CYP2AA8            4.588112e-21  1.3222393 0.524 0.177  7.909446e-17
## GPD1B              1.548849e-20  0.8342592 0.353 0.055  2.670061e-16
## ST6GALNAC          5.900302e-20  0.5938137 0.271 0.021  1.017153e-15
## LHFP               7.063367e-20  0.7597643 0.335 0.047  1.217654e-15
## CA9                4.190237e-19  0.9620898 0.759 0.435  7.223550e-15
## TMEM223            2.588961e-18  0.6313876 0.359 0.069  4.463110e-14
## FOXD5              5.651336e-18  0.9612686 0.518 0.166  9.742337e-14
## SP5L               7.210786e-17  0.6625336 0.971 0.821  1.243067e-12
## PLTP               1.139902e-15  0.6498533 0.306 0.058  1.965077e-11
## NT5C2B             1.407864e-15  0.7021921 0.229 0.024  2.427017e-11
## MRPL15             1.725145e-15  0.6633827 0.418 0.129  2.973977e-11
## AHI1               2.079765e-15  1.0366174 0.635 0.338  3.585307e-11
## VOX                2.562759e-15  0.8750166 0.888 0.702  4.417940e-11
## TUBB5              2.745328e-15  0.8128632 0.459 0.164  4.732670e-11
## SLC43A2B           3.268524e-15  0.5731481 0.241 0.029  5.634608e-11
## GRAMD3             4.436296e-15  0.8119428 0.747 0.438  7.647730e-11
## SI:CH211-155E24.3  8.102932e-15  0.7341792 0.576 0.266  1.396864e-10
## ARL6IP5A           1.022448e-14  0.7154692 0.341 0.084  1.762598e-10
## SLC25A36B          1.740854e-14  0.5007171 0.318 0.071  3.001058e-10
## NT5C3A             2.629664e-14  0.5053307 0.165 0.005  4.533278e-10
## DNMT3BB.1          2.726120e-14  0.7564956 0.359 0.098  4.699558e-10
## SP5A               3.144625e-14  0.7140936 0.794 0.496  5.421019e-10
## ASPH               5.299719e-14  0.6067080 0.988 0.876  9.136186e-10
## CPN1               1.322199e-13  0.8104985 0.718 0.393  2.279339e-09
## COMMD7             1.518988e-13  0.5848313 0.547 0.227  2.618583e-09
## PHTF1              1.609350e-13  0.7026282 0.312 0.077  2.774359e-09
## SOX13              1.655729e-13  0.6817096 0.294 0.063  2.854311e-09
## ABTB2              3.367073e-13  0.3621937 0.153 0.005  5.804497e-09
## STOX1              3.712814e-13  0.7811559 0.676 0.375  6.400520e-09
## ASB11              4.396631e-13  0.8594048 0.706 0.433  7.579352e-09
## JPH3               9.422410e-13  0.2786527 0.129 0.000  1.624329e-08
## AOC2               9.936803e-13  0.6370061 0.506 0.222  1.713005e-08
## TPD52L2A           1.145072e-12  0.4855320 0.324 0.087  1.973989e-08
## CMTM7              1.929247e-12  0.7548694 0.565 0.282  3.325829e-08
## TSPAN18A           2.246689e-12  0.3206508 0.135 0.003  3.873068e-08
## RBMS2B             2.738799e-12  0.4000984 0.265 0.053  4.721415e-08
## ETV5B              4.385294e-12  0.7132336 0.835 0.662  7.559809e-08
## LIMA1A             5.715481e-12  0.7296123 0.659 0.369  9.852917e-08
## FAM212AB           6.853723e-12  0.6387819 0.824 0.554  1.181513e-07
## LMO4B              7.769858e-12  0.5537667 0.329 0.095  1.339446e-07
## DLA                1.064722e-11  0.6475618 0.353 0.111  1.835474e-07
## PLEKHG5            1.176200e-11  0.4261057 0.353 0.108  2.027652e-07
## CHAC1              1.668042e-11  0.7168634 0.588 0.330  2.875538e-07
## CDH2               2.220711e-11  0.6277011 0.806 0.599  3.828283e-07
## TP53I11A           3.342862e-11  0.4327228 0.253 0.058  5.762760e-07
## TRIP10A            7.039894e-11  0.2686062 0.135 0.008  1.213607e-06
## ANGPTL7            8.145563e-11  0.4331668 0.118 0.003  1.404214e-06
## CCDC39             8.434847e-11  0.2667581 0.118 0.003  1.454083e-06
## SI:CH211-193L2.5   1.862725e-10  0.4280401 0.165 0.021  3.211152e-06
## PHLDB1B            2.023803e-10  0.6096508 0.424 0.185  3.488835e-06
## TSKU               2.152644e-10  0.5721713 0.641 0.367  3.710943e-06
## CXXC5B             2.410171e-10  0.4662821 0.453 0.187  4.154894e-06
## HER5               2.474306e-10  0.8232250 0.306 0.098  4.265456e-06
## SI:DKEY-147F3.4    2.983330e-10  0.2532356 0.129 0.008  5.142962e-06
## SESN1              3.638580e-10  0.3722318 0.218 0.045  6.272548e-06
## ARG2               3.642809e-10  0.4641406 0.229 0.053  6.279839e-06
## ARHGAP6            4.081463e-10  0.2694520 0.135 0.011  7.036034e-06
## AHCYL2             5.528821e-10  0.5990052 0.671 0.454  9.531135e-06
```

```
SOX3260 <- AddMetaData( object = SOX3260,
                       metadata = GetAssayData( SOX3260, slot = "counts" )[ "SOX32", ] < 1,
                       col.name = "SOX32_negative" )
SOX3260 <- SetIdent( object = SOX3260, value = "SOX32_negative" )
SOX32.markers <- FindMarkers( SOX3260, ident.2 = "FALSE", ident.1 = "TRUE", min.diff.pct = 0.1, only.pos = TRUE )

head(SOX32.markers,n = 80)
```

```
##                         p_val avg_log2FC pct.1 pct.2    p_val_adj
## APLNRB           1.224266e-41  1.7767334 0.784 0.182 2.110512e-37
## APLNRA           4.757997e-33  1.5497239 0.847 0.453 8.202312e-29
## MSGN1            1.598879e-28  2.4455272 0.739 0.347 2.756308e-24
## MYCH             1.904875e-26  1.2220393 0.807 0.488 3.283813e-22
## OSR1             6.027242e-25  1.4482966 0.599 0.153 1.039036e-20
## FOXC1A           4.227990e-24  1.2828422 0.507 0.047 7.288633e-20
## CXCL12B          9.334189e-24  1.2988858 0.749 0.365 1.609121e-19
## ZIC3             2.770096e-23  1.2454282 0.628 0.188 4.775368e-19
## ALDH1A2          1.340314e-22  1.5396873 0.578 0.147 2.310568e-18
## CXCL12A          1.472859e-22  1.4119439 0.784 0.500 2.539062e-18
## RBM38            4.892992e-22  1.5451572 0.631 0.229 8.435029e-18
## EFNB2B           7.350157e-20  1.2200323 0.694 0.365 1.267094e-15
## CYP27C1          1.102315e-19  1.1413675 0.736 0.406 1.900281e-15
## SIX4A            1.994424e-18  0.9878079 0.573 0.182 3.438187e-14
## IM:7138239       6.135119e-18  1.2566484 0.639 0.306 1.057633e-13
## WNT8A            3.256225e-17  1.2265746 0.496 0.124 5.613407e-13
## RDH10A           1.097882e-16  1.2206862 0.609 0.271 1.892638e-12
## RIPPLY1          1.199063e-16  2.0541738 0.507 0.159 2.067064e-12
## MESPAA           1.363062e-16  1.3042245 0.393 0.047 2.349782e-12
## DDIT4            1.474147e-15  1.0817145 0.707 0.441 2.541282e-11
## TBX6L            1.670477e-14  0.8313508 0.325 0.024 2.879735e-10
## FOXA             7.996294e-14  1.1149871 0.770 0.565 1.378481e-09
## P4HA2            1.223646e-13  0.9197008 0.472 0.141 2.109443e-09
## SKILB            3.529585e-13  0.7854776 0.388 0.088 6.084652e-09
## SERPINH1B        5.034492e-13  0.9796308 0.480 0.171 8.678960e-09
## ADD3B            7.922635e-13  0.6599541 0.734 0.412 1.365783e-08
## HER1             1.118270e-12  0.9803502 0.356 0.065 1.927785e-08
## PPRC1            2.320000e-12  0.8412495 0.670 0.412 3.999448e-08
## MESPAB           3.125116e-12  1.1521424 0.728 0.559 5.387388e-08
## LFT1             5.334691e-12  0.8307784 0.380 0.088 9.196474e-08
## DACT2            1.139355e-11  0.7701399 0.499 0.200 1.964134e-07
## LPAR1            3.076581e-11  0.7923215 0.472 0.194 5.303719e-07
## PCDH8            3.450050e-11  0.7673720 0.665 0.412 5.947542e-07
## RHOV             6.305255e-11  0.6866819 0.607 0.347 1.086963e-06
## TOB1A            7.937131e-11  0.7287505 0.303 0.053 1.368282e-06
## DLC              1.301244e-10  0.6887729 0.235 0.012 2.243215e-06
## PCDH10B          1.310907e-10  0.6282156 0.325 0.071 2.259873e-06
## ZIC2A            1.727894e-10  0.7234909 0.757 0.600 2.978716e-06
## MIXL1            4.264637e-10  0.7350119 0.504 0.241 7.351808e-06
## CRABP2B          7.069356e-10  0.8858713 0.525 0.265 1.218686e-05
## SURF6            2.315252e-09  0.6096898 0.763 0.612 3.991264e-05
## ATP1B1A          9.173895e-09  0.7044443 0.865 0.753 1.581488e-04
## RIPPLY2          1.098437e-08  0.5105717 0.206 0.018 1.893596e-04
## CITED4A          1.558746e-08  0.4604160 0.264 0.053 2.687123e-04
## PPP1R3CA         1.681206e-08  0.7075230 0.522 0.294 2.898232e-04
## RRS1             1.910752e-08  0.5607978 0.810 0.676 3.293946e-04
## CITED4B          2.479196e-08  1.0463953 0.641 0.418 4.273886e-04
## HER7             2.789029e-08  0.9152930 0.325 0.106 4.808007e-04
## IER5             2.829684e-08  0.4999905 0.826 0.676 4.878093e-04
## TWIST1A          3.138467e-08  0.7562488 0.208 0.024 5.410403e-04
## SI:DKEY-261H17.1 3.735503e-08  0.4350744 0.216 0.029 6.439634e-04
## GAS1B            4.685840e-08  0.6505763 0.404 0.182 8.077920e-04
## HN1L             2.139638e-07  0.6023637 0.675 0.506 3.688522e-03
## SI:DKEY-261J4.5  3.174711e-07  0.7607887 0.631 0.453 5.472885e-03
## DLD              5.617261e-07  0.5458070 0.227 0.053 9.683597e-03
## POLR3GLA         5.640435e-07  0.5737086 0.831 0.724 9.723546e-03
## ZEB2A            6.916118e-07  0.4025915 0.219 0.047 1.192270e-02
## FGF17            7.188919e-07  0.4926567 0.309 0.118 1.239298e-02
## ADMP             7.359257e-07  0.6847775 0.195 0.035 1.268662e-02
## RRP1             8.854034e-07  0.4577300 0.773 0.659 1.526347e-02
## ID2A             1.085255e-06  0.5657790 0.285 0.100 1.870871e-02
## CDKN1CB          1.294049e-06  0.5728804 0.557 0.341 2.230811e-02
## ARL4AA           1.429613e-06  0.5333433 0.216 0.053 2.464509e-02
## MFSD2AB          2.039070e-06  0.3250075 0.169 0.024 3.515152e-02
## LBX2             4.002665e-06  0.3496351 0.140 0.012 6.900194e-02
## FADD             5.329204e-06  0.3509975 0.137 0.012 9.187015e-02
## PPP1R14AB        5.712714e-06  0.3110869 0.137 0.012 9.848148e-02
## PPP1R14C         5.782561e-06  0.4513118 0.193 0.047 9.968556e-02
## TDH              5.837224e-06  0.5885703 0.343 0.159 1.006279e-01
## TUBA1A           7.051483e-06  0.4987958 0.433 0.241 1.215605e-01
## CDX4             7.192840e-06  0.9038595 0.559 0.376 1.239974e-01
## MEIS1B           1.754294e-05  0.3569456 0.248 0.088 3.024227e-01
## FGF8A            1.820447e-05  0.4798517 0.269 0.106 3.138269e-01
## FOXH1            2.263684e-05  0.4317720 0.388 0.206 3.902364e-01
## PCDH1B           2.501704e-05  0.4426169 0.290 0.129 4.312688e-01
## STIM2            2.893703e-05  0.3029218 0.256 0.106 4.988455e-01
## NDUFV3           3.012951e-05  0.4856393 0.588 0.453 5.194026e-01
## SI:CH73-299H12.3 3.049028e-05  0.3546175 0.372 0.194 5.256219e-01
## DKK1B            3.231128e-05  1.0380722 0.383 0.206 5.570141e-01
## SI:DKEY-261J4.4  5.479399e-05  0.5158399 0.335 0.176 9.445935e-01
```

```
FeaturePlot (SOX3260, features = c("SOX32_positive" ), pt.size = 2)
```

```
FeaturePlot (SOX3260, features = c("SOX32_negative" ), pt.size = 2)
```
